# Ligand efficacy shifts a nuclear receptor conformational ensemble between transcriptionally active and repressive states

**DOI:** 10.1101/2024.04.23.590805

**Authors:** Brian MacTavish, Di Zhu, Jinsai Shang, Qianzhen Shao, Zhongyue J. Yang, Theodore M. Kamenecka, Douglas J. Kojetin

## Abstract

Nuclear receptors (NRs) are thought to dynamically alternate between transcriptionally active and repressive conformations, which are stabilized upon ligand binding. Most NR ligand series exhibit limited bias, primarily consisting of transcriptionally active agonists or neutral antagonists, but not repressive inverse agonists—a limitation that restricts understanding of the functional NR conformational ensemble. Here, we report a NR ligand series for peroxisome proliferator-activated receptor gamma (PPARγ) that spans a pharmacological spectrum from repression (inverse agonism) to activation (agonism) where subtle structural modifications switch compound activity. While crystal structures provide snapshots of the fully repressive state, NMR spectroscopy and conformation-activity relationship analysis reveals that compounds within the series shift the PPARγ conformational ensemble between transcriptionally active and repressive conformations that are populated in the apo/ligand-free ensemble. Our findings reveal a molecular framework for minimal chemical modifications that enhance PPARγ inverse agonism and elucidate their influence on the dynamic PPARγ conformational ensemble.

## INTRODUCTION

Nuclear receptors (NRs) are ligand-regulated transcription factors that control gene expression in response to binding endogenous metabolites and synthetic ligands, including about 15% of FDA-approved drugs ^1^. NRs contain a modular domain architecture consisting of an N-terminal disordered activation domain, a central DNA-binding domain, and a C-terminal ligand-binding domain (LBD). A fundamental hypothesis in the NR field posits that in the absence of ligand the LBD exchanges between transcriptionally active and repressive states that can be stabilized upon ligand binding ^2,3^. Binding of an agonist stabilizes an LBD conformation that favors recruitment of coactivator proteins over corepressors, resulting in transcriptional activation of gene expression. Pharmacological antagonists block ligand binding and are generally considered transcriptionally neutral, stabilizing an LBD conformation that does not significantly alter coregulator recruitment compared to the unliganded state. Inverse agonists stabilize an LBD conformation that promotes interaction with corepressor proteins over coactivators, leading to transcriptional repression of gene expression. These ligand-dependent activities, which result in distinct coregulator recruitment and transcriptional outcomes, are known as ligand bias ^4,5^.

Although the field has a relatively good understanding of the structural basis of agonist-induced, coactivator-selective NR functions associated with transcriptional activation, several features remain poorly understood. These include graded transcriptional activity (e.g., partial vs. full agonism), transcriptional repression (inverse agonism), and whether agonists and inverse agonists select natively populated structural conformations (conformational selection) or induce non-native conformations. X-ray crystallography and cryo-electron microscopy have provided structural insights into DNA- and coregulator-bound NR transcriptional complexes, including LBDs bound to different pharmacological ligands ^6,7^. However, these structural methods that capture static snapshots of functionally relevant NR complexes fail to explain features relevant to the dynamic functional LBD conformational ensemble, which are detectable by other methods such as NMR spectroscopy ^8^. Moreover, studies into the mechanisms of graded NR transcriptional repression have been hampered because most NR ligand series consist of compounds of a single pharmacological type (e.g., agonists). Less common are reports of transcriptionally repressive NR inverse agonists and compounds that span the full pharmacological spectrum, from agonism to inverse agonism, which display distinct coregulator recruitment profiles or ligand bias.

Peroxisome proliferator-activated receptor gamma (PPARγ) is a lipid-sensing NR and target of antidiabetic drugs that function as pharmacological agonists. NMR studies revealed that in the absence of ligand, the apo-PPARγ LBD is a dynamic conformational ensemble and samples two or more structural conformations, which can be stabilized into a single active state upon binding an agonist ^9^. Hydrogen/deuterium exchange mass spectrometry (HDX-MS) revealed that binding of graded agonists differentially stabilize a critical structural element, helix 12, that is critical for NR activation ^10^ supporting a role for protein dynamics in the mechanism of graded PPARγ agonism ^11^. Crystal structures of PPARγ LBD bound to graded agonists typically show the same transcriptionally active LBD conformation ^12^. In contrast, functional profiling of graded agonists binding to the PPARγ in biochemical assays using purified LBD correlate well to graded transcriptional activation of full-length PPARγ in cells and NMR-detected shifts of the PPARγ LBD conformational ensemble from a ground state towards an active state ^13,14^.

The PPARγ inverse agonist, T0070907, was originally described as an antagonist ^15^ because it and a chemically related compound, GW9662 ^16^, bind covalently through a nucleophilic substitution mechanism to Cys285 within the LBD and blocked binding of other PPARγ ligands. However, subsequent studies showed these compounds do not block all ligands from binding PPARγ ^17–20^. Separate from their potential functions in blocking ligand binding, these compounds have distinct pharmacological properties. GW9662 is a transcriptionally neutral PPARγ ligand, whereas T0070907 is an inverse agonist that represses PPARγ transcription ^21^. T0070907 and GW9662 share the same 2-chloro-5-nitrobenzamide scaffold, and substitutions at the amide R_1_ group can be modified to elicit PPARγ agonism ^22^ or inverse agonism ^21,23–26^. Covalent PPARγ inverse agonists are currently being developed as potential therapeutics in bladder cancer where hyperactivation of PPARγ transcription occurs ^27–30^. Crystal structures of PPARγ LBD bound to corepressor peptide and inverse agonists analogs of T0070907 ^23–26^ have provided insight into low-energy structural snapshots of PPARγ forced into a fully repressive state by the peptide. However, these static structures do not inform on the structural mechanism, or conformation-activity relationship, behind the differential graded activity observed within a ligand series during structure-activity relationship development, as they do not elucidate how compounds with graded activity influence the dynamic PPARγ LBD conformational ensemble.

We surmised that the 2-chloro-5-nitrobenzamide scaffold offers a unique opportunity to understand how ligands with graded activity across the entire pharmacological spectrum influence the PPARγ LBD conformational ensemble. We found that relatively minimal chemical modifications to this scaffold yielded a ligand series spanning graded inverse agonism to agonism. Crystal structures of PPARγ LBD bound with ligands and corepressor peptide, along with density functional theory (DFT) calculations, provided structural insight into improved inverse agonist efficacy. NMR studies revealed that compounds within the ligand series influence the PPARγ LBD conformational ensemble by selecting active- and repressive-like states that are natively populated in the apo-LBD ensemble.

## RESULTS

### Design hypothesis to improve inverse agonism

Crystal structures of PPARγ LBD in transcriptionally active and repressive conformations suggest features that may be important for inverse agonism imparted by T0070907 ^23^. In the transcriptionally active conformation, where PPARγ LBD is bound to agonist (e.g., rosiglitazone) and coactivator (e.g., TRAP220/MED1) peptide, helix 12 is solvent exposed and forms the AF-2 surface along with helix 3–5 for LXXLL-containing motifs in coactivator proteins (**Figure 1A**). In contrast, in the repressive conformation helix 12 occupies the orthosteric ligand-binding pocket leaving the remaining AF-2 surface regions exposed for binding the longer corepressor peptide motif. Second, a pi-stacking interaction is observed between the polar T0070907 pyridyl group and three residues (His323, His449, and Tyr473) forming an “aromatic triad” that includes a water-bridged interaction between the pyridyl nitrogen and the His323 imidazole ring (**Figure 1B**).

**Figure 1.**
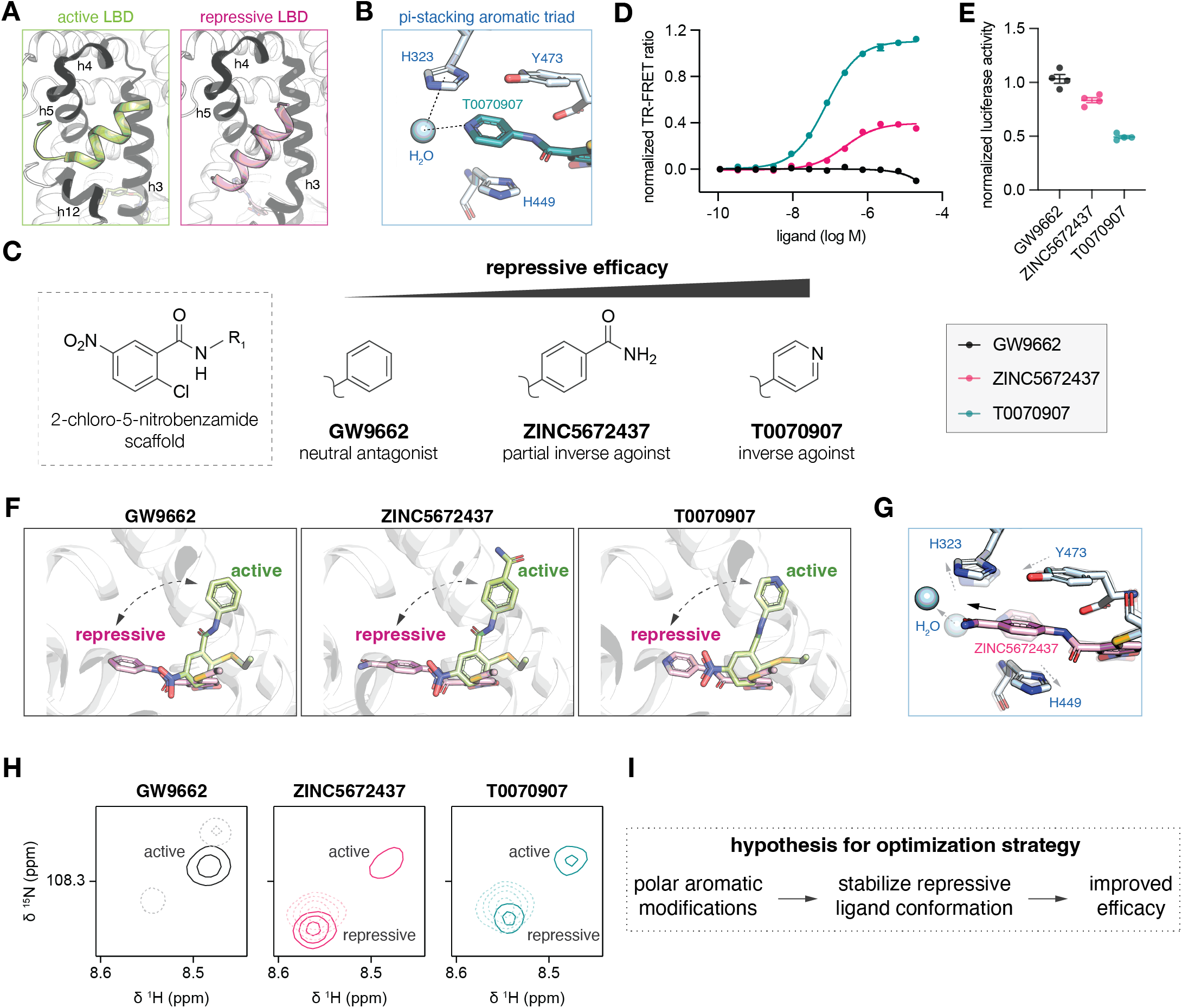
Structure-function data inform a design hypothesis to improve PPARγ inverse agonism. (**A**) AF-2 surface differences between PPARγ LBD in the transcriptionally active conformation bound to agonist rosiglitazone and TRAP220/MED1 coactivator peptide (PDB 6ONJ) and repressive conformation bound to inverse agonist T0070907 and NCoR1 corepressor peptide (PDB 6ONI) peptides. Coactivator and corepressor peptides are shown in green and pink, respectively. (**B**) Pi stacking between the polar T0070907 pyridyl group and three residues (His323, His449, and Tyr473) and a bridging water molecule form in the transcriptionally repressive conformation (PDB 6ONI). (**C**) Compound scaffold of parent molecules containing polar pyridyl (T0070907) and hydrophobic phenyl (GW9662) groups in relation to the polar benzamide group in ZINC5672437. (**D**) TR-FRET NCoR1 corepressor biochemical interaction assay (n=3; mean ± s.e.m.). (**E**) Cellular luciferase transcriptional reporter assay in HEK293T cells (n=4; mean ± s.e.m.). (**F**) Crystal structures showing the compound flipped binding modes in transcriptionally active vs. repressive conformations when bound to GW9662 (PDB 3B0R and 8FHE), T0070907 (PDB 6C1I and 6ONI), or ZINC5672437 (PDB 8FHF and 8FHE). (**G**) ZINC5672437 pi stacking interaction in the repressive state (PDB 8FHE) where the benzamide replaces the bridging water in the T0070907-bound structure (PDB 6ONI). (**H**) 2D [^1^H,^15^N]-TROSY-HSQC NMR focused on Gly399 of ^15^N-labeled PPARγ LBD bound to GW9662, T0070907, or ZINC5672437 in the absence (solid lines) or presence (dashed lines) of NCoR1 corepressor peptide. (**I**) Data informed design hypothesis to improve PPARγ inverse agonism.

NMR studies also inform the structural mechanism of PPARγ inverse agonism. In the absence of ligand, helix 12 exchanges in and out of the orthosteric ligand-binding pocket in repressive- and active-like conformations, respectively, on the microsecond-to-millisecond time scale ^23^. T0070907 binding slows the rate of exchange between these two natively populated LBD conformations such that they are long-lived and simultaneously observed by NMR ^21,31,32^. The active-like T0070907-bound state binds coactivator peptides with high affinity, and the repressive-like state binds corepressor peptides with high affinity ^21^. Binding of GW9662, which contains a hydrophobic phenyl R_1_ moiety instead of the polar pyridyl group in T0070907, also exchanges between these two conformations, but primarily populates an active-like conformation that is forced into a repressive conformation upon binding corepressor peptide ^21,32^.

Taken together, these findings informed a hypothesis that 2-chloro-5-nitrobenzamide compounds with polar aromatic R_1_ groups may stabilize the transcriptional repressive conformation via pi-stacking of the R_1_ group with the aromatic triad (**Figure 1C**). To test this hypothesis, we searched the ZINC database ^33^ and discovered a commercially available compound (ZINC5672437) with a polar benzamide aromatic ring. ZINC5672437 displays partial corepressor-selective inverse agonism relative to GW9662 and T0070907 in a time-resolved fluorescence resonance energy transfer (TR-FRET) corepressor peptide interaction assay (**Figure 1D**) and a cell-based transcriptional reporter assay (**Figure 1E**). To determine the structural basis for this activity, we compared crystal structures of PPARγ LBD bound to the compounds in repressive and active LBD conformations. We previously reported repressive LBD conformation crystal structures where PPARγ LBD is bound to T0070907 and NCoR1 ID2 or SMRT ID2 corepressor peptides ^23^. We solved two new NCoR1 ID2 peptide-bound crystal structures of PPARγ LBD bound to GW9662 or ZINC5672437 each to 1.8Å resolution (**Supplementary Table S1**). For the active conformation structures, we compared structures where crystals of apo-PPARγ LBD, which adopt an active conformation, were soaked with GW9662 ^19^ or T0070907 ^21^, which displayed flipped ligand binding modes compared to the repressive conformation T0070907-bound structures ^23^. We used a similar method to obtain an active conformation crystal structure for ZINC5672437-bound PPARγ LBD to 2.1Å resolution (**Supplementary Table S1**).

The crystal structures show that all three compounds are capable of adopting the active- and repressive-like ligand conformations (**Figure 1F**). The benzamide moiety extended from the otherwise nonpolar aromatic ring in ZINC5672437 replaces the water-bridged T0070907 pyridyl interaction with His323 and pushes the bridging water towards the AF-2 surface (**Figure 1G**). 2D [^1^H,^15^N]-TROSY-HSQC NMR focused on Gly399, a residue located near the AF-2 coregulator interaction surface that is sensitive to ligand-dependent active and repressive conformations ^13,21^, reveals the presence of two long-lived ZINC5672437-bound LBD conformations or structural populations (**Figure 1H**) that are similar to the T0070907-bound active- and repressive-like conformations ^21^. Addition of NCoR1 ID2 corepressor peptide consolidated these two structural populations into a single conformation. In contrast, GW9662 populates an active-like LBD conformation in the absence of peptide and two distinct conformations when bound to NCoR1 peptide ^21^.

The biochemical, cellular, and NMR data show that ZINC5672437 functions as a partial PPARγ inverse agonist with graded activity between T0070907 and GW9662. Taken together, these data provide support for the aromatic triad-stabilizing hypothesis and indicate that chemical modifications containing aromatic polarity may stabilize a repressive PPARγ LBD conformation and improve transcriptional repression efficacy (**Figure 1I**).

### Ligand series that spans graded repression and activation

We synthesized twenty 2-chloro-5-nitrobenzamide analogs containing an aromatic R_1_ group decorated with different substituents to determine if we can improve efficacy and generate a ligand set with graded repressive activity with minimal chemical modifications (**Figure 2**). We tested the compounds in several biochemical and cellular ligand profiling assays that report on active and repressive PPARγ functions.

**Figure 2.**
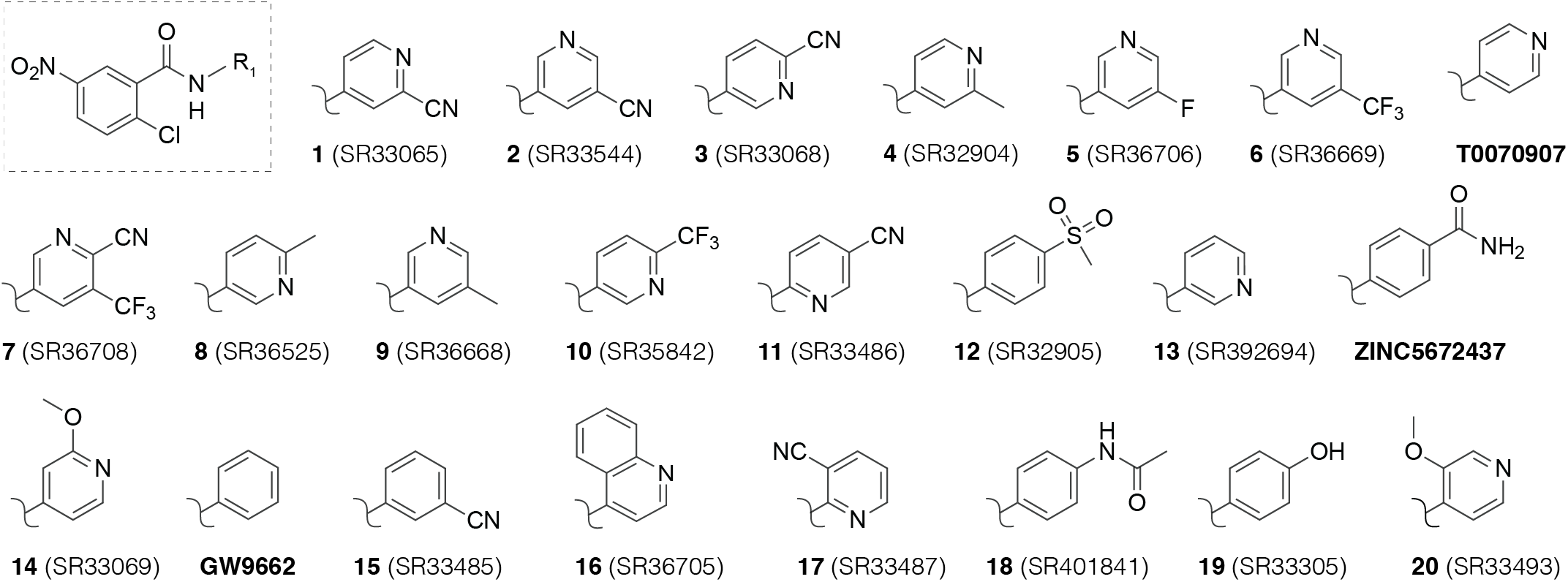
Compound analogs synthesized in this study. Compounds are numbered via the rank order in efficacy in the TR-FRET NCoR1 corepressor peptide interaction assay data shown in Figure 3.

Using TR-FRET biochemical assays, we determined how the analogs affect interaction between the PPARγ LBD and coregulator peptides derived from the NCoR1 corepressor and MED1 coactivator proteins (**Figure 3A**). In these experiments, the assay values report on the relative change in affinity or interaction efficacy between PPARγ LBD and the coregulator peptides; larger values correspond to increased interaction efficacy. To directly measure how the analogs affect coregulator peptide binding affinity to PPARγ LBD, we performed fluorescence polarization (FP) assays using NCoR1 corepressor and TRAP220/MED1 coactivator peptides (**Figure 3B**). To determine how the analogs affect PPARγ transcription, we performed a luciferase reporter assay where HEK293T cells were transfected with a full-length PPARγ expression plasmid along with a second plasmid containing three copies of the PPAR DNA-binding response element sequence (PPRE) upstream of luciferase gene and treated the cells with compounds (**Figure 3C**). Finally, to determine how the analogs affect PPARγ-mediated gene expression, we cultured 3T3-L1 preadipocytes in the presence of compounds and measured the expression of the adipogenic PPARγ target gene *aP2/FABP4* after two days of differentiation (**Figure 3D**).

**Figure 3.**
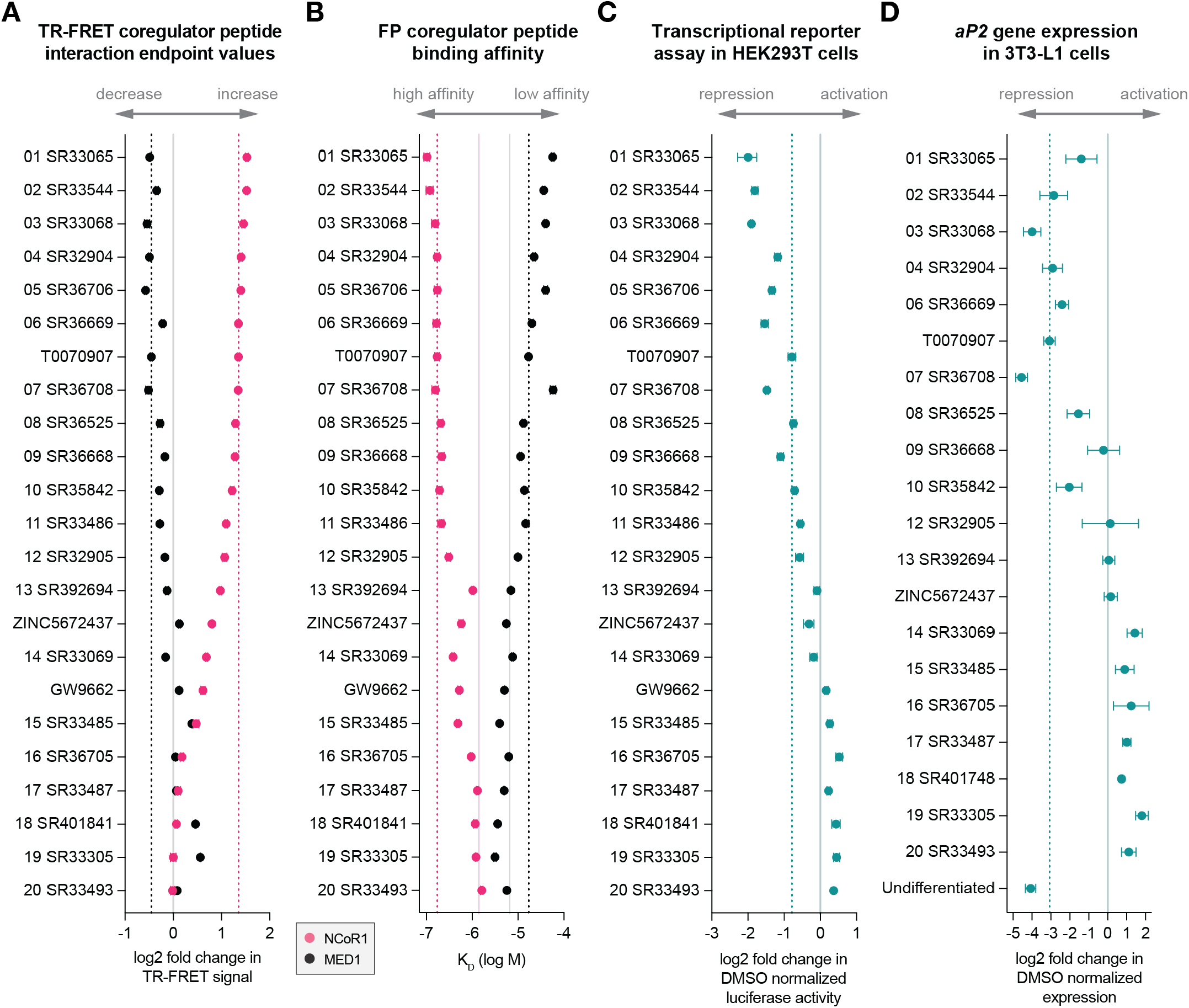
Compounds show graded activity in biochemical and cellular functional profiling assays. (**A**) Time-resolved fluorescence resonance energy transfer (TR-FRET) NCoR1 corepressor peptide biochemical interaction assay (n=3; mean ± s.e.m.). (**B**) Fluorescence polarization (FP) NCoR1 corepressor peptide binding assay (n=3) fitted affinity (mean ± s.d.). (**C**) Cellular luciferase transcriptional reporter assay in HEK293T cells (n=4; mean ± s.e.m.). (**D**) Expression of *aP2* in 3T3-L1 cells after two days of differentiation (n=3; mean ± s.d.). Solid vertical lines indicate activity for DMSO/control conditions, and dotted vertical lines indicate the activity of the parent inverse agonist T0070907.

The compound analogs show a wide range of graded activities spanning transcriptional repression and increased corepressor recruitment and binding affinity to transcriptional activation and increased coactivator recruitment and binding affinity. Approximately half of the compounds display similar or improved inverse agonism compared to T0070907; of these, compounds 3 (SR33068) and 7 (SR36708) are the most efficacious in repressing *aP2* expression in 3T3-L1 cells to levels similar to undifferentiated cells. Several analogs show properties of PPARγ agonism via increased coactivator peptide interaction, decreased corepressor peptide interaction, and increased transcription and *aP2* expression, including compounds 17 (SR33487), 19 (SR33305), and 20 (SR33493).

In our previous study of thiazolidinedione (TZD) PPARγ agonists, we found that graded activation within the TZD ligand series via increased coactivator peptide recruitment and binding affinity is highly correlated to cellular transcription ^13^. We calculated Spearman correlation coefficients from pairwise comparisons of the 2-chloro-5-nitrobenzamide biochemical and cellular profiling assay data (**Figure 4**). Transcription repression and decreased *aP2* expression are highly correlated to the NCoR1 TR-FRET and FP biochemical assays and highly anticorrelated to the MED1 TR-FRET and FP biochemical assay data. These data show that functional profiling of the compounds in biochemical microplate assays using purified PPARγ LBD protein explains the effect of the ligand series on the transcriptional activity of full-length PPARγ and the expression of a PPARγ target gene during adipogenesis.

**Figure 4.**
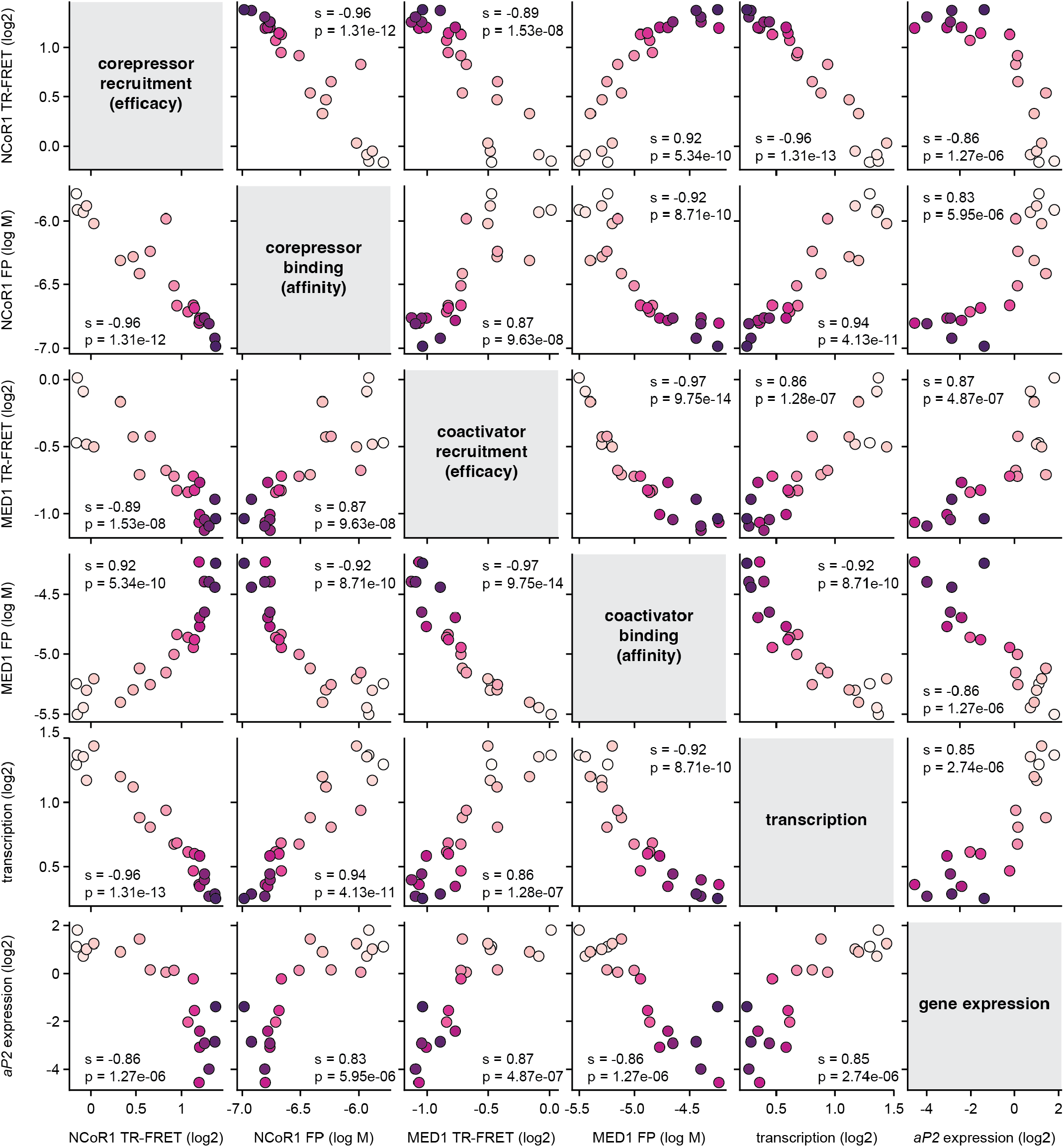
Pairwise correlation analysis of the compound profiling assay data. Spearman correlation coefficients and associated p-values are listed. Data are colored from white to purple according to compound numbers displayed in Figure 1.

### Crystal structures inform on improved corepressor-selective efficacy

To determine the structural basis of improved inverse agonism, we solved crystal structures of PPARγ LBD bound to NCoR1 corepressor peptide and compounds 2 (SR33544), 3 (SR33068), 4 (SR32904), 5 (SR36706), and 11 (SR33486) with resolutions ranging from 1.42Å to 2.22Å (**Supplementary Table S1**). Overall, the repressive PPARγ LBD conformation in these structures is highly similar to crystal structures of PPARγ LBD bound to NCoR1 corepressor peptide and GW9662, ZINC5672437, and T0070907 (0.15Å Cα rmsd).

The compounds contain a pyridyl nitrogen, which in T0070907 is at the third position, with additional R_1_ groups that interact with the aromatic triad residues (His323, His449, Tyr473) via pi-stacking interactions (**Figure 5**). Compounds 3 and 11 contain a cyano group at the third position with pyridyl nitrogens at the second and first positions, respectively, which substitutes for the water that bridges the interaction of T0070907 with His323. Compound 2 contains a pyridyl nitrogen at the second position pointing towards the NCoR1 corepressor peptide bound at the AF-2 surface, and a cyano group at the fourth position that points towards helix 11 with no water bridged interaction with His323. Compound 5 contains a pyridyl nitrogen at the second position that points towards the NCoR1 corepressor peptide, similar to compound 2, but contains a fluorine at the fourth position instead of a cyano group that points towards helix 11. Finally, compound 4 contains a pyridyl nitrogen at the third position similar to T0070907, which retains the water bridged interaction with His232, and a methyl substitution at the fourth position that points towards the NCoR1 corepressor peptide.

**Figure 5.**
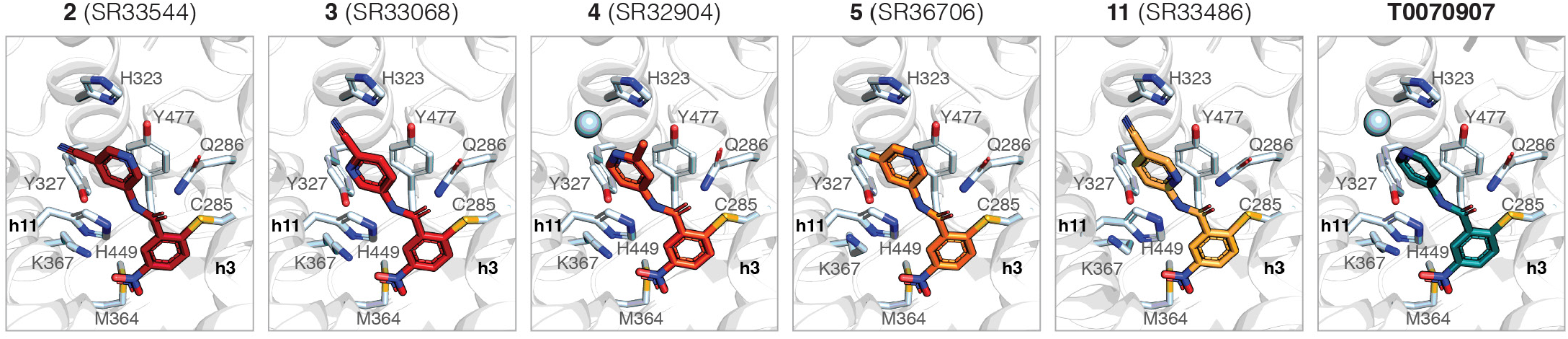
Inverse agonist binding modes in co-crystal structures of PPARγ LBD cobound to NCoR1 corepressor peptide. Structures are oriented with a view where the ligand R_1_ group is pointed towards the AF-2 surface. Residues annotated were considered in density functional theory (DFT) quantum mechanical (QM) calculations of ligand interaction free energies (ΔG_bind_), and α-helices are noted by h3 (helix 3) and h11 (helix 11).

We performed density functional theory (DFT) quantum mechanics (QM) calculations to estimate the interaction free energies (ΔG_bind_) between the compounds with residues comprising the pi-stacking aromatic triad residues (His323, His449, Tyr473) and others nearby residues (Cys285, Gln286, Tyr327, Met364, Lys367). For two compounds, 3 and 11, we also modeled a R_1_ ring conformation flipped by 180º and performed DFT QM calculation, as the electron density could not differentiate the final modeled or flipped orientations. Compared to T0070907, the substituted pyridyl R_1_ groups in compounds 2, 3, 5, and 11 afford more favorable ΔG_bind_ values (**Supplementary Table S2**). This suggests that the improved compound functional efficacy may originate in part from additional aromatic ring polarity and polar compound modifications projecting towards helix 11 or the bound corepressor peptide. However, among the pyridyl-substituted analogs, ΔG_bind_ is most favorable for compound 11, which displays the lowest efficacy among this group. Thus, there may be additional contributions from other residues, water-bridging interactions, or other phenomena not considered in the DFT-QM calculations such as compound structural rigidity that may contribute to experimental ligand efficacy.

### Ligands shift the functional LBD conformational ensemble

We posited that the improved inverse agonism within the ligand series could originate from two effects on the PPARγ LBD conformational ensemble: the ligands could either maintain the two long-lived repressive and active-like LBD conformations observed for T0070907-bound LBD ^21^ and further slow the rate of exchange between these states, or alternatively, the ligands could shift the ensemble towards and stabilize only the repressive-like conformation, while agonists would shift the ensemble towards an active-like conformation. To probe these mechanisms, we compared 2D [^1^H,^15^N]-TROSY-HSQC NMR data of ^15^N-labeled PPARγ LBD bound to the ligand series.

Focusing on Gly399, compounds with improved inverse agonism shift the LBD conformational ensemble towards a single peak representative of the repressive-like conformation (**Figure 6A**). Other improved analogs or compounds with partial inverse agonism show two NMR peaks, a repressive-like conformation peak and a weakly populated peak that is on-path (along the diagonal relative to the two T0070907-observed states) to the active-like conformation and in slow exchange on the NMR time scale (milliseconds or longer). Analogs within the ligand series that function as PPARγ agonists, compounds 15–20, stabilize either a single active-like conformation peak or show two NMR peaks that are shifted towards a graded agonism state similar to the graded TZD agonist series^13^. In addition to Gly399, other residues throughout the LBD with well-resolved NMR peaks show ligand-activity-dependent NMR peak shifts between repressive- and active-like states (**Supplementary Figures S1-S6**). We observed similar shifts for these residues in our previous study on T0070907 ^21^, including Arg234 (helix 2), Gly338 (β-sheet in the ligand-binding pocket), Arg350 (helix 6), Asn375 (helix 7), and Asp380 (helix 8-9 loop). This indicates the entire LBD is sensitive to the ligand-induced graded shift in the conformational ensemble between repressive- and active-like states.

**Figure 6.**
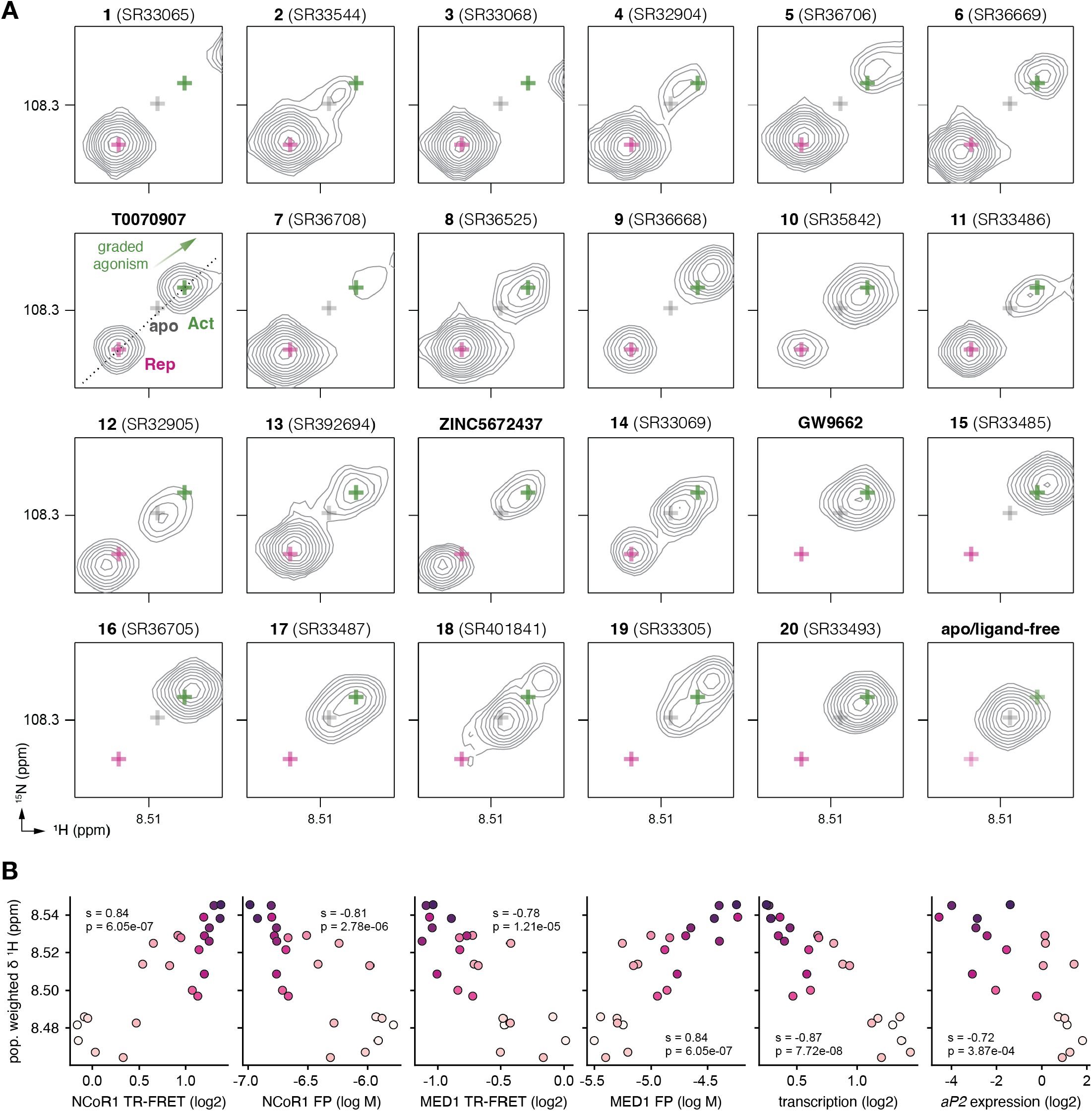
NMR analysis PPARγ LBD conformational ensemble describes ligand series function. (**A**) 2D [^1^H,^15^N]-TROSY-HSQC NMR focused on Gly399 of ^15^N-labeled PPARγ LBD bound to compounds in the ligand series. The peak positions of the active-like (Act) and repressive-like (Rep) states populated by T0070907 are noted with green and pink plus signs, respectively; and the peak position of the apo/ligand-free state is noted with a gray plus sign. The direction peaks shift for graded agonism, partial-to-full, is indicated with a white-to-green arrow. (**B**) Pairwise correlation plots between compound profiling data and weighted NMR chemical shift population shifts. Spearman correlation coefficients and associated p-values are listed. Data are colored from white to purple according to compound numbers displayed in Figure 1.

To quantitatively determine the influence of the ligand series on the PPARγ LBD conformational ensemble, we calculated population-weighted ^1^H NMR chemical shift values for Gly399 in ^15^N-labeled PPARγ LBD when bound to the compounds. This measurement reports on the degree to which the compounds shift the LBD ensemble between repressive- and active-like states. Spearman correlation coefficients calculated from pairwise comparisons of the population-weighted NMR chemical shifts and biochemical and cellular ligand profiling assays show a strong correlation (**Figure 6B**). However, there are limitations to this analysis, particularly the assumption that the populations of distinct, long-lived, ligand-bound LBD conformations are linked to the measured integrated volumes of well-resolved NMR peaks. This assumption may not be accurate, as dynamic motions on different time scales can influence NMR peak line shapes and thus integrated peak volumes. Furthermore, in some cases, there may be overlapped peaks that cannot be easily distinguished and picked separately for peak volume integration. Notwithstanding these limitations, the analysis indicates that the NMR-detected shift in the structural populations explains the graded activity within the ligand series with relatively high correlation spearman coefficients (>0.7), supporting the underlying hypothesis that compounds within the ligand series shift the PPARγ LBD conformational ensemble between active and repressive conformations.

## DISCUSSION

Structural biology studies of NRs focused on understanding the molecular basis for ligand-dependent changes in transcription have been primarily focused on generating crystal structures of NR LBDs bound to coregulator peptide and/or ligand. However, while crystal structures provide low-energy static snapshots of the functional endpoints (e.g., fully active states) they do not explain graded activity that is critical for understanding how ligands influence the dynamic NR LBD conformational ensemble. PPARγ is a good model system to study how pharmacological ligands influence the dynamic NR LBD conformational ensemble. Although crystal structures of PPARγ LBD bound to partial or full agonists all show a similar low-energy active LBD conformation and do not explain the molecular basis of graded agonism ^11,14^, NMR studies revealed that graded activation of PPARγ is correlated to shifting the LBD conformational ensemble from a ground state to an active state ^13^. Here, we define a more comprehensive LBD conformational ensemble and show that compounds within the same scaffold select natively populated repressive- and active-like conformations present in the apo/ligand-free state ^23^.

The ligand series we developed, based on the T0070907 2-chloro-5-nitrobenzamide compound scaffold, displays graded activity that selects LBD conformations detected by NMR along the transcriptionally repressive-to-active continuum. To the best of our knowledge, this NR ligand series is the first to span inverse agonism and agonism that selects natively populated active and repressive LBD conformations. Other reported PPARγ inverse agonists appear to function via a different mechanism that does not select for a natively populated repressive state. Instead, crystals structures of PPARγ LBD bound to these compounds show solvent exposed non-active helix 12 conformations with ligand binding modes that overlap with the repressive helix 12 conformation within the orthosteric pocket ^34,35^. These compounds may bind to the LBD similar to TZDs, which are orthosteric agonists that bind via a two-step induced fit mechanism with an initial encounter complex followed by transition into the orthosteric pocket, which displaces helix 12 from its apo-conformation within the orthosteric pocket ^36^. Thus, there appear to be two different mechanisms of PPARγ inverse agonism: ligands that select a native-like repressive conformation where helix 12 adopts a conformation within the orthosteric ligand-binding pocket, and ligands that bind to the orthosteric pocket via an induced fit mechanism competing with and displacing helix 12 from the orthosteric pocket.

Our work here shows that NMR enables conformation-activity relationship analysis and explains activities for a ligand series spanning transcriptional activation and repression. In contrast, crystal structures coupled to SAR studies only reveal the fully repressive LBD conformation and do not reveal the structural mechanism of graded repression. Recent studies have reported improved 2-chloro-5-nitrobenzamide PPARγ inverse agonists ^24–26^; these studies reported more elaborate R_1_ group chemical modifications in contrast to the relatively minimal chemical modifications that we show here are sufficient to push the LBD conformational ensemble towards a fully repressive state. With the growing interest in developing PPARγ inverse agonists for bladder cancer therapeutics, including an analog similar to the 2-chloro-5-nitrobenzamide scaffold that is progressing toward phase 1 clinical trials ^37,38^, our findings demonstrate a platform that can structurally assess, explain, and potentially predict the activity of PPARγ compounds ranging from agonism to inverse agonism.

## MATERIALS AND METHODS

### Ligands and peptides

Commercially available compounds, T0070907 (CAS 313516-66-4) and GW9662 (CAS 22978-25-2), were purchased from Cayman Chemical. Details for the new compounds reported in this study can be found in **Supplementary File 1**. Peptides derived from human NCoR1 ID2 (2256-2278; DPASNLGLEDIIRKALMGSFDDK) and human TRAP220/ MED1 ID2 (residues 638–656; NTKNHPMLM NLLKDNPAQD) were synthesized by LifeTein with an amidated C-terminus for stability, with or without a N-terminal FITC label and a six-carbon linker (Ahx).

### Cell lines for mammalian cell culture

HEK293T (ATCC #CRL-11268) and 3T3-1L (ATCC #CL-173) cells were cultured according to ATCC guidelines. HEK293T cells were grown at 37°C and 5% CO_2_ in Dulbecco’s Modified Eagle Medium, high glucose, GlutaMAX Supplement (DMEM+GlutaMAX, Gibco) supplemented with 10% fetal bovine serum (FBS, Gibco), 100 units/mL of penicillin, 100 μg/ mL of streptomycin (Gibco) until 90 to 95% confluence in T-75 flasks prior to subculture or use. 3T3-L1 cells were grown at 37 °C and 5% CO_2_ in DMEM, high glucose, GlutaMAX, pyruvate (Gibco) supplemented with 10% bovine calf serum (BCS, Gibco) and 100 units/mL of penicillin, 100 μg/mL of streptomycin (Gibco) until 70% confluence in T-75 flasks prior to subculture or use.

### Protein expression and purification

Human PPARγ ligand-binding domain (LBD) protein, residues 203-477 (isoform 1 numbering), was expressed as a TEV-cleavable N-terminal hexa-his-tag fusion protein (6xHis-PPARγ LBD). Expression was performed in BL21(DE3) *Escherichia coli* cells in either autoinduction ZY media (unlabeled protein) or using M9 minimal media supplemented with ^15^N ammonium chloride (for NMR studies). For autoinduction, cells were grown at 37 °C for 5 hours, 30 °C for 1 hour, and 18°C for 16 hours before harvesting by centrifugation (4,000g, 30 min). For minimal media, cells were grown until the OD_600_ was 0.6 before adding 0.5 mM (final concentration) isopropyl β-D-thiogalactoside (IPTG) and incubating at 18 °C for 16 h before harvesting by centrifugation (4,000g, 30 min). Cells were resuspended in lysis buffer (50 mM potassium phosphate (pH 7.4), 500 mM KCl, 10 mM imidazole) and lysed by sonication on ice. Cell lysate was clarified by centrifugation (20,000g, 30 min) and filtration (0.2 μm). For His-tagged PPARγ-LBD, the protein was purified using Ni-NTA affinity chromatography followed by size exclusion chromatography (Superdex 75) on an AKTA pure in assay buffer (20 mM potassium phosphate, pH 7.4, 50 mM KCl, 0.5 mM EDTA). For NMR and crystallography, the His-tag was cleaved with TEV protease in dialysis buffer (20 mM potassium phosphate pH 7.4, 200 mM KCl) overnight at 4 °C. The cleaved protein was reloaded on the Ni-NTA column, the flow through collected, and further purified by size exclusion chromatography (Superdex 75) in assay buffer (20 mM potassium phosphate, pH 7.4, 50 mM KCl, 0.5 mM EDTA). Protein was confirmed to be >95% pure by SDS-PAGE. Purified samples were stored at –80 °C.

### Time-resolved fluorescence resonance energy transfer (TR-FRET) coregulator interaction assays

Assays were performed in black 384-well plates (Greiner) with 23 μL final well volume. For the coregulator recruitment assay, each well containing 4 nM 6xHis-PPARγ LBD, 1 nM LanthaScreen Elite Tb-anti-His Antibody (ThermoFisher), and 400 nM FITC-labeled NCoR1 ID2 or MED1 ID2 peptide in a buffer containing 20 mM potassium phosphate (pH 7.4), 50 mM potassium chloride, 5 mM TCEP, 0.005% Tween 20. Ligands were added as a single concentration (10 μM). Compounds stocks were prepared in DMSO, added to wells in triplicate, and plates were read using BioTek Synergy Neo multimode plate reader after incubation at 25 °C for 1 h. The Tb donor was excited at 340 nm; the Tb donor emission was measured at 495 nm, and the acceptor FITC emission was measured at 520 nm. Data were plotted using GraphPad Prism as TR-FRET ratio 520 nm/495 nm. Data are representative of two or more independent experiments.

### Fluorescence polarization coregulator interaction assays

Assays were performed using 6xHis-PPARγ LBD preincubated with or without 2 molar equivalents of compound at 4 °C overnight and buffer exchanged to remove excess ligand. Protein samples were serially diluted into a buffer containing 20 mM potassium phosphate (pH 8), 50 mM potassium chloride, 5 mM TCEP, 0.5 mM EDTA, and 0.01% Tween-20 and plated with 180 nM FITC-labeled NCoR1 ID2 or TRAP220/MED1 ID2 peptide in black 384-well plates (Greiner).The plate was incubated at 25°C for 1 hr, and fluorescence polarization was measured on a BioTek Synergy Neo multimode plate reader at 485 nm emission and 528 nm excitation wavelengths. Data were plotted using GraphPad Prism as fluorescence polarization signal in millipolarization units vs. protein concentration and fit to a one site — total binding equation using a consistent, fixed Bmax value as binding for some conditions did not saturate at the highest protein concentration used (45 μM). Data are representative of two or more independent experiments.

### Transcriptional reporter assays

HEK293T cells were cultured in Dulbecco’s minimal essential medium (DMEM, Gibco) supplemented with 10% fetal bovine serum (FBS) and 50 units ml−1 of penicillin, streptomycin, and glutamine. Cells were grown to 90% confluency in T-75 flasks; from this, 2 million cells were seeded in a 10-cm cell culture dish for transfection using X-tremegene 9 (Roche) and Opti-MEM (Gibco) with full-length human PPARγ isoform 2 expression plasmid (4 μg), and a luciferase reporter plasmid containing the three copies of the PPAR-binding DNA response element (PPRE) sequence (3xPPRE-luciferase; 4 μg). After an 18-h incubation, cells were transferred to white 384-well cell culture plates (Thermo Fisher Scientific) at 10,000 cells/well in 20 μL total volume/well. After a 4 h incubation, cells were treated in quadruplicate with 20 μL of either vehicle control (1.5% DMSO in DMEM media) or 5 μM ligand. After a final 18-h incubation, cells were harvested with 20 μL Britelite Plus (PerkinElmer), and luminescence was measured on a BioTek Synergy Neo multimode plate reader. Data were plotted in GraphPad Prism as mean ± s.e.m. and are representative of two or more independent experiments.

### Gene expression analysis

3T3-L1 cells were cultured in DMEM medium supplemented with 10% FBS and 50 units ml–1 of penicillin, streptomycin, and glutamine. Cells were grown to 70% confluency and then seeded in 12-well dishes at 50,000 cells per well and incubated overnight at 37 °C, 5% CO2. The following day, cells were treated with media supplemented with 0.5 mM 3-iso-butyl-1-methylxanthine, 1 μM dexamethasone, and 877 nM insulin. Following 2-days of incubation, cells were treated with 10 μM compound in media supplemented with 877 nM insulin for 24 h. RNA was extracted using quick-RNA MiniPrep Kit (Zymo) and used to generate complementary DNA using qScript cDNA synthesis kit (Quantabio). Expression levels of the PPARγ target gene *aP2/FABP4* (forward primer: 5’-AAGGTGAAGAGCATCATAACCCT-3’) and the housekeeping gene *TBP* (forward primer: 5’-ACCCTTCACCAATGACTCCTATG-3’) used for normalization was measured using Applied Biosystems 7500 Real-Time PCR system. Relative gene expression was calculated via the ddCt method using Applied Biosystems Relative Quantitation Analysis Module Software, which reported values as mean with upper and lower limits. Data were plotted in GraphPad Prism and are representative of two or more independent experiments.

### NMR spectroscopy

Two-dimensional [^1^H,^15^N]-TROSY HSQC NMR data of ^15^N-labeled PPARγ LBD (200 μM) were acquired at 298 K on a Bruker 700 MHz NMR instrument equipped with a QCI cryoprobe in NMR buffer (50 mM potassium phosphate, 20 mM potassium chloride, 1 mM TCEP, pH 7.4, 10% D2O) with ligands preincubated overnight at 4 ºC with 2 molar equivalents and buffer exchanged to remove excess ligand. Data were collected using Topspin 3.0 (Bruker Biospin) and processed/ analyzed using NMRFx ^39^. NMR chemical shift assignments previously transferred from rosiglitazone-bound PPARγ LBD ^14^ to T0070907- and GW9662-bound states ^21,23^ were used in this study for well-resolved residues with conversed NMR peak positions to the previous ligand-bound forms using the minimum chemical shift perturbation procedure ^40^. Population-weighted ^1^H NMR chemical shift analysis was performed focusing on the NMR peaks observed for Gly399 in each ligand-bound NMR spectrum. Peak volumes for well-resolved peaks (e.g., one peak = one state/conformation, two peaks = two states/conformations in slow exchange) were calculated using an elliptical peak fitting algorithm. Population weighted average ^1^H chemical shift values were calculated with Python using Jupyter Notebook using NumPy and Pandas packages using an equation (‘lambda x: np.average(x.H1_P, weights=x.weighted_ vol)’)) with two inputs: peak volumes estimated the relative population sizes (weighted_vol) of each state, and the ^1^H NMR chemical shift values of each state (H1_P).

### Crystallization and structure determination

Two molar equivalents of compounds were added to purified PPARγ-LBD (1:2 protein/compound ratio) and incubated at 4 °C overnight, followed by the addition of 5 molar equivalents of appropriate peptides (1:5 protein/peptide ratio) and a final incubation at 4 °C overnight. Protein complexes were purified by gel filtration (Superdex 75) on an AKTA pure, in assay buffer (20 mM potassium phosphate, pH 7.4, 50 mM KCl, 0.5 mM EDTA), and concentrated to 10 mg/mL. Crystals of the protein complex were obtained by sitting-drop vapor diffusion against 50 μL of reservoir solution (100 mM MES [pH 6.5], 200 mM ammonium sulfate, 30% PEG 8,000) at a 1:1 protein/reservoir solution ratio using 96-well crystallization plates at 22 °C. Crystals were transferred to cryoprotectant (reservoir solution plus 10% ethylene glycol) and flash-frozen in liquid nitrogen prior to data collection at the SLAC National Accelerator Laboratory/Stanford Synchrotron Radiation Lightsource (SSRL) beamline 12-2. Data were processed with XDS and scaled with Aimless. Structures were solved by molecular replacement using Phaser in the Phenix software package, using the previously published crystal structure of PPARγ bound to T0070907 and NCoR1 ID2 peptide (PDB code: 6ONI) as the search model. Structures were built with iterative rounds of manual model rebuilding in COOT followed by refinement using phenix.refine. Pairwise structural alignment and rmsd calculations were performed via the RCSB webserver (https://www.rcsb.org/alignment/) using the jFATCAT rigid structural alignment algorithm.

### Density functional theory (DFT) quantum mechanical (QM) binding free energy calculations

We evaluated the binding free energies of ligands through the QM cluster method. For each PPARγ-ligand complex, we curated a cluster structure comprising the ligand and binding site residues (i.e., His323, His449, Tyr473, Cys285, Gln286, Tyr327, Met364, Lys367) based on its crystal structure ^23^. The cluster is further converted into the complex, ligand, and binding site models for subsequent binding free energy calculation. Using Gaussian16 ^41^, these structural models were optimized with PBE0-D3/def2-SVP method under fixed Cα and Cβ coordinates, followed by a single point energy correction with PBE0-D3/def2-TZVP method. Each optimized structure underwent normal mode analysis with PBE0-D3/def2-SVP method to validate that it is a stationary point on the potential energy surface. The analysis also informs the free energy correction value, which was further corrected using quasi-harmonic method ^42^ and then added to the single point energy to give the standard Gibbs free energy of the structure. Eventually, the binding free energy was calculated by subtracting the free energy of the complex by the free energy sum of the ligand and binding site.

### Statistical analysis of ligand profiling data

Correlation data plotting and analysis of Spearman (s) correlation coefficients and two-sided alternative hypothesis p-value testing of the biochemical and cellular ligand profiling data were performed in Python using Jupyter Notebook using several libraries including seaborn, matplotlib, numpy, and scipy. Reported p-values (p) represent the probability that the absolute value of the spearman or pearson coefficient of random (x,y) value drawn from the population with zero correlation would be greater than or equal to abs(r or s), according to the scipy stats documentation.

## ACKNOWLEDGMENTS

We thank Paola Munoz-Tello, Christopher Williams, and Zane Laughlin for critical reading and discussions. This work was supported in part by the National Institutes of Health (NIH) grant R01DK124870 from the National Institute of Diabetes and Digestive and Kidney Diseases (NIDDK). Use of the Stanford Synchrotron Radiation Lightsource, SLAC National Accelerator Laboratory, is supported by the U.S. Department of Energy, Office of Science, Office of Basic Energy Sciences under Contract No. DE-AC02-76SF00515. The SSRL Structural Molecular Biology Program is supported by the DOE Office of Biological and Environmental Research and by the NIH National Institute of General Medical Sciences (NIGMS) grant P30GM133894. The contents of this publication are solely the responsibility of the authors and do not necessarily represent the official views of NIDDK, NIGMS, NIH, or DOE.

## DATA AVAILABILITY

Crystal structures generated during the current study are available in the Protein Data Bank (PDB) under accession codes 8FHE, 8FHG, 8FHF, 8FKC, 8FKD, 8FKE, 8FKF, and 8FKG as detailed in **Supplementary Table S1**. The source data underlying Figure 3A-D, Figure 4, and Figure 6B are provided in a separate file, **Source Data 1** (Microsoft Excel file). The input and output files for all structural models used in the DFT QM calculations are provided in **Source Data 2** (ZIP file). ^1^H and ^13^C NMR data for all compounds synthesized are provided in **Supplementary File 1**. All other datasets generated and/or analyzed during the current study are available from the corresponding author on reasonable request.

## AUTHOR CONTRIBUTIONS

B.M. and D.J.K. conceived and designed the research. D.Z. and T.M.K. synthesized compounds. B.M. and J.S. expressed and purified protein and determined crystal structures. B.M. performed biochemical and cellular assays and collected NMR data. Q.Z. and Z.J.Y. performed DFT QM calculations. D.K. supervised the research and wrote the manuscript along with B.S. and input from all authors who approved the final version.

