## Supplemental File 1 for "Ligand efficacy shifts a nuclear receptor conformational ensemble between transcriptionally active and repressive states"

***General information:.*** NMR spectra were recorded in DMSO-*d6,* MeOD or deuterated chloroform ($CDCl_3$) on a Bruker AVANCE DPX-400 (400 MHz) and an AVANCE III 600 (600 MHz) spectrometer. Chemical shifts (δ) were calibrated relative to solvent peaks and are reported in parts per million (ppm), whereas the coupling constants (*J*) are reported in Hertz (Hz). Abbreviations for the peak multiplicities are as follows: s (singlet), brs (broad singlet), d (doublet), t (triplet), q (quartet), dd (doublet of doublets), dt (doublet of triplets) and m (multiplet). Mass spectrometry (MS) was performed using electrospray ionization on an Agilent 6230 TOF LC/MS mass spectrometer in the positive ion mode. The purity (≥ 95%) of all final synthesized compounds was determined by NMR. Compounds are listed by the chemical name (compound number, internal referencing code).

***Chemical Synthesis of Analogs:***

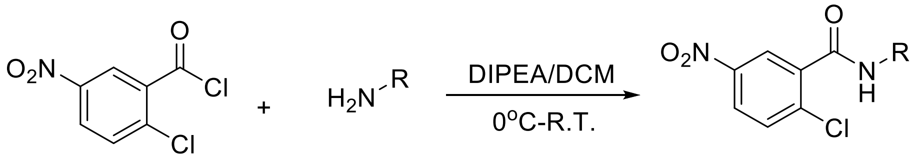


To a solution of aniline (1.1 mmol, 1.1 equiv.) and diisopropylethylamine (0.2 mL, 1.1 mmol, 1.1 equiv.) in dichloromethane (DCM, 1.5 mL) was added a solution of 2-chloro-5-nitrobenzoyl chloride (220 mg, 1mmol, 1.0 equiv.) in DCM (0.5L) dropwise in an ice-water bath. After 15 min, the reaction was allowed to warm to room temperature for 3-8 hours. The reaction mixture was diluted with ethyl acetate (EtOAc, 10 mL), washed with water (10 mL), saturated $NaHCO_3$ (10 mL) and brine (10 mL). The organic layer was dried over $MgSO_4$, filtered, and concentrated under reduced pressure. The crude material was purified by silica gel chromatography to yield the title compound as a white solid.

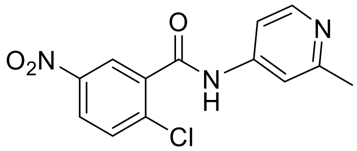


**SR32904**

**2-chloro-N-(2-methylpyridin-4-yl)-5-nitrobenzamide**

**$^1$HNMR ($CDCl_3$, 600 MHz)**, δ(ppm): 8.61 (d, *J*=2.8 Hz, 1H), 8.47 (d, *J*=6.0 Hz, 1H), 8.30 (dd, $J_1$=2.8Hz, $J_2$=8.0 Hz, 1H), 8.02 (br, 1H), 7.70 (d, *J*=9.0 Hz, 1H), 7.51 (s, 1H), 7.36 (d, *J*=4.2 Hz, 1H), 2.58 (s, 3H). **$^{13}$CNMR ($CDCl_3$, 150 MHz)**, δ (ppm): 162.56, 160.18, 150.33, 146.75, 144.23, 137.41, 135.72, 131.84, 126.50, 125.48, 113.20, 111.26, 24.65. **LCMS (ESI)**: Expected mass for $C_{13}H_{11}ClN_3O_3$ $(M + H)^+$: 292.05 Da, found 292.05 Da.

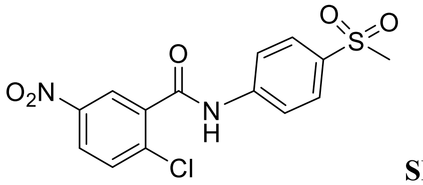


**SR32905**

**2-chloro-N-(4-(methylsulfonyl)phenyl)-5-nitrobenzamide**

**$^1$HNMR (DMSO-d6, 400 MHz)**, δ (ppm): 11.16 (s, 1H), 8.55 (d, *J*=2.9 Hz, 1H), 8.37 (dd, $J_1$=2.9Hz, $J_2$=8.8 Hz, 1H), 7.95 (s, 4H), 7.92 (d, *J*=8.8 Hz, 1H), 3.20 (s, 3H); **$^{13}$CNMR ($CDCl_3$, 100 MHz)**, δ (ppm): 163.90, 146.63, 143.33, 137.54, 137.49, 136.14, 131.90, 128.80, 126.52, 124.43, 120.20, 44.24. **LCMS (ESI)**: Expected mass for $C_{14}H_{12}ClN_2O_5S$ $(M + H)^+$: 355.02 Da, found 355.31 Da.

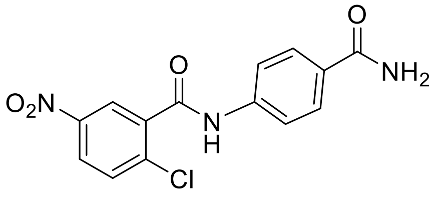

**SR33064**

**N-(4-carbamoylphenyl)-2-chloro-5-nitrobenzamide**

**$^{1}$HNMR (DMSO-d6, 400 MHz)**, δ (ppm): 10.93 (s, 1H), 8.51 (d, *J*=2.7 Hz, 1H ), 8.36 (dd, $J_1$=2.7Hz, $J_2$=8.9 Hz, 1H ), 7.92 (br, 2 H), 7.91 (d, *J*=8.7 Hz, 2H), 7.77 (d, *J*=8.7Hz, 2H), 7.31 (br, 1H). **$^{13}$CNMR ($CDCl_3$, 100 MHz)**, δ (ppm): 168.30, 163.74, 146.58, 141.47, 137.62, 137.50, 131.87, 129.98, 128.97, 126.37, 124.23, 119.58. **LCMS (ESI)**: Expected mass for $C_{14}H_{11}ClN_3O_4$ $(M + H)^+$: 320.04 Da, found 320.04 Da.

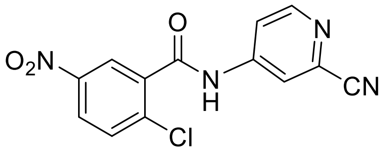

**SR33065**

**2-chloro-N-(2-cyanopyridin-4-yl)-5-nitrobenzamide**

**$^{1}$HNMR (DMSO-d6, 400 MHz)**, δ (ppm): 11.50 (s, 1H), 8.70 (d, *J*=5.8 Hz, 1H), 8.61 (d, *J*=2.5Hz, 1H), 8.40 (dd, $J_1$=2.8Hz, $J_2$=8.4 Hz, 1H), 8.21 (d, *J*=1.8 Hz, 1H), 7.95 (d, *J*=8.6Hz, 1H), 7.91 (dd, $J_1$=2.2Hz, $J_2$=5.4 Hz, 1H); **$^{13}$CNMR ($CDCl_3$, 100 MHz)**, δ (ppm): 164.61, 153.03, 146.68, 146.61, 137.48, 136.67, 133.88, 132.04, 126.93, 124.61, 119.05, 117.86, 117.42. **LCMS (ESI)**: Expected mass for $C_{13}H_8ClN_4O_3$ $(M + H)^+$: 303.03 Da, found 303.03 Da.

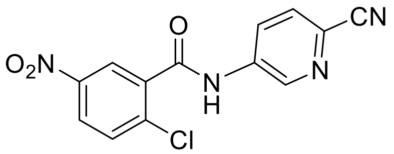

**SR33068**

**2-chloro-N-(6-cyanopyridin-3-yl)-5-nitrobenzamide**

**$^{1}$HNMR (DMSO-d6, 400 MHz)**, δ (ppm): 11.44 (s, 1H), 8.96 (d, *J*=2.2Hz, 1H), 8.61 (d, *J*=2.9Hz, 1H), 8.40 (t, *J*=2.6Hz, 1H), 8.38 (t, *J*=2.6Hz, 1H), 8.08 (d, *J*=8.7 Hz, 1H), 7.94 (d, *J*=8.7 Hz, 1H). **$^{13}$CNMR ($CDCl_3$, 100 MHz)**, δ (ppm): 164.21, 146.63, 142.87, 138.91, 137.54, 136.99, 132.00, 130.24, 127.42, 127.22, 126.80, 124.69, 118.05. **LCMS (ESI)**: Expected mass for $C_{13}H_8ClN_4O_3$ $(M + H)^+$: 303.03 Da, found 303.18 Da.

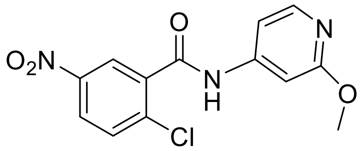

**SR33069**

**2-chloro-N-(2-methoxypyridin-4-yl)-5-nitrobenzamide**

**$^{1}$HNMR (DMSO-d6, 400 MHz)**, δ (ppm): 11.02 (s, 1H), 8.53 (d, *J*=2.5 Hz, 1H), 8.37 (dd, $J_1$=2.8Hz, $J_2$=8.8 Hz, 1H), 8.11 (dd, $J_1$=1.9 Hz, $J_2$=5.5 Hz, 1H), 7.92 (d, *J*=8.8Hz, 1H), 7.20-7.21 (m, 2H), 3.86 (s, 3H). **$^{13}$CNMR ($CDCl_3$, 100 MHz)**, δ (ppm):165.08, 164.33, 148.58, 147.97, 146.60, 137.80, 137.46, 131.87, 126.36, 124.44, 109.23, 99.76,53.69. **LCMS (ESI)**: Expected mass for $C_{13}H_{11}ClN_3O_4$ $(M + H)^+$: 308.04 Da, found 308.04 Da.

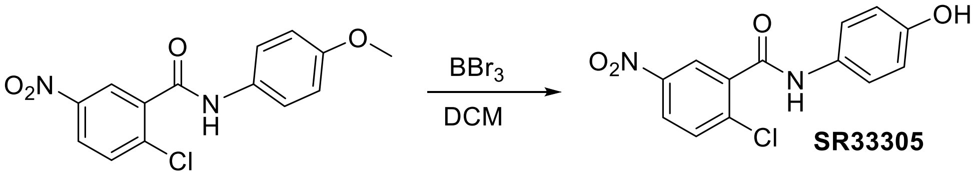

**SR33305**

**2-chloro-N-(4-hydroxyphenyl)-5-nitrobenzamide**

To a stirred solution of 2-chloro-N-(4-methoxyphenyl)-5-nitrobenzamide (0.12 g, 0.4 mmol) in dry DCM (5 mL), cooled to -78°C, was added slowly $BBr_3$ (0.8 mL of a 1M solution, 0.8 mmol) and the reaction was allowed to warm slowly to room temperature with stirring overnight before being quenched with water. The organic layer was separated and dried over $MgSO_4$, filtered, and concentrated under reduced pressure. The crude material was purified by silica gel chromatography to yield the title compound as a white solid. **$^1$HNMR (DMSO-d6, 400 MHz)**, δ (ppm): 11.02 (s, 1H), 8.54 (d, *J*=2.7 Hz, 1H), 8.37 (dd, $J_1$=2.6Hz, $J_2$=8.8 Hz, 1H), 7.92 (d, *J*=8.8Hz, 1H), 7.53 (d, *J*=7.2 Hz, 1H), 7.00 (s, 1H), 6.60 (d, *J*=6.9Hz, 1H). **$^{13}$C NMR ($CDCl_3$, 100 MHz)**, δ (ppm): 162.65, 154.54, 146.59, 138.45, 137.60, 131.77, 130.58, 125.98, 124.25, 122.01, 115.66. **LCMS (ESI)**: Expected mass for $C_{13}H_{10}ClN_2O_4$ $(M + H)^+$: 293.03 Da, found 293.07 Da.

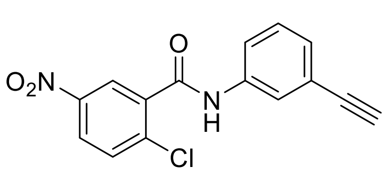

**SR33485**

**2-chloro-N-(3-ethynylphenyl)-5-nitrobenzamide**

**$^1$HNMR (DMSO, 400 MHz)**, δ (ppm): 8.62 (d, *J*=2.7 Hz, 1H), 8.29 (dd, $J_1$=8.9Hz, $J_2$=2.8Hz, 1H), 7.85 (br, 1H), 7.76 (s, 1H), 7.70-7.66 (m, 2H), 7.39-7.33 (m, 2H), 3.11 (s, 1H). **$^{13}$CNMR (DMSO, 100 MHz)**, δ (ppm):163.48, 146.64, 139.13, 137.89, 137.53, 131.86, 129.90, 127.95, 126.32, 124.38, 123.09, 122.63, 120.89, 83.63, 81.33. **LCMS (ESI)**: Expected mass for $C_{15}H_{10}ClN_2O_3$ $(M + H)^+$: 301.04 Da, found 301.28 Da.

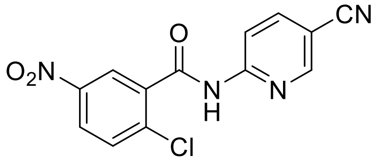

**SR33486**

**2-chloro-N-(5-cyanopyridin-2-yl)-5-nitrobenzamide**

**$^1$HNMR (DMSO-d6, 400 MHz)**, δ (ppm): 11.78 (s, 1H), 8.85 (t, *J*=1.45 Hz, 1H), 8.55 (d, *J*=2.90Hz, 1H), 8.33-8.36 (m, 3H), 7.87 (d, *J*=9.3Hz, 1H). **$^{13}$C NMR (DMSO, 150 MHz)**, δ (ppm): 164.71, 154.51, 152.67, 146.48, 142.81, 137.48, 137.14, 131.71, 126.56, 124.75, 117.60, 114.23, 104.89. **LCMS (ESI)**: Expected mass for $C_{13}H_8ClN_4O_3$ $(M + H)^+$: 303.03 Da, found 303.25 Da.

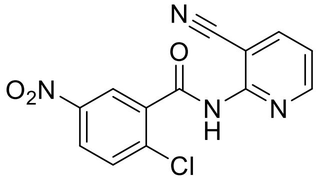

**SR33487**

**2-chloro-N-(3-cyanopyridin-2-yl)-5-nitrobenzamide**

**$^1$HNMR (DMSO-d6, 400 MHz)**, δ (ppm): 11.71 (s, 1H), 8.77 (dd, $J_1$=1.7Hz, $J_2$=4.8 Hz, 1H), 8.45 (dd, $J_1$=1.9Hz, $J_2$=7.7 Hz, 1H), 8.38 (m, 2H), 7.93 (d, *J*=9.4Hz, 1H), 7.56 (dd, $J_1$=4.8Hz, $J_2$=7.8Hz, 1H). **$^{13}$C NMR (DMSO, 150 MHz)**, δ (ppm): 164.03, 153.29, 151.58, 146.54, 143.46, 137.78, 136.45, 132.18, 126.86, 124.47, 122.37, 115.96, 105.55. **LCMS (ESI)**: Expected mass for $C_{13}H_8ClN_4O_3$ $(M + H)^+$: 303.03 Da, found 303.25 Da.

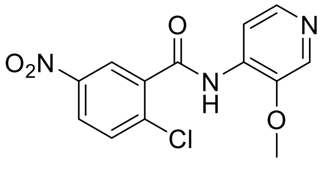

**SR33493**

**2-chloro-N-(3-methoxypyridin-4-yl)-5-nitrobenzamide**
$^{1}$**H NMR (DMSO-d6, 400 MHz)**, δ (ppm): 10.42 (s, 1H), 8.44 (d, *J*=2.7 Hz, 1H), 8.40 (br, 1H), 8.33 (dd, $J_1$=2.7Hz, $J_2$=8.8 Hz, 1H), 8.21 (m, 2H), 7.86 (d, *J*=8.9 Hz, 1H), 3.93 (s, 3H). $^{13}$**CNMR (DMSO, 100 MHz)**, δ (ppm): 164.49, 146.47, 143.40, 137.71, 137.52, 134.62, 134.04, 131.55, 126.19, 124.59, 57.00. **LCMS (ESI)**: Expected mass for $C_{13}H_{11}ClN_3O_4$ $(M + H)^+$: 308.04 Da, found 308.04 Da.

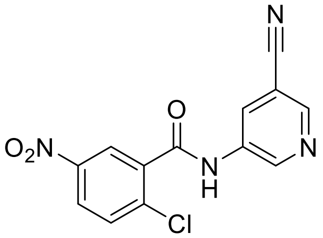

**SR33544**

**2-chloro-N-(5-cyanopyridin-3-yl)-5-nitrobenzamide**
$^{1}$**HNMR (DMSO-d6, 400 MHz)**, δ (ppm): 11.33 (s, 1H), 9.03 (d, *J*=2.5 Hz, 1H), 8.83 (d, *J*=2.5Hz, 1H), 8.59 (t, *J*=2.1 Hz, 2H), 8.39 (dd, $J_1$=2.9Hz, $J_2$=8.9 Hz, 1H), 7.94 (d, *J*=8.9 Hz, 1H). $^{13}$**CNMR (MeOH-d4, 100 MHz)**, δ (ppm): 199.47, 164.63, 147.18, 146.62, 144.27, 139.29, 137.53, 136.56, 131.95, 131.30, 129.64, 126.02, 125.87, 125.47, 123.80, 115.78, 37.48. **LCMS (ESI)**: Expected mass for $C_{13}H_8ClN_4O_3$ $(M + H)^+$: 303.03 Da, found 303.25 Da.

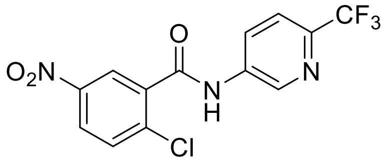

**SR35842**

**2-chloro-5-nitro-N-(6-(trifluoromethyl)pyridin-3-yl)benzamide**
$^{1}$**HNMR (DMSO-d6, 400 MHz)**, δ (ppm): 11.35 (s, 1H), 8.97 (d, *J*=2.82Hz, 1H), 8.44 (dd, $J_1$=2.07 Hz, $J_2$=8.46Hz), 8.39 (dd, $J_1$=2.07Hz, $J_2$=8.46Hz, 1H), 7.97 (d, *J*=8.96Hz, 1H), 7.94 (d, *J*=8.96Hz, 1H). $^{13}$**C NMR ($CDCl_3$, 100 MHz)**, δ (ppm): 164.21, 146.63, 141.74, 138.57, 137.55, 137.14, 131.98, 127.97, 126.72, 124.64, 121.92. **LCMS (ESI)**: Expected mass for $C_{13}H_8ClF_3N_3O_3$ $(M + H)^+$: 346.02 Da, found 346.02 Da.

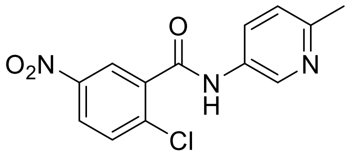

**SR36525**

**2-chloro-N-(6-methylpyridin-3-yl)-5-nitrobenzamide**
$^{1}$**HNMR (DMSO-d6, 400 MHz)**, δ (ppm): 10.85 (s, 1H), 8.71 ( d, *J*=2.4 Hz, 1H), 8.51 (d, *J*=2.7 Hz, 1H), 8.35 (dd, $J_1$=8.8 Hz, $J_2$=2.7 Hz, 1H), 8.02 (dd, $J_1$=8.4 Hz, $J_2$=2.7 Hz, 1H), 7.90 (d, *J*=8.9 Hz, 1H), 7.29 (d, *J*= 8.3Hz, 1H), 2.46 (s, 3H). $^{13}$**CNMR ($CDCl_3$, 100 MHz)**, δ (ppm): 163.52, 153.93, 146.61, 141.05, 137.73, 137.56, 133.20, 131.88, 127.97, 126.36, 124.44, 123.43, 23.90. **LCMS (ESI)**: Expected mass for $C_{13}H_{11}ClN_3O_3$ $(M + H)^+$: 292.05 Da, found 292.05 Da.

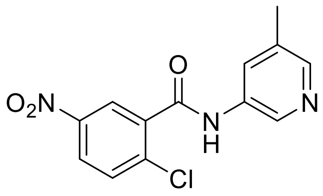

**SR36668**

**2-chloro-N-(5-methylpyridin-3-yl)-5-nitrobenzamide**

**$^1$HNMR (MeOD-d4, 400 MHz)**, δ (ppm): 9.33 (s, 1H), 8.56 (d, *J*=2.78Hz, 1H), 8.54 (br, 1H), 8.42 (d, *J*=2.78Hz, 1H), 8.40 (d, *J*=2.62Hz, 1H), 7.86 (d, *J*=8.83Hz, 1H), 2.60 (s, 3H). **$^{13}$CNMR (MeOD-d4, 100 MHz)**, δ (ppm): 164.55, 146.61, 137.56, 136.07, 135.14, 131.41, 130.61, 126.12, 123.87, 17.10. **LCMS (ESI)**: Expected mass for $C_{13}H_{11}ClN_3O_3$ $(M + H)^+$: 292.04 Da, found 292.04 Da.

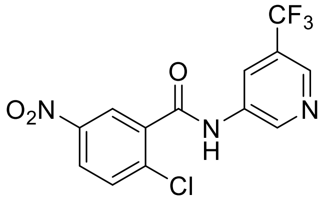

**SR36669**

**2-chloro-5-nitro-N-(5-(trifluoromethyl)pyridin-3-yl)benzamide**

**$^1$ HNMR (MeOD-d4, 400 MHz)**, δ (ppm): 9.03 (d, *J*=2.03Hz, 1H), 8.70 (s, 1H), 8.67 (t, *J*=1.85Hz, 1H), 8.56 (d, *J*=2.49Hz, 1H), 8.39 (dd, $J_1$=2.49Hz, $J_2$=8.78Hz, 1H), 7.85 (d, *J*=8.78Hz, 1H). **$^{13}$C NMR (MeOD-d4, 100 MHz)**, δ (ppm): 164.67, 146.63, 144.18, 141.11, 141.07, 137.55, 136.64, 135.68, 131.28, 126.88, 125.83, 123.79, 122.00. **LCMS (ESI)**: Expected mass for $C_{13}H_8ClF_3N_3O_3$ $(M +H)^+$: 346.02 Da, found 346.02 Da.

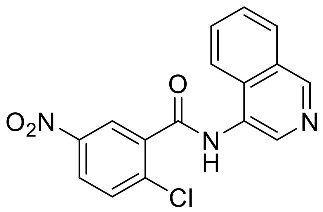

**SR36705**

**2-chloro-N-(isoquinolin-4-yl)-5-nitrobenzamide**

**$^1$HNMR (DMSO-d6, 400 MHz)**, δ (ppm): 11.08 (s, 1H), 8.91 (d, *J*=4.93 Hz, 1H), 8.64 (d, *J*=2.78 Hz, 1H), 8.39 (dd, $J_1$=2.78 Hz, $J_2$=8.64 Hz, 1H), 8.34 (dd, $J_1$=0.85 Hz, $J_2$= 8.78 Hz, 1H), 8.15 (d, *J*=4.58 Hz, 1H), 8.06 (dd, $J_1$= 0.62Hz, $J_2$= 8.40Hz, 1H), 7.94 (d, *J*= 8.77Hz, 1H), 7.81 (dt, $J_1$= 1.24Hz, $J_2$= 8.31Hz, 1H), 7.65 (dt, $J_1$= 1.24Hz, $J_2$= 8.31Hz, 1H). **$^{13}$CNMR ($CDCl_3$, 100 MHz)**, δ (ppm): 164.70, 151.30, 149.19, 146.68, 141.32, 137.88, 137.55, 131.72, 130.18, 129.87, 126.72, 126.40, 124.71, 123.13, 121.77, 113.96. **LCMS (ESI)**: Expected mass for $C_{16}H_{11}ClN_3O_3$ $(M + H)^+$: 328.05 Da, found 328.25 Da.

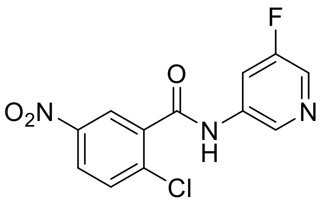

**SR36706**

**2-chloro-N-(5-fluoropyridin-3-yl)-5-nitrobenzamide**

**$^1$HNMR (DMSO-d6, 400 MHz)**, δ (ppm): 11.22 (br, 1H), 8.69 (br, 1H), 8.85 (d, *J*=2.70 Hz, 1H), 8.41 (br, 1H), 8.37 (dd, $J_1$=2.70 Hz, $J_2$=8.70 Hz, 1H), 8.14 (d, *J*=10.90 Hz, 1H), 7.92 (d, *J*= 8.66Hz, 1H). **$^{13}$C NMR ($CDCl_3$, 100 MHz)**, δ (ppm): 164.05, 146.63, 137.86, 137.54, 137.22, 131.97, 126.66, 124.57, 114.27, 114.05. **LCMS (ESI)**: Expected mass for $C_{12}H_8ClFN_3O_3$ $(M + H)^+$: 296.02 Da, found 296.02 Da.

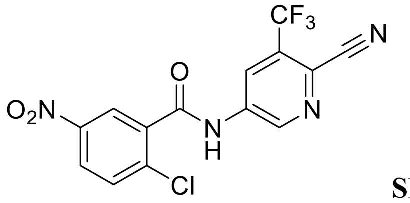


**SR36708**

**2-chloro-N-(6-cyano-5-(trifluoromethyl)pyridin-3-yl)-5-nitrobenzamide**

**$^{1}$HNMR (DMSO-d6, 400 MHz)**, δ (ppm): 11.78 (s, 1H), 9.14 (d, *J*=1.84 Hz, 1H), 8.76 (d, *J*=2.12 Hz, 1H), 8.64 (d, *J*=2.74 Hz, 1H), 8.41 (dd, $J_1$=2.87 Hz, $J_2$= 8.78 Hz, 1H), 7.95 (d, *J*=8.86 Hz, 1H). **$^{13}$CNMR ($CDCl_3$, 100 MHz)**, δ (ppm): 164.60, 146.61, 145.15, 138.91, 137.60, 136.38, 132.15, 129.80, 127.14, 124.84, 124.21, 123.79, 115.21. **LCMS (ESI)**: Expected mass for $C_{14}H_7ClF_3N_4O_3$ $(M + H)^+$: 371.02 Da, found 371.25 Da.

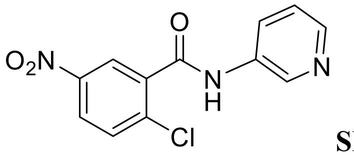


**SR1000392694**

**2-chloro-5-nitro-N-(pyridin-3-yl)benzamide**

**$^{1}$HNMR ($CDCl_3$, 400 MHz)**, δ (ppm): 8.67 (d, *J*=2.7 Hz, 1H), 8.63 (d, *J*=2.7 Hz, 1H), 8.44 (d, *J*=4.5 Hz, 1H), 8.29 (dd, $J_1$=3.2Hz, $J_2$=9.1Hz, 2H), 8.16 (br, 1H), 7.69 (d, *J*=8.6 Hz, 1H), 7.37 (d, *J*=3.2Hz, 2H). **$^{13}$CNMR ($CDCl_3$, 100 MHz)**, δ (ppm): 163.72, 146.63, 145.65, 141.79, 137.65, 137.55, 135.67, 131.90, 127.33, 126.44, 124.48, 124.29. **LCMS (ESI)**: Expected mass for $C_{12}H_9ClN_3O_3$ $(M + H)^+$: 278.03 Da, found 278.03 Da.

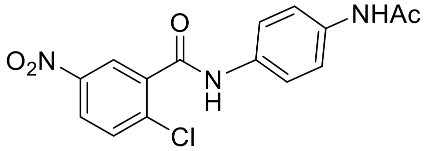


**SR1000401841**

**N-(4-acetamidophenyl)-2-chloro-5-nitrobenzamide**

**$^{1}$HNMR (DMSO-d6, 400 MHz)**, δ (ppm): 10.65 (br, 1H), 9.96 (br, 1H), 8.42 (d, *J*=2.55 Hz, 1H), 8.33 (dd, $J_1$=2.55Hz, $J_2$=8.92 Hz, 1H), 7.87 (d, *J*=8.92Hz, 1H), 7.55-7.62 (m, 4H), 2.03 (s, 3H). **$^{13}$CNMR (DMSO-d6, 100 MHz)**, δ (ppm): 168.65, 162.96, 146.60, 138.21, 137.57, 136.14, 134.11, 131.80, 126.11, 124.30, 120.69, 119.86, 24.35. **LCMS (ESI)**: Expected mass for $C_{15}H_{13}ClN_3O_4$ $(M + H)^+$: 334.06 Da, found 334.23 Da.

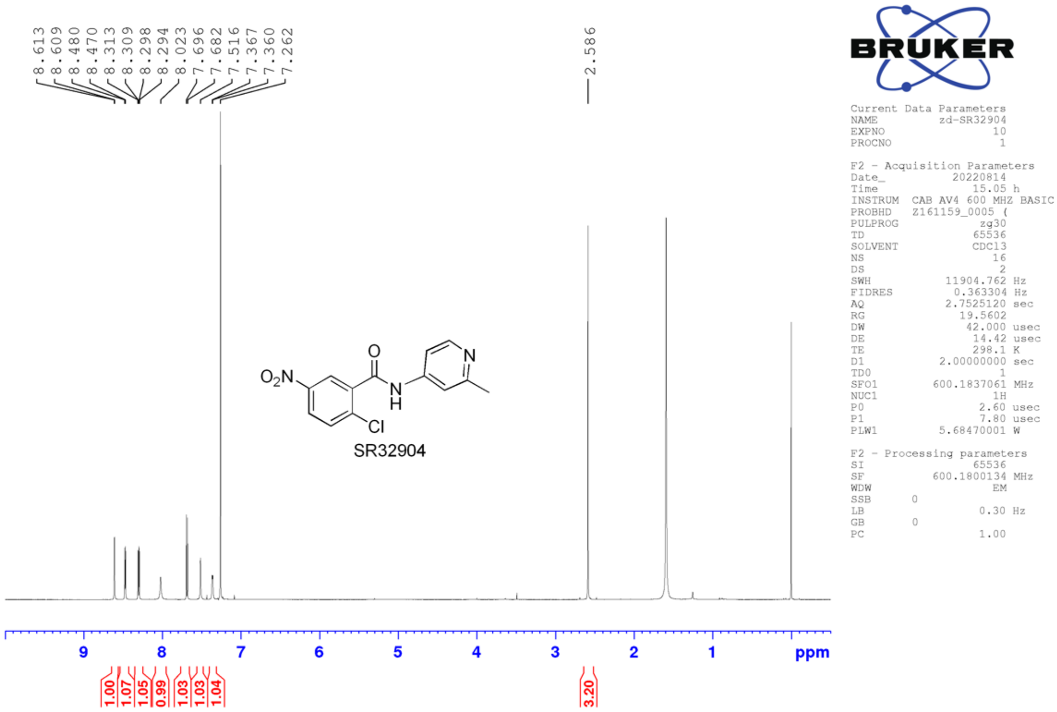

8.613
8.609
8.480
8.470
8.313
8.309
8.298
8.294
8.023
7.696
7.682
7.516
7.367
7.360
7.262
2.586
$O_2N$
O
N
H
N
Cl
SR32904
9
8
7
6
5
4
3
2
1
ppm
1.00
1.07
1.05
0.99
1.03
1.03
1.04
3.20
BRUKER
Current Data Parameters
NAME zd-SR32904
EXPNO 10
PROCNO 1
F2 - Acquisition Parameters
Date_ 20220814
Time 15.05 h
INSTRUM CAB AV4 600 MHZ BASIC
PROBHD Z161159_0005 (
PULPROG zg30
TD 65536
SOLVENT CDCl3
NS 16
DS 2
SWH 11904.762 Hz
FIDRES 0.363304 Hz
AQ 2.7525120 sec
RG 19.5602
DW 42.000 usec
DE 14.42 usec
TE 298.1 K
D1 2.00000000 sec
TD0 1
SFO1 600.1837061 MHz
NUC1 1H
P0 2.60 usec
P1 7.80 usec
PLW1 5.68470001 W
F2 - Processing parameters
SI 65536
SF 600.1800134 MHz
WDW EM
SSB 0
LB 0.30 Hz
GB 0
PC 1.00


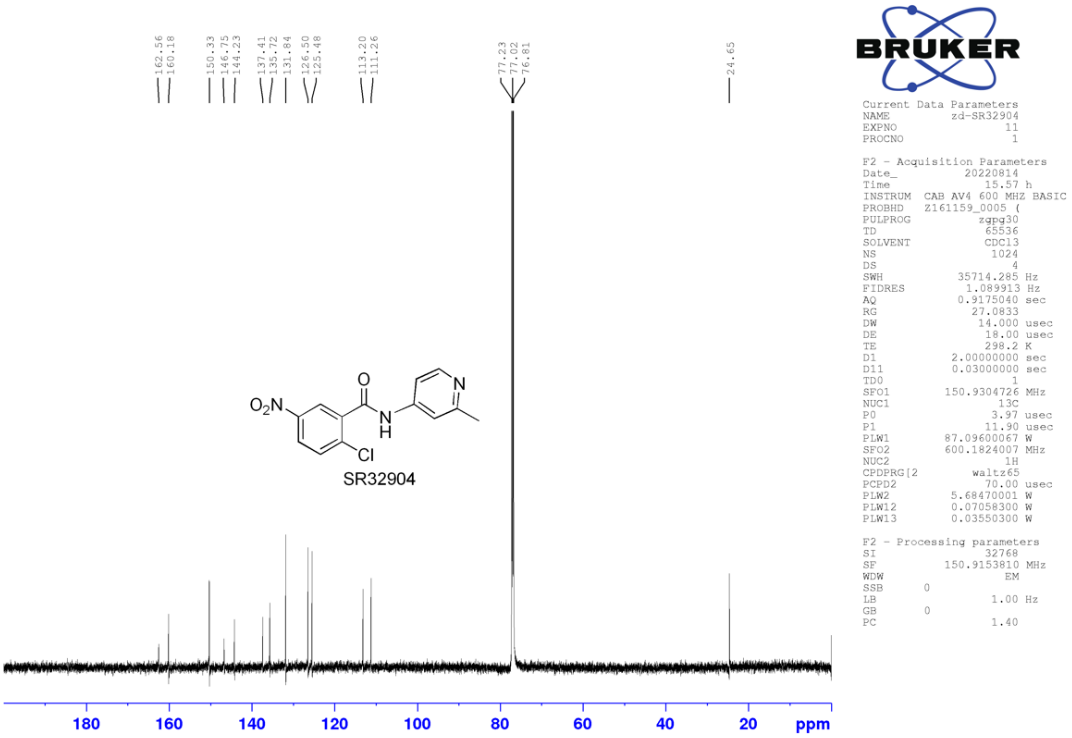

162.56
160.18
150.33
146.75
144.23
137.41
135.72
131.84
126.50
125.48
113.20
111.26
77.23
77.02
76.81
24.65
$O_2N$
O
N
H
N
Cl
SR32904
180
160
140
120
100
80
60
40
20
ppm
BRUKER
Current Data Parameters
NAME zd-SR32904
EXPNO 11
PROCNO 1
F2 - Acquisition Parameters
Date_ 20220814
Time 15.57 h
INSTRUM CAB AV4 600 MHZ BASIC
PROBHD Z161159_0005 (
PULPROG zgpg30
TD 65536
SOLVENT CDCl3
NS 1024
DS 4
SWH 35714.285 Hz
FIDRES 1.089913 Hz
AQ 0.9175040 sec
RG 27.0833
DW 14.000 usec
DE 18.00 usec
TE 298.2 K
D1 2.00000000 sec
D11 0.03000000 sec
TD0 1
SFO1 150.9304726 MHz
NUC1 13C
P0 3.97 usec
P1 11.90 usec
PLW1 87.09600067 W
SFO2 600.1824007 MHz
NUC2 1H
CPDPRG[2 waltz65
PCPD2 70.00 usec
PLW2 5.68470001 W
PLW12 0.07058300 W
PLW13 0.03550300 W
F2 - Processing parameters
SI 32768
SF 150.9153810 MHz
WDW EM
SSB 0
LB 1.00 Hz
GB 0
PC 1.40

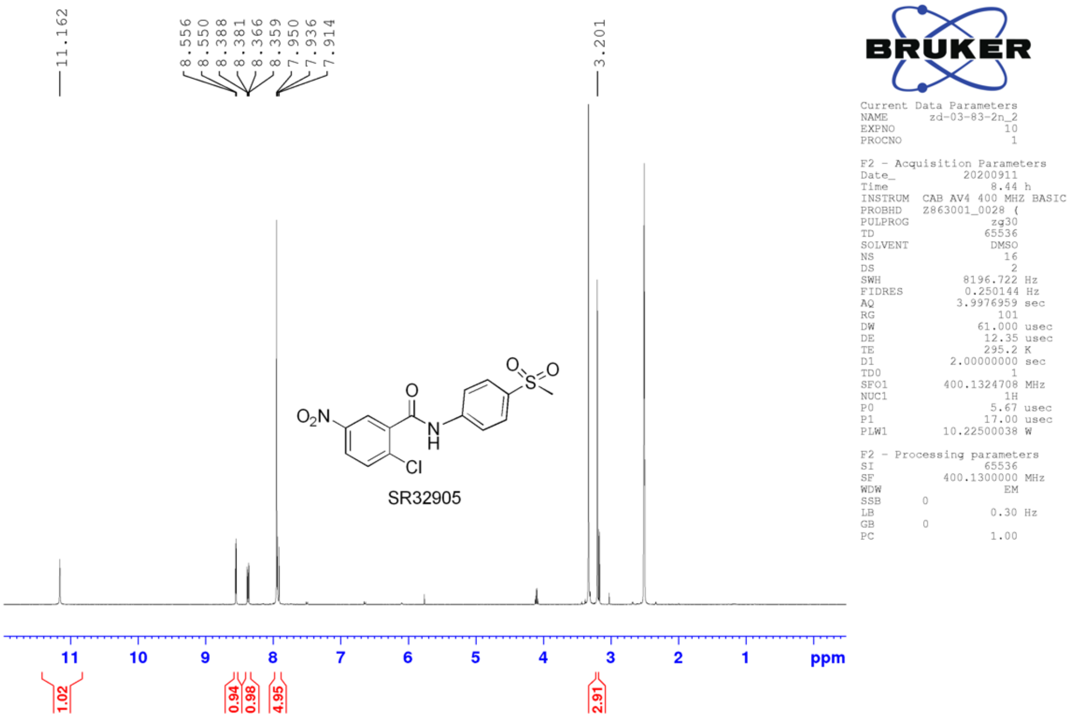
11.162
8.556
8.550
8.388
8.381
8.366
8.359
7.950
7.936
7.914
3.201
SR32905
11 10 9 8 7 6 5 4 3 2 1 ppm
1.02
0.94
0.98
4.95
2.91
BRUKER
Current Data Parameters
NAME zd-03-83-2n_2
EXPNO 10
PROCNO 1
F2 - Acquisition Parameters
Date_ 20200911
Time 8.44 h
INSTRUM CAB AV4 400 MHZ BASIC
PROBHD Z863001_0028 (
PULPROG zg30
TD 65536
SOLVENT DMSO
NS 16
DS 2
SWH 8196.722 Hz
FIDRES 0.250144 Hz
AQ 3.9976959 sec
RG 101
DW 61.000 usec
DE 12.35 usec
TE 295.2 K
D1 2.00000000 sec
TD0 1
SFO1 400.1324708 MHz
NUC1 1H
P0 5.67 usec
P1 17.00 usec
PLW1 10.22500038 W
F2 - Processing parameters
SI 65536
SF 400.1300000 MHz
WDW EM
SSB 0
LB 0.30 Hz
GB 0
PC 1.00

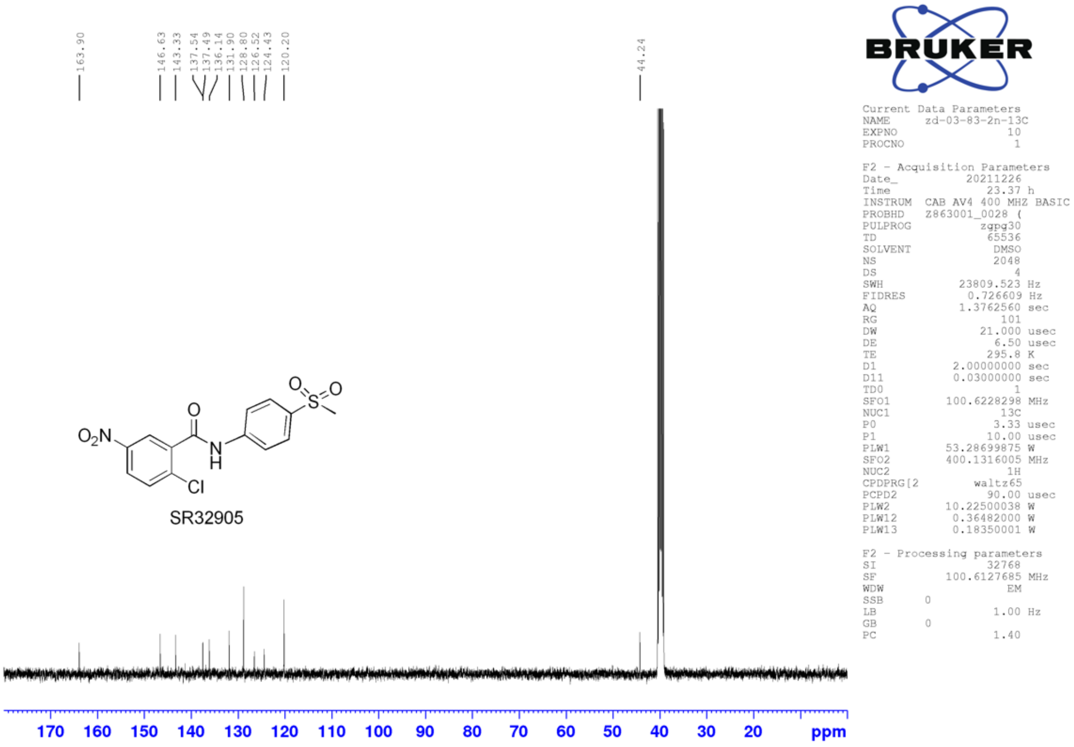
163.90
146.63
143.33
137.54
137.49
136.14
131.90
128.80
126.52
124.43
120.20
44.24
SR32905
170 160 150 140 130 120 110 100 90 80 70 60 50 40 30 20 ppm
BRUKER
Current Data Parameters
NAME zd-03-83-2n-13C
EXPNO 10
PROCNO 1
F2 - Acquisition Parameters
Date_ 20211226
Time 23.37 h
INSTRUM CAB AV4 400 MHZ BASIC
PROBHD Z863001_0028 (
PULPROG zgpg30
TD 65536
SOLVENT DMSO
NS 2048
DS 4
SWH 23809.523 Hz
FIDRES 0.726609 Hz
AQ 1.3762560 sec
RG 101
DW 21.000 usec
DE 6.50 usec
TE 295.8 K
D1 2.00000000 sec
D11 0.03000000 sec
TD0 1
SFO1 100.6228298 MHz
NUC1 13C
P0 3.33 usec
P1 10.00 usec
PLW1 53.28699875 W
SFO2 400.1316005 MHz
NUC2 1H
CPDPRG[2 waltz65
PCPD2 90.00 usec
PLW2 10.22500038 W
PLW12 0.36482000 W
PLW13 0.18350001 W
F2 - Processing parameters
SI 32768
SF 100.6127685 MHz
WDW EM
SSB 0
LB 1.00 Hz
GB 0
PC 1.40

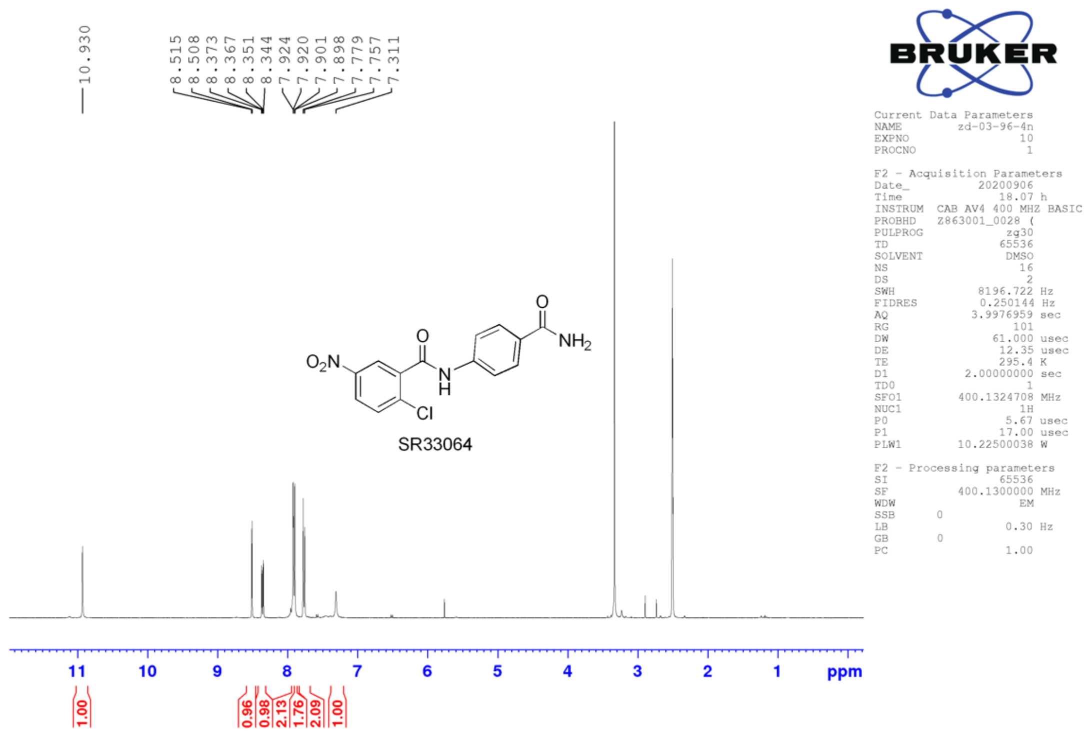
10.930
8.515
8.508
8.373
8.367
8.351
8.344
7.924
7.920
7.901
7.898
7.779
7.757
7.311
SR33064
11
10
9
8
7
6
5
4
3
2
1
ppm
1.00
0.96
0.98
2.13
1.76
2.09
1.00
BRUKER
Current Data Parameters
NAME zd-03-96-4n
EXPNO 10
PROCNO 1
F2 - Acquisition Parameters
Date_ 20200906
Time 18.07 h
INSTRUM CAB AV4 400 MHZ BASIC
PROBHD Z863001_0028 (
PULPROG zg30
TD 65536
SOLVENT DMSO
NS 16
DS 2
SWH 8196.722 Hz
FIDRES 0.250144 Hz
AQ 3.9976959 sec
RG 101
DW 61.000 usec
DE 12.35 usec
TE 295.4 K
D1 2.00000000 sec
TD0 1
SFO1 400.1324708 MHz
NUC1 1H
P0 5.67 usec
P1 17.00 usec
PLW1 10.22500038 W
F2 - Processing parameters
SI 65536
SF 400.1300000 MHz
WDW EM
SSB 0
LB 0.30 Hz
GB 0
PC 1.00

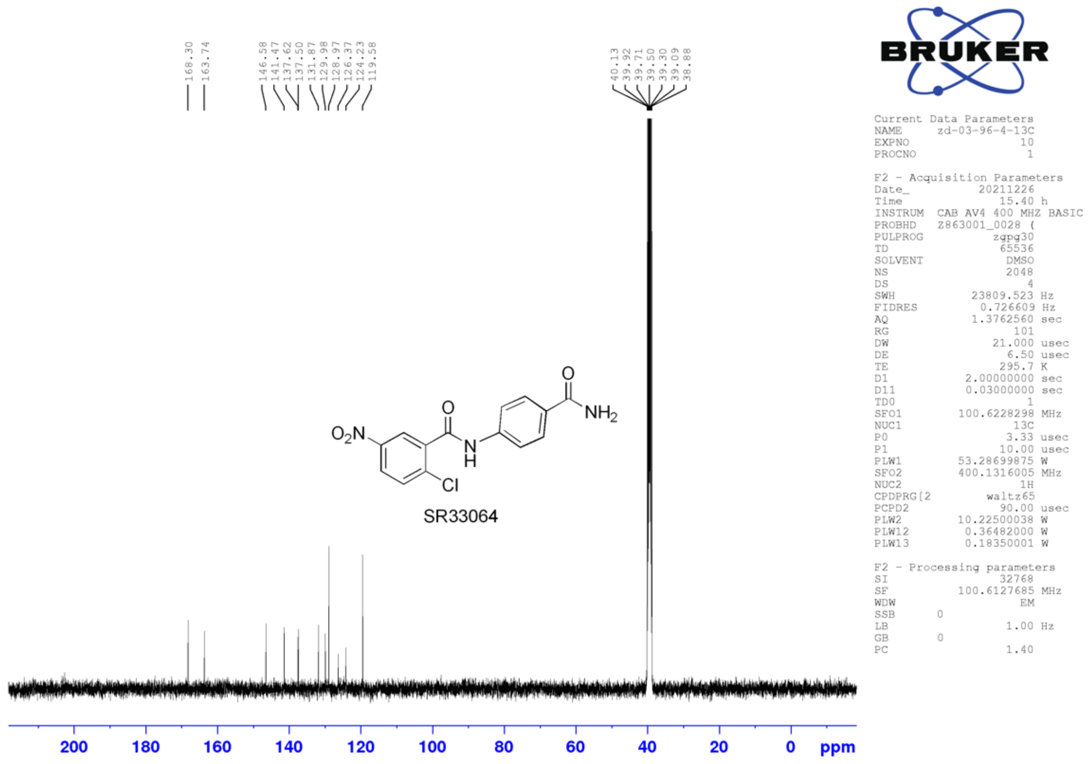
168.30
163.74
146.58
141.47
137.62
137.50
131.87
129.98
128.97
126.37
124.23
119.58
40.13
39.92
39.71
39.50
39.30
39.09
38.88
SR33064
200
180
160
140
120
100
80
60
40
20
0
ppm
BRUKER
Current Data Parameters
NAME zd-03-96-4-13C
EXPNO 10
PROCNO 1
F2 - Acquisition Parameters
Date_ 20211226
Time 15.40 h
INSTRUM CAB AV4 400 MHZ BASIC
PROBHD Z863001_0028 (
PULPROG zgpg30
TD 65536
SOLVENT DMSO
NS 2048
DS 4
SWH 23809.523 Hz
FIDRES 0.726609 Hz
AQ 1.3762560 sec
RG 101
DW 21.000 usec
DE 6.50 usec
TE 295.7 K
D1 2.00000000 sec
D11 0.03000000 sec
TD0 1
SFO1 100.6228298 MHz
NUC1 13C
P0 3.33 usec
P1 10.00 usec
PLW1 53.28699875 W
SFO2 400.1316005 MHz
NUC2 1H
CPDPRG[2 waltz65
PCPD2 90.00 usec
PLW2 10.22500038 W
PLW12 0.36482000 W
PLW13 0.18350001 W
F2 - Processing parameters
SI 32768
SF 100.6127685 MHz
WDW EM
SSB 0
LB 1.00 Hz
GB 0
PC 1.40

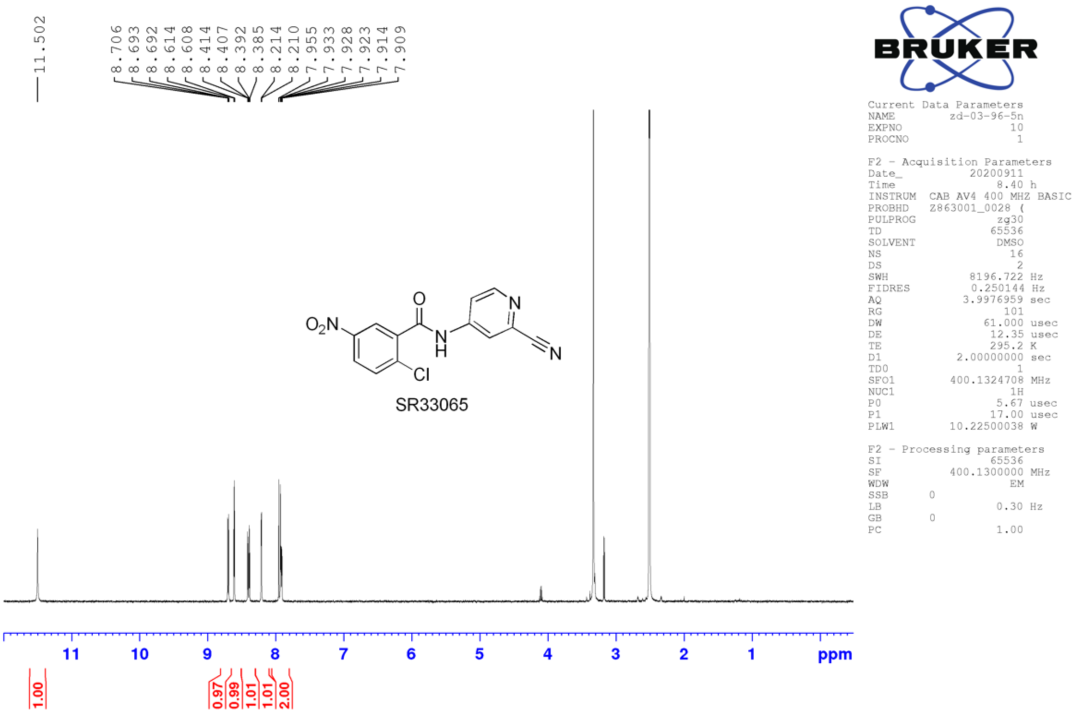
11.502
8.706
8.693
8.692
8.614
8.608
8.414
8.407
8.392
8.385
8.214
8.210
7.955
7.933
7.928
7.923
7.914
7.909
O2N
O
N
H
N
N
Cl
SR33065
BRUKER
Current Data Parameters
NAME zd-03-96-5n
EXPNO 10
PROCNO 1
F2 - Acquisition Parameters
Date_ 20200911
Time 8.40 h
INSTRUM CAB AV4 400 MHZ BASIC
PROBHD Z863001_0028 (
PULPROG zg30
TD 65536
SOLVENT DMSO
NS 16
DS 2
SWH 8196.722 Hz
FIDRES 0.250144 Hz
AQ 3.9976959 sec
RG 101
DW 61.000 usec
DE 12.35 usec
TE 295.2 K
D1 2.00000000 sec
TD0 1
SFO1 400.1324708 MHz
NUC1 1H
P0 5.67 usec
P1 17.00 usec
PLW1 10.22500038 W
F2 - Processing parameters
SI 65536
SF 400.1300000 MHz
WDW EM
SSB 0
LB 0.30 Hz
GB 0
PC 1.00
11 10 9 8 7 6 5 4 3 2 1 ppm
1.00
0.97
0.99
1.01
1.01
2.00

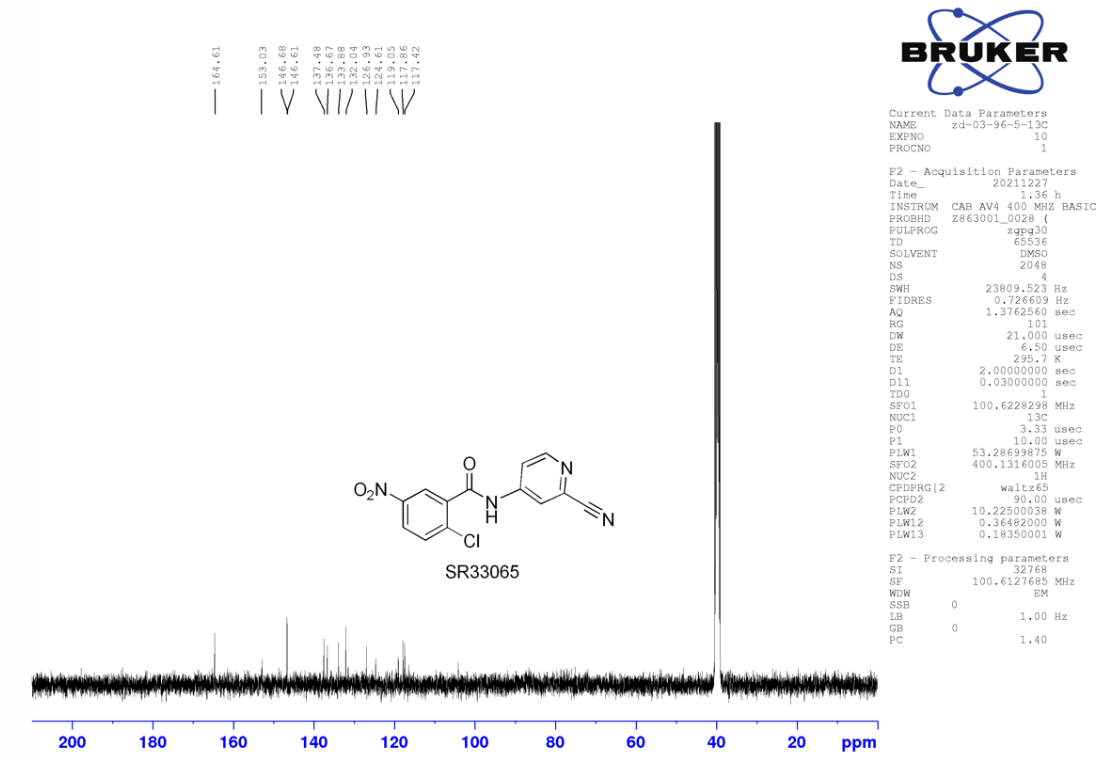
164.61
153.03
146.68
146.61
137.48
136.67
133.88
132.04
126.93
124.61
119.05
117.86
117.42
O2N
O
N
H
N
N
Cl
SR33065
BRUKER
Current Data Parameters
NAME zd-03-96-5-13C
EXPNO 10
PROCNO 1
F2 - Acquisition Parameters
Date_ 20211227
Time 1.36 h
INSTRUM CAB AV4 400 MHZ BASIC
PROBHD Z863001_0028 (
PULPROG zgpg30
TD 65536
SOLVENT DMSO
NS 2048
DS 4
SWH 23809.523 Hz
FIDRES 0.726609 Hz
AQ 1.3762560 sec
RG 101
DW 21.000 usec
DE 6.50 usec
TE 295.7 K
D1 2.00000000 sec
D11 0.03000000 sec
TD0 1
SFO1 100.6228298 MHz
NUC1 13C
P0 3.33 usec
P1 10.00 usec
PLW1 53.28699875 W
SFO2 400.1316005 MHz
NUC2 1H
CPDPRG[2 waltz65
PCPD2 90.00 usec
PLW2 10.22500038 W
PLW12 0.36482000 W
PLW13 0.18350001 W
F2 - Processing parameters
SI 32768
SF 100.6127685 MHz
WDW EM
SSB 0
LB 1.00 Hz
GB 0
PC 1.40
200 180 160 140 120 100 80 60 40 20 ppm

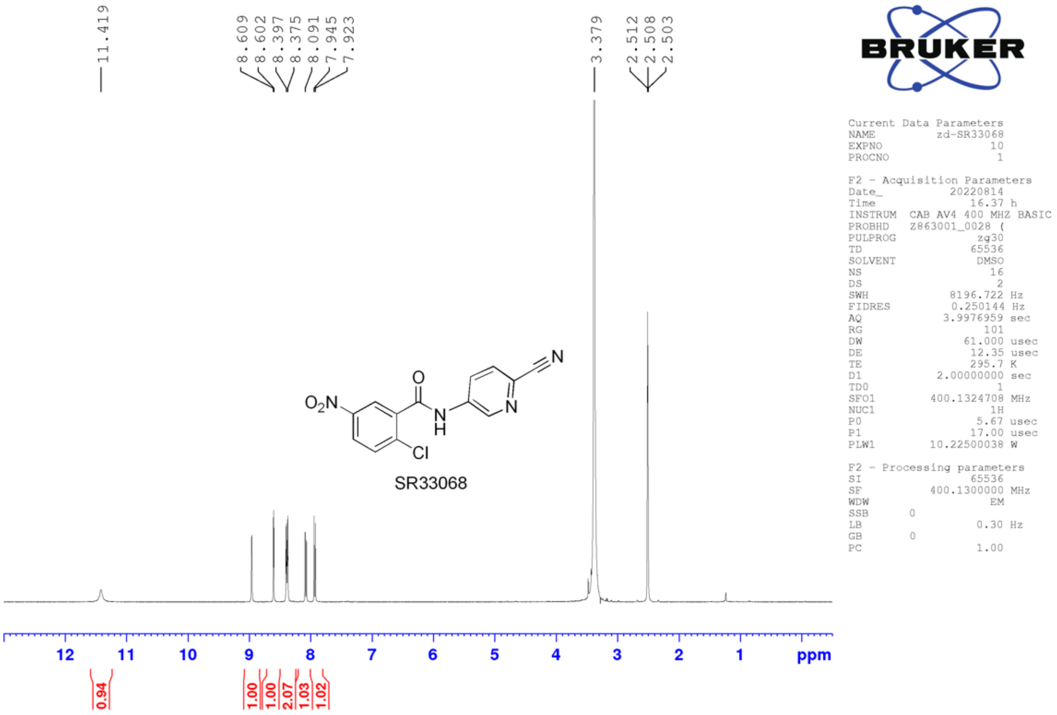
11.419
8.609
8.602
8.397
8.375
8.091
7.945
7.923
3.379
2.512
2.508
2.503
SR33068
BRUKER
Current Data Parameters
NAME zd-SR33068
EXPNO 10
PROCNO 1
F2 - Acquisition Parameters
Date_ 20220814
Time 16.37 h
INSTRUM CAB AV4 400 MHZ BASIC
PROBHD Z863001_0028 (
PULPROG zg30
TD 65536
SOLVENT DMSO
NS 16
DS 2
SWH 8196.722 Hz
FIDRES 0.250144 Hz
AQ 3.9976959 sec
RG 101
DW 61.000 usec
DE 12.35 usec
TE 295.7 K
D1 2.00000000 sec
TD0 1
SFO1 400.1324708 MHz
NUC1 1H
P0 5.67 usec
P1 17.00 usec
PLW1 10.22500038 W
F2 - Processing parameters
SI 65536
SF 400.1300000 MHz
WDW EM
SSB 0
LB 0.30 Hz
GB 0
PC 1.00
12 11 10 9 8 7 6 5 4 3 2 1 ppm
0.94
1.00
1.00
2.07
1.03
1.02

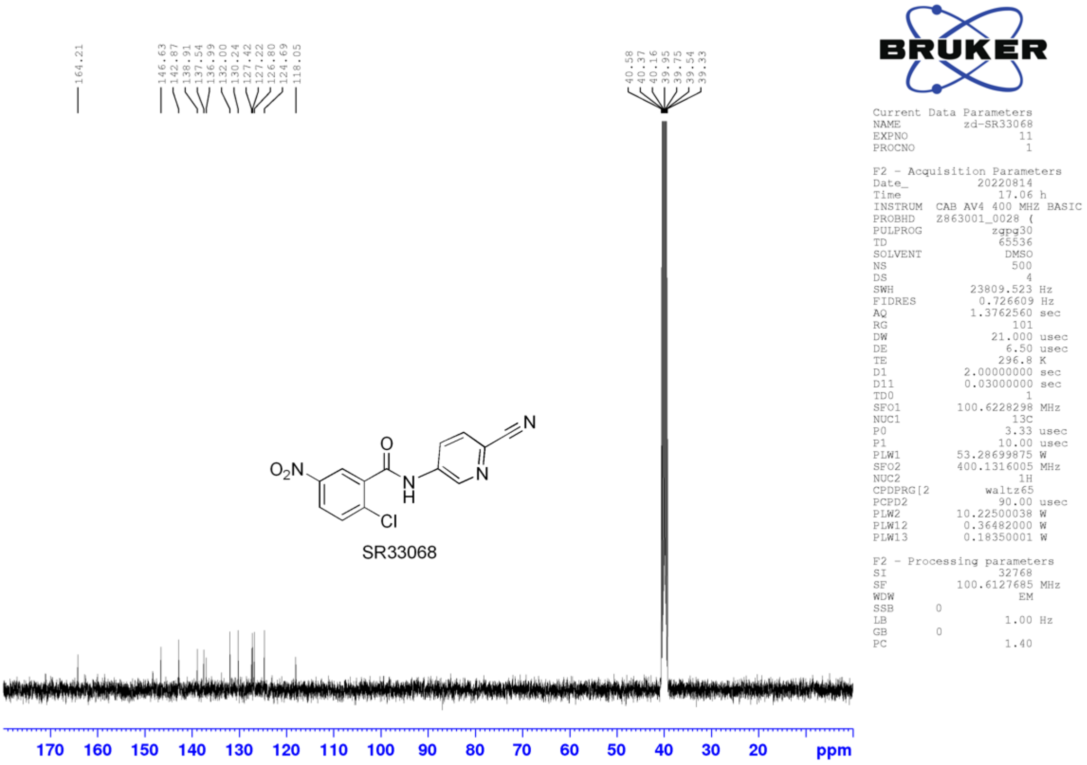
164.21
146.63
142.87
138.91
137.54
136.99
132.00
130.24
127.42
127.22
126.80
124.69
118.05
40.58
40.37
40.16
39.95
39.75
39.54
39.33
SR33068
BRUKER
Current Data Parameters
NAME zd-SR33068
EXPNO 11
PROCNO 1
F2 - Acquisition Parameters
Date_ 20220814
Time 17.06 h
INSTRUM CAB AV4 400 MHZ BASIC
PROBHD Z863001_0028 (
PULPROG zgpg30
TD 65536
SOLVENT DMSO
NS 500
DS 4
SWH 23809.523 Hz
FIDRES 0.726609 Hz
AQ 1.3762560 sec
RG 101
DW 21.000 usec
DE 6.50 usec
TE 296.8 K
D1 2.00000000 sec
D11 0.03000000 sec
TD0 1
SFO1 100.6228298 MHz
NUC1 13C
P0 3.33 usec
P1 10.00 usec
PLW1 53.28699875 W
SFO2 400.1316005 MHz
NUC2 1H
CPDPRG[2 waltz65
PCPD2 90.00 usec
PLW2 10.22500038 W
PLW12 0.36482000 W
PLW13 0.18350001 W
F2 - Processing parameters
SI 32768
SF 100.6127685 MHz
WDW EM
SSB 0
LB 1.00 Hz
GB 0
PC 1.40
170 160 150 140 130 120 110 100 90 80 70 60 50 40 30 20 ppm

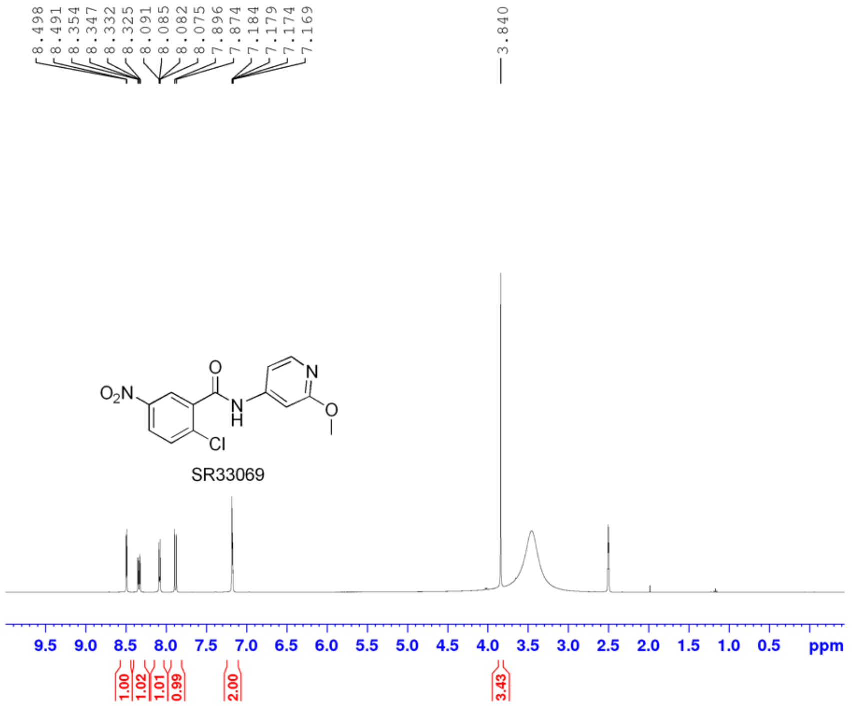


```
Current Data Parameters
NAME          zd-SR33069
EXPNO                 10
PROCNO                 1

F2 - Acquisition Parameters
Date_           20220831
Time              10.29 h
INSTRUM  CAB AV4 400 MHZ BASIC
PROBHD   Z863001_0028 (
PULPROG             zg30
TD                 65536
SOLVENT             DMSO
NS                    16
DS                     2
SWH             8196.722 Hz
FIDRES          0.250144 Hz
AQ             3.9976959 sec
RG                   101
DW                61.000 usec
DE                 12.35 usec
TE                 295.3 K
D1            2.00000000 sec
TD0                    1
SFO1         400.1324708 MHz
NUC1                  1H
P0                  5.67 usec
P1                 17.00 usec
PLW1         10.22500038 W

F2 - Processing parameters
SI                 65536
SF           400.1300030 MHz
WDW                   EM
SSB      0
LB                  0.30 Hz
GB       0
PC                  1.00
```

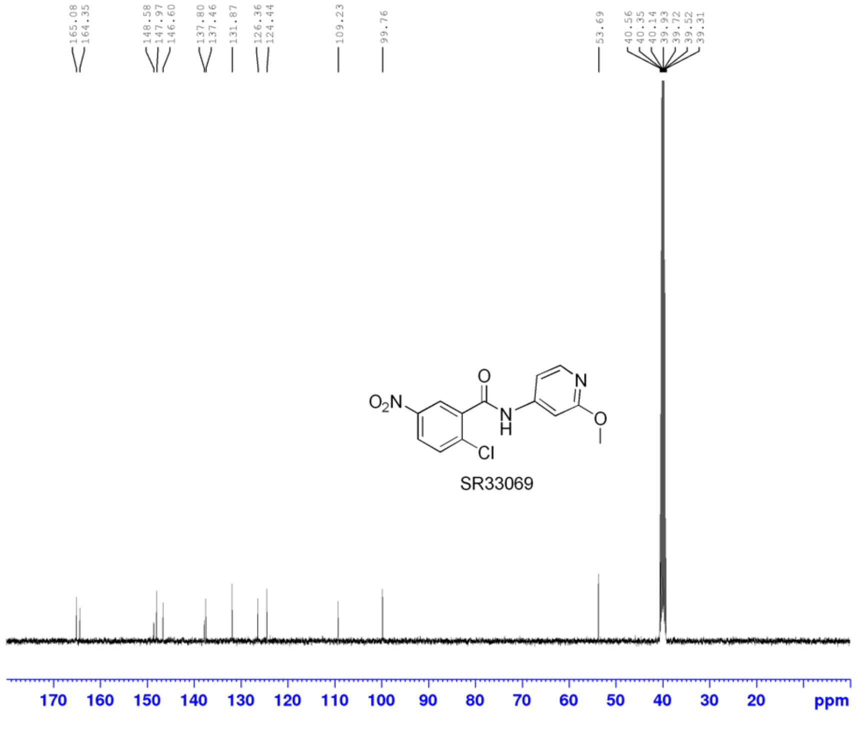


```
Current Data Parameters
NAME          zd-SR33069
EXPNO                 11
PROCNO                 1

F2 - Acquisition Parameters
Date_           20220831
Time              10.53 h
INSTRUM  CAB AV4 400 MHZ BASIC
PROBHD   Z863001_0028 (
PULPROG           zgpg30
TD                 65536
SOLVENT             DMSO
NS                   400
DS                     4
SWH            23809.523 Hz
FIDRES          0.726609 Hz
AQ             1.3762560 sec
RG                   101
DW                21.000 usec
DE                  6.50 usec
TE                 296.4 K
D1            2.00000000 sec
D11           0.03000000 sec
TD0                    1
SFO1         100.6228298 MHz
NUC1                 13C
P0                  3.33 usec
P1                 10.00 usec
PLW1         53.28699875 W
SFO2         400.1316005 MHz
NUC2                  1H
CPDPRG[2         waltz65
PCPD2              90.00 usec
PLW2         10.22500038 W
PLW12         0.36482000 W
PLW13         0.18350001 W

F2 - Processing parameters
SI                 32768
SF           100.6127685 MHz
WDW                   EM
SSB      0
LB                  1.00 Hz
GB       0
PC                  1.40
```

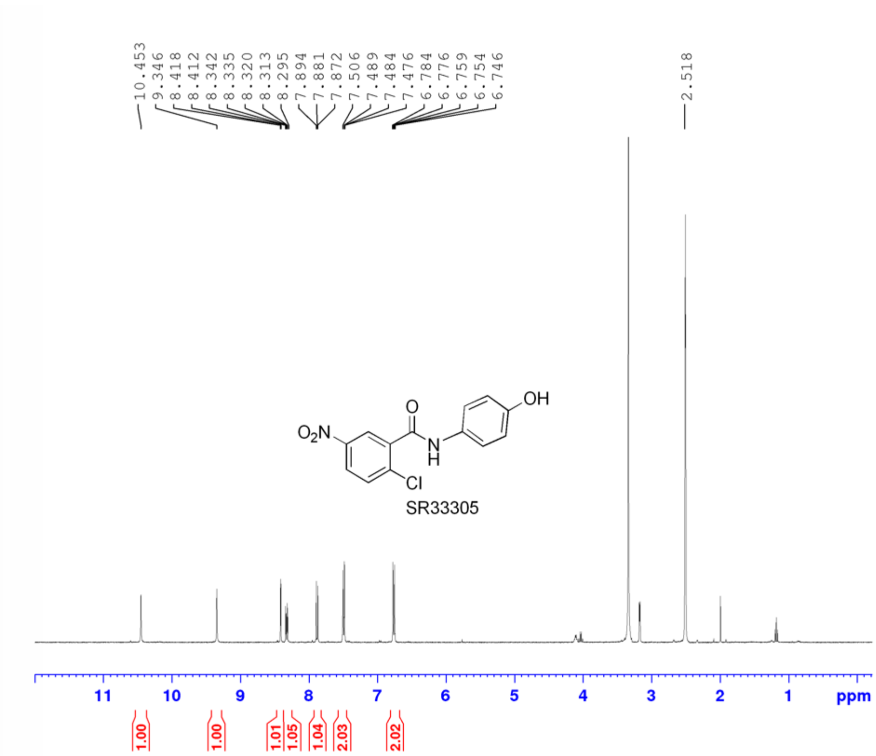


Current Data Parameters
NAME zd-03-99-2n
EXPNO 10
PROCNO 1

F2 - Acquisition Parameters
Date_ 20200921
Time 11.09
INSTRUM spect
PROBHD 5 mm PAQNP 13C
PULPROG zg30
TD 58188
SOLVENT DMSO
NS 16
DS 2
SWH 7183.908 Hz
FIDRES 0.123460 Hz
AQ 4.0498848 sec
RG 574.7
DW 69.600 usec
DE 6.00 usec
TE 296.1 K
D1 1.50000000 sec
TD0 1

======== CHANNEL f1 ========
NUC1 1H
P1 15.00 usec
PL1 0 dB
PL1W 9.31909847 W
SFO1 400.1324710 MHz

F2 - Processing parameters
SI 32768
SF 400.1300000 MHz
WDW EM
SSB 0
LB 0.30 Hz
GB 0
PC 1.00

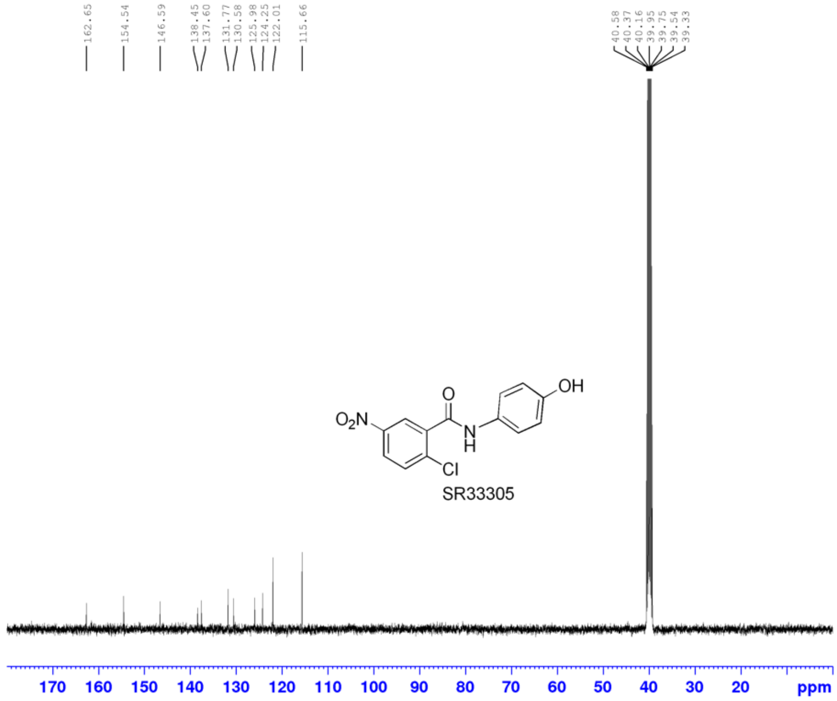


Current Data Parameters
NAME zd-SR33305
EXPNO 11
PROCNO 1

F2 - Acquisition Parameters
Date_ 20220814
Time 15.59 h
INSTRUM CAB AV4 400 MHZ BASIC
PROBHD Z863001_0028 (
PULPROG zgpg30
TD 65536
SOLVENT DMSO
NS 512
DS 4
SWH 23809.523 Hz
FIDRES 0.726609 Hz
AQ 1.3762560 sec
RG 101
DW 21.000 usec
DE 6.50 usec
TE 296.7 K
D1 2.00000000 sec
D11 0.03000000 sec
TD0 1
SFO1 100.6228298 MHz
NUC1 13C
P0 3.33 usec
P1 10.00 usec
PLW1 53.28699875 W
SFO2 400.1316005 MHz
NUC2 1H
CPDPRG[2 waltz65
PCPD2 90.00 usec
PLW2 10.22500038 W
PLW12 0.36482000 W
PLW13 0.18350001 W

F2 - Processing parameters
SI 32768
SF 100.6127685 MHz
WDW EM
SSB 0
LB 1.00 Hz
GB 0
PC 1.40

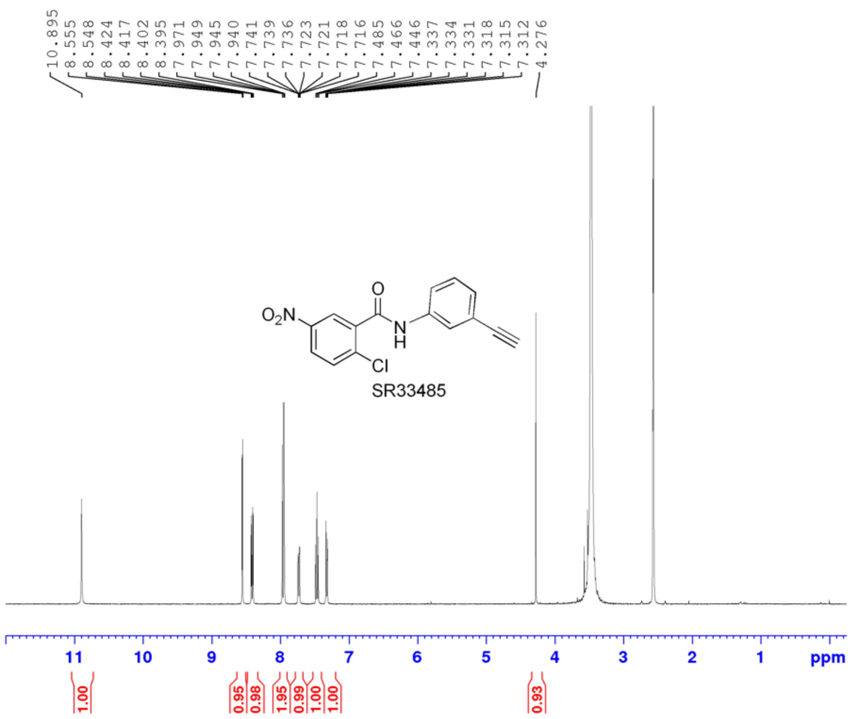


```
Current Data Parameters
NAME          zd-SR33485-
EXPNO                  10
PROCNO                  1

F2 - Acquisition Parameters
Date_            20220830
Time                22.42 h
INSTRUM   CAB AV4 400 MHZ BASIC
PROBHD    Z863001_0028 (
PULPROG              zg30
TD                  65536
SOLVENT              DMSO
NS                     16
DS                      2
SWH              8196.722 Hz
FIDRES           0.250144 Hz
AQ              3.9976959 sec
RG                    101
DW                 61.000 usec
DE                  12.35 usec
TE                  295.3 K
D1             2.00000000 sec
TD0                     1
SFO1          400.1324708 MHz
NUC1                   1H
P0                   5.67 usec
P1                  17.00 usec
PLW1          10.22500038 W

F2 - Processing parameters
SI                  65536
SF            400.1299771 MHz
WDW                    EM
SSB      0
LB                   0.30 Hz
GB       0
PC                   1.00
```

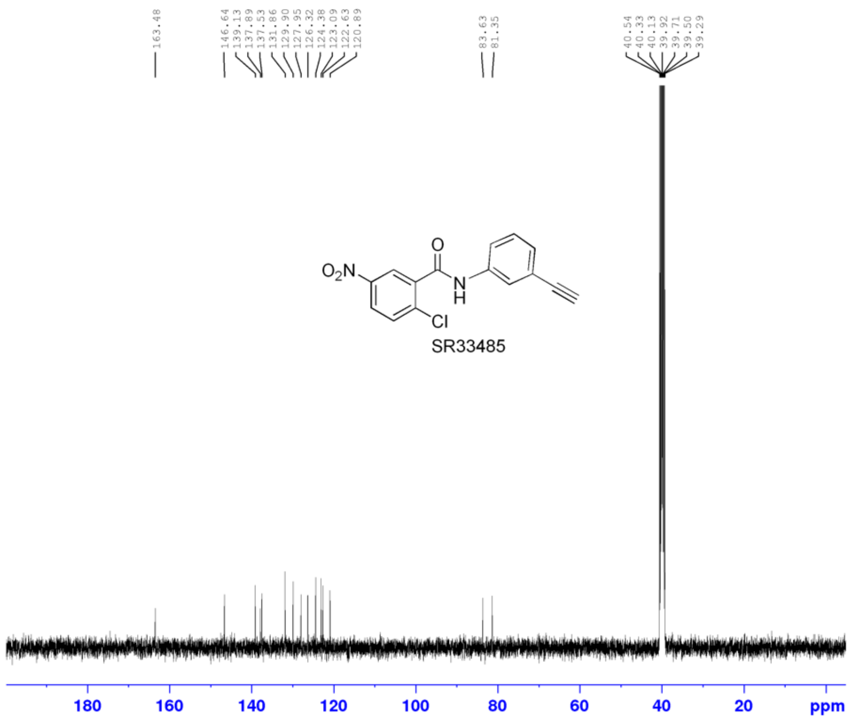


```
Current Data Parameters
NAME          zd-SR33485-
EXPNO                  11
PROCNO                  1

F2 - Acquisition Parameters
Date_            20220830
Time                23.42 h
INSTRUM   CAB AV4 400 MHZ BASIC
PROBHD    Z863001_0028 (
PULPROG            zgpg30
TD                  65536
SOLVENT              DMSO
NS                   1024
DS                      4
SWH             23809.523 Hz
FIDRES           0.726609 Hz
AQ              1.3762560 sec
RG                    101
DW                 21.000 usec
DE                   6.50 usec
TE                  296.3 K
D1             2.00000000 sec
D11            0.03000000 sec
TD0                     1
SFO1          100.6228298 MHz
NUC1                  13C
P0                   3.33 usec
P1                  10.00 usec
PLW1          53.28699875 W
SFO2          400.1316005 MHz
NUC2                   1H
CPDPRG[2          waltz65
PCPD2               90.00 usec
PLW2          10.22500038 W
PLW12          0.36482000 W
PLW13          0.18350001 W

F2 - Processing parameters
SI                  32768
SF            100.6127685 MHz
WDW                    EM
SSB      0
LB                   1.00 Hz
GB       0
PC                   1.40
```

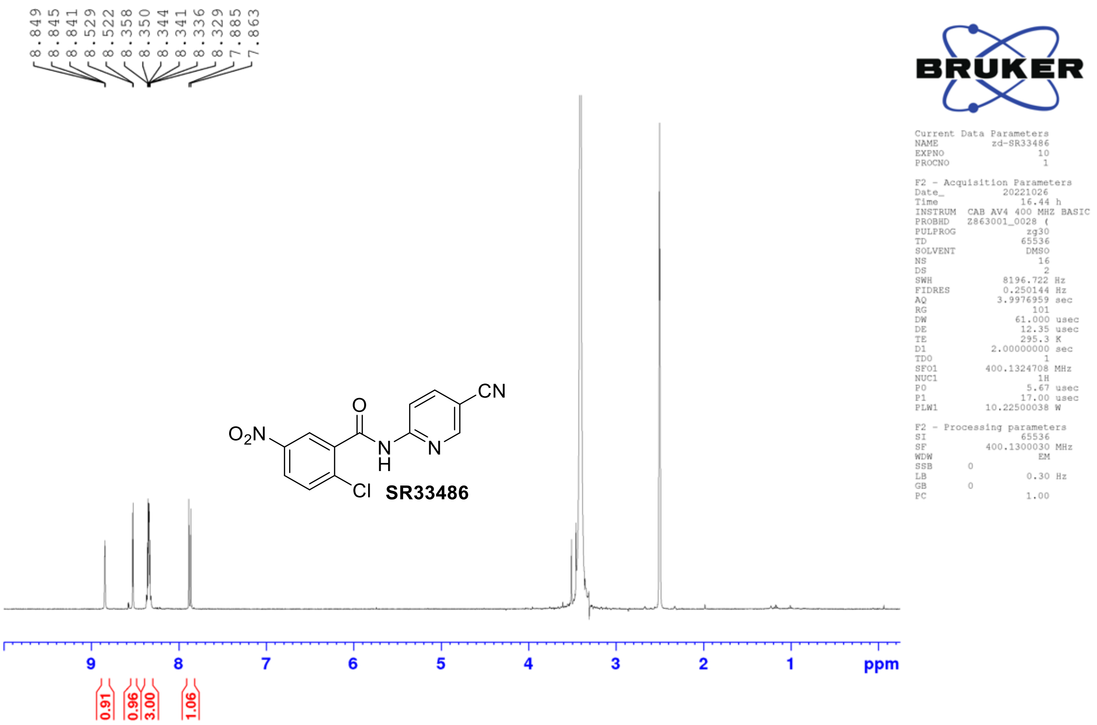

8.849
8.845
8.841
8.529
8.522
8.358
8.350
8.344
8.341
8.336
8.329
7.885
7.863
BRUKER
Current Data Parameters
NAME zd-SR33486
EXPNO 10
PROCNO 1
F2 - Acquisition Parameters
Date_ 20221026
Time 16.44 h
INSTRUM CAB AV4 400 MHZ BASIC
PROBHD Z863001_0028 (
PULPROG zg30
TD 65536
SOLVENT DMSO
NS 16
DS 2
SWH 8196.722 Hz
FIDRES 0.250144 Hz
AQ 3.9976959 sec
RG 101
DW 61.000 usec
DE 12.35 usec
TE 295.3 K
D1 2.00000000 sec
TD0 1
SFO1 400.1324708 MHz
NUC1 1H
P0 5.67 usec
P1 17.00 usec
PLW1 10.22500038 W
F2 - Processing parameters
SI 65536
SF 400.1300030 MHz
WDW EM
SSB 0
LB 0.30 Hz
GB 0
PC 1.00
O2N
O
N
H
N
CN
Cl
SR33486
9
8
7
6
5
4
3
2
1
ppm
0.91
0.96
3.00
1.06


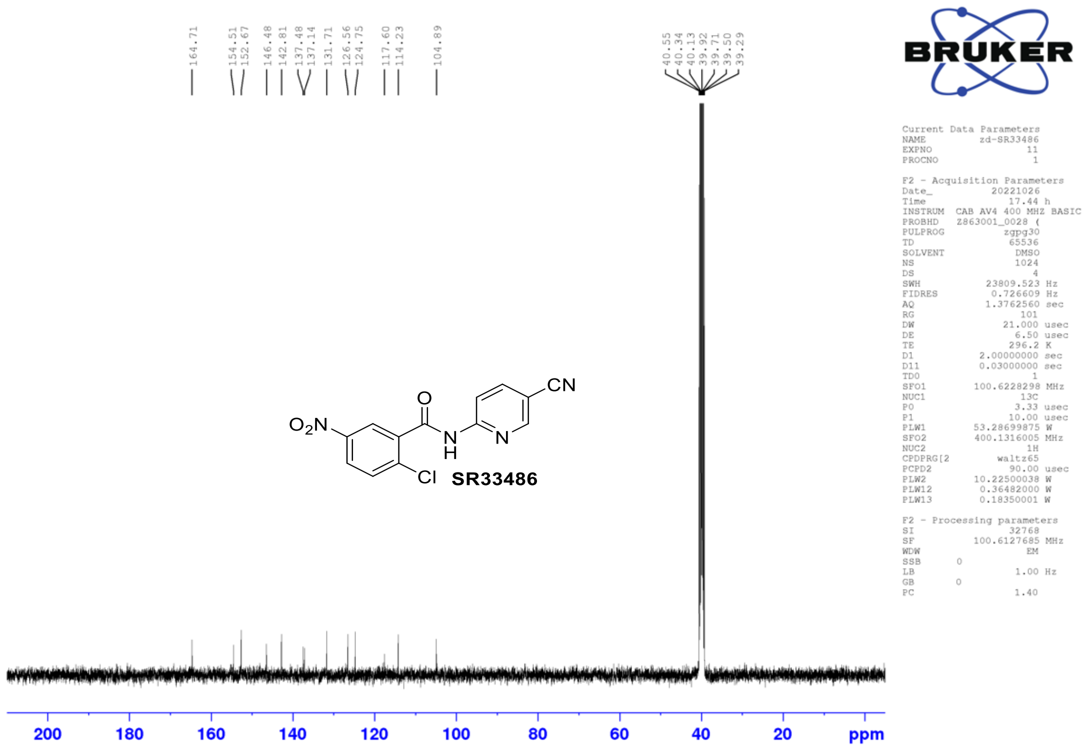

164.71
154.51
152.67
146.48
142.81
137.48
137.14
131.71
126.56
124.75
117.60
114.23
104.89
40.55
40.34
40.13
39.92
39.71
39.50
39.29
BRUKER
Current Data Parameters
NAME zd-SR33486
EXPNO 11
PROCNO 1
F2 - Acquisition Parameters
Date_ 20221026
Time 17.44 h
INSTRUM CAB AV4 400 MHZ BASIC
PROBHD Z863001_0028 (
PULPROG zgpg30
TD 65536
SOLVENT DMSO
NS 1024
DS 4
SWH 23809.523 Hz
FIDRES 0.726609 Hz
AQ 1.3762560 sec
RG 101
DW 21.000 usec
DE 6.50 usec
TE 296.2 K
D1 2.00000000 sec
D11 0.03000000 sec
TD0 1
SFO1 100.6228298 MHz
NUC1 13C
P0 3.33 usec
P1 10.00 usec
PLW1 53.28699875 W
SFO2 400.1316005 MHz
NUC2 1H
CPDPRG[2 waltz65
PCPD2 90.00 usec
PLW2 10.22500038 W
PLW12 0.36482000 W
PLW13 0.18350001 W
F2 - Processing parameters
SI 32768
SF 100.6127685 MHz
WDW EM
SSB 0
LB 1.00 Hz
GB 0
PC 1.40
O2N
O
N
H
N
CN
Cl
SR33486
200
180
160
140
120
100
80
60
40
20
ppm

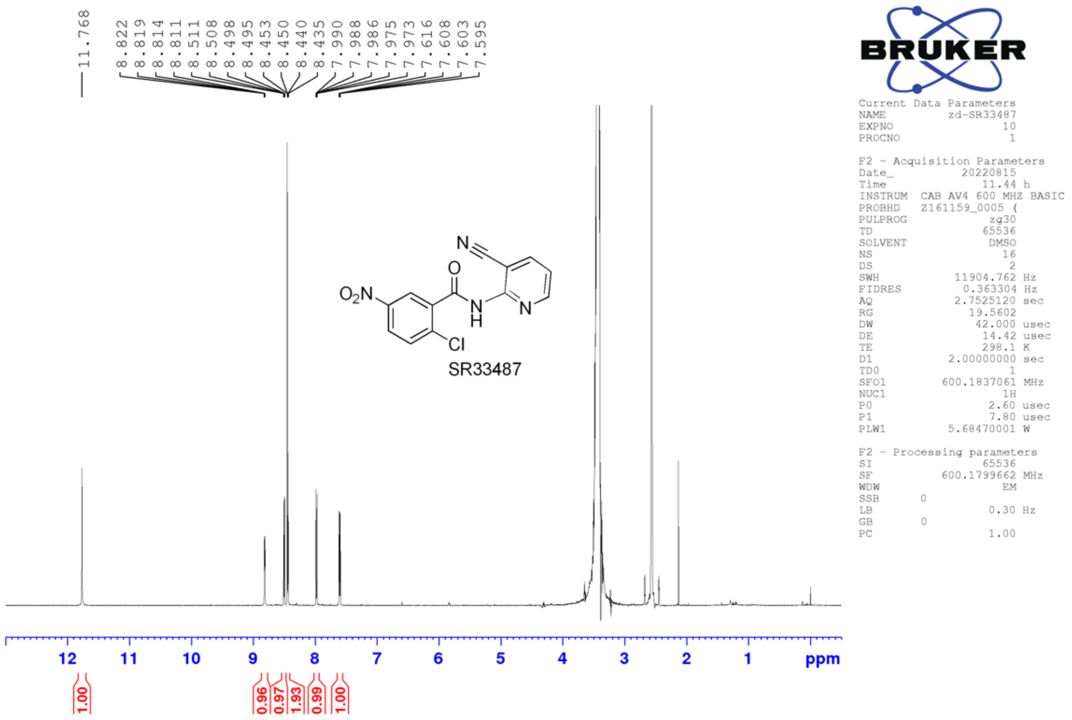
11.768
8.822 8.819 8.814 8.811 8.511 8.508 8.498 8.495 8.453 8.450 8.440 8.435 7.990 7.988 7.986 7.975 7.973 7.616 7.608 7.603 7.595
O2N
O
N
H
N
Cl
SR33487
12 11 10 9 8 7 6 5 4 3 2 1 ppm
1.00 0.96 0.97 1.93 0.99 1.00
BRUKER
Current Data Parameters
NAME zd-SR33487
EXPNO 10
PROCNO 1
F2 - Acquisition Parameters
Date_ 20220815
Time 11.44 h
INSTRUM CAB AV4 600 MHZ BASIC
PROBHD Z161159_0005 (
PULPROG zg30
TD 65536
SOLVENT DMSO
NS 16
DS 2
SWH 11904.762 Hz
FIDRES 0.363304 Hz
AQ 2.7525120 sec
RG 19.5602
DW 42.000 usec
DE 14.42 usec
TE 298.1 K
D1 2.00000000 sec
TD0 1
SFO1 600.1837061 MHz
NUC1 1H
P0 2.60 usec
P1 7.80 usec
PLW1 5.68470001 W
F2 - Processing parameters
SI 65536
SF 600.1799662 MHz
WDW EM
SSB 0
LB 0.30 Hz
GB 0
PC 1.00

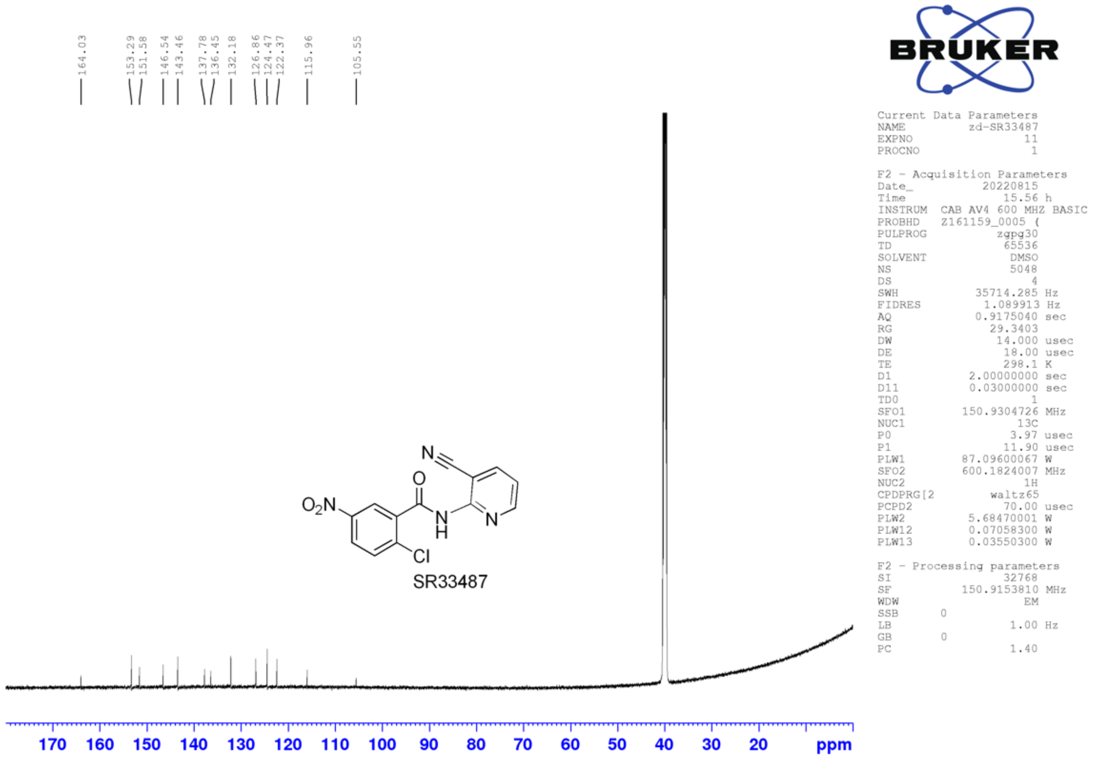
164.03 153.29 151.58 146.54 143.46 137.78 136.45 132.18 126.86 124.47 122.37 115.96 105.55
O2N
O
N
H
N
Cl
SR33487
170 160 150 140 130 120 110 100 90 80 70 60 50 40 30 20 ppm
BRUKER
Current Data Parameters
NAME zd-SR33487
EXPNO 11
PROCNO 1
F2 - Acquisition Parameters
Date_ 20220815
Time 15.56 h
INSTRUM CAB AV4 600 MHZ BASIC
PROBHD Z161159_0005 (
PULPROG zgpg30
TD 65536
SOLVENT DMSO
NS 5048
DS 4
SWH 35714.285 Hz
FIDRES 1.089913 Hz
AQ 0.9175040 sec
RG 29.3403
DW 14.000 usec
DE 18.00 usec
TE 298.1 K
D1 2.00000000 sec
D11 0.03000000 sec
TD0 1
SFO1 150.9304726 MHz
NUC1 13C
P0 3.97 usec
P1 11.90 usec
PLW1 87.09600067 W
SFO2 600.1824007 MHz
NUC2 1H
CPDPRG[2 waltz65
PCPD2 70.00 usec
PLW2 5.68470001 W
PLW12 0.07058300 W
PLW13 0.03550300 W
F2 - Processing parameters
SI 32768
SF 150.9153810 MHz
WDW EM
SSB 0
LB 1.00 Hz
GB 0
PC 1.40

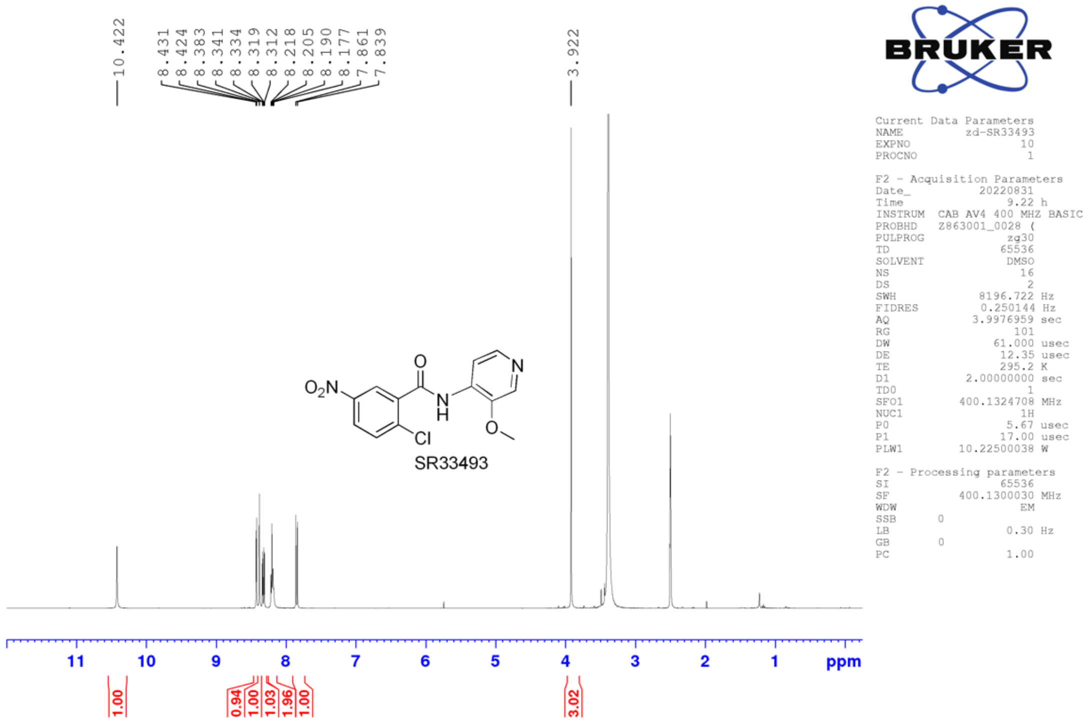
10.422
8.431
8.424
8.383
8.341
8.334
8.319
8.312
8.218
8.205
8.190
8.177
7.861
7.839
3.922
$O_2N$
O
N
N
H
Cl
O
SR33493
11 10 9 8 7 6 5 4 3 2 1 ppm
1.00
0.94
1.00
1.03
1.96
1.00
3.02
Current Data Parameters
NAME zd-SR33493
EXPNO 10
PROCNO 1
F2 - Acquisition Parameters
Date_ 20220831
Time 9.22 h
INSTRUM CAB AV4 400 MHZ BASIC
PROBHD Z863001_0028 (
PULPROG zg30
TD 65536
SOLVENT DMSO
NS 16
DS 2
SWH 8196.722 Hz
FIDRES 0.250144 Hz
AQ 3.9976959 sec
RG 101
DW 61.000 usec
DE 12.35 usec
TE 295.2 K
D1 2.00000000 sec
TD0 1
SFO1 400.1324708 MHz
NUC1 1H
P0 5.67 usec
P1 17.00 usec
PLW1 10.22500038 W
F2 - Processing parameters
SI 65536
SF 400.1300030 MHz
WDW EM
SSB 0
LB 0.30 Hz
GB 0
PC 1.00

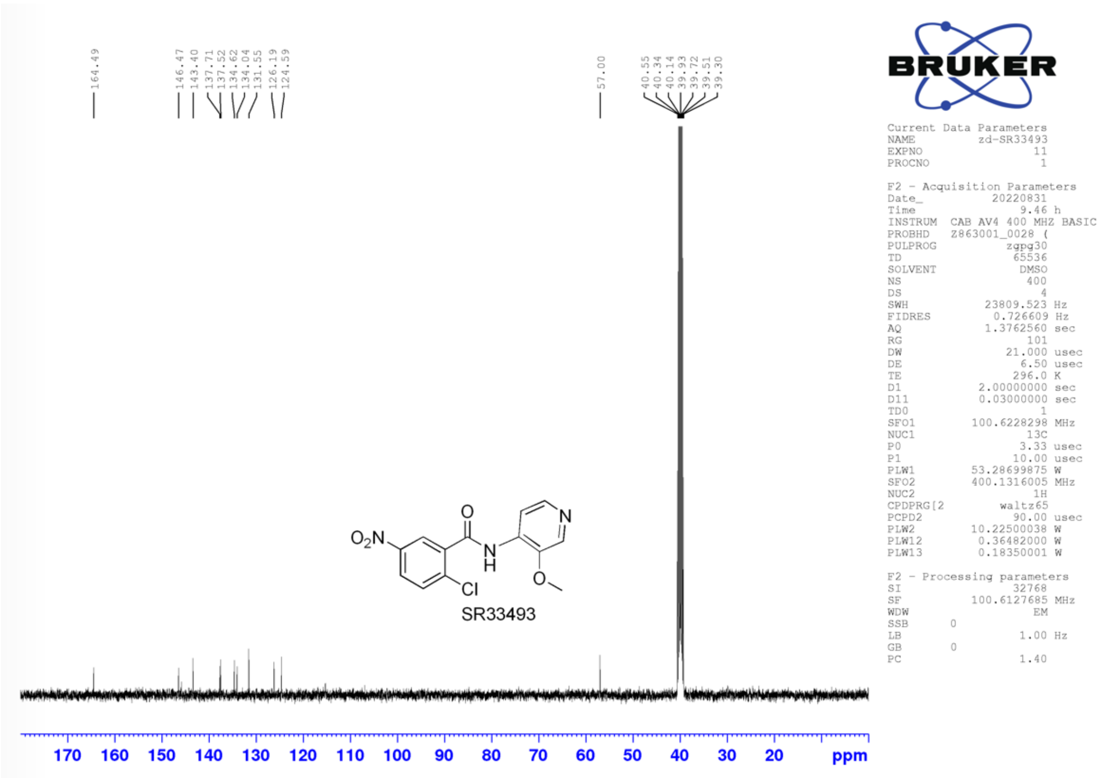
164.49
146.47
143.40
137.71
137.52
134.62
134.04
131.55
126.19
124.59
57.00
40.55
40.34
40.14
39.93
39.72
39.51
39.30
$O_2N$
O
N
N
H
Cl
O
SR33493
170 160 150 140 130 120 110 100 90 80 70 60 50 40 30 20 ppm
Current Data Parameters
NAME zd-SR33493
EXPNO 11
PROCNO 1
F2 - Acquisition Parameters
Date_ 20220831
Time 9.46 h
INSTRUM CAB AV4 400 MHZ BASIC
PROBHD Z863001_0028 (
PULPROG zgpg30
TD 65536
SOLVENT DMSO
NS 400
DS 4
SWH 23809.523 Hz
FIDRES 0.726609 Hz
AQ 1.3762560 sec
RG 101
DW 21.000 usec
DE 6.50 usec
TE 296.0 K
D1 2.00000000 sec
D11 0.03000000 sec
TD0 1
SFO1 100.6228298 MHz
NUC1 13C
P0 3.33 usec
P1 10.00 usec
PLW1 53.28699875 W
SFO2 400.1316005 MHz
NUC2 1H
CPDPRG[2 waltz65
PCPD2 90.00 usec
PLW2 10.22500038 W
PLW12 0.36482000 W
PLW13 0.18350001 W
F2 - Processing parameters
SI 32768
SF 100.6127685 MHz
WDW EM
SSB 0
LB 1.00 Hz
GB 0
PC 1.40

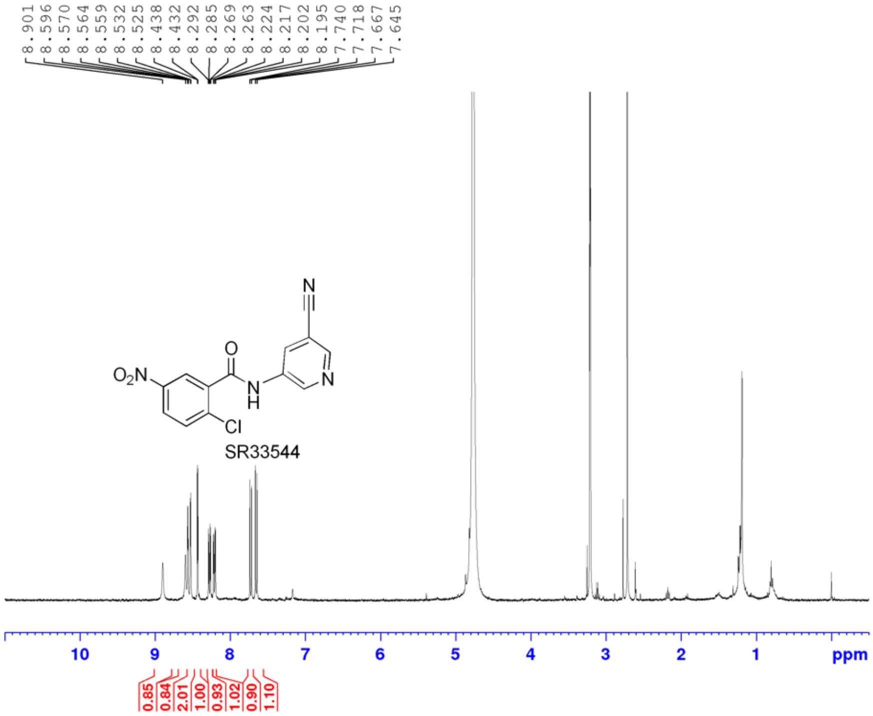


```
Current Data Parameters
NAME          zd-SR33544
EXPNO                 10
PROCNO                 1

F2 - Acquisition Parameters
Date_           20220831
Time               11.35 h
INSTRUM   CAB AV4 400 MHZ BASIC
PROBHD    Z863001_0028 (
PULPROG             zg30
TD                 65536
SOLVENT              MeOD
NS                    16
DS                     2
SWH             8196.722 Hz
FIDRES          0.250144 Hz
AQ             3.9976959 sec
RG                   101
DW                61.000 usec
DE                 12.35 usec
TE                 295.3 K
D1            2.00000000 sec
TD0                    1
SFO1         400.1324708 MHz
NUC1                  1H
P0                  5.67 usec
P1                 17.00 usec
PLW1         10.22500038 W

F2 - Processing parameters
SI                 65536
SF           400.1300476 MHz
WDW                   EM
SSB      0
LB                  0.30 Hz
GB       0
PC                  1.00
```

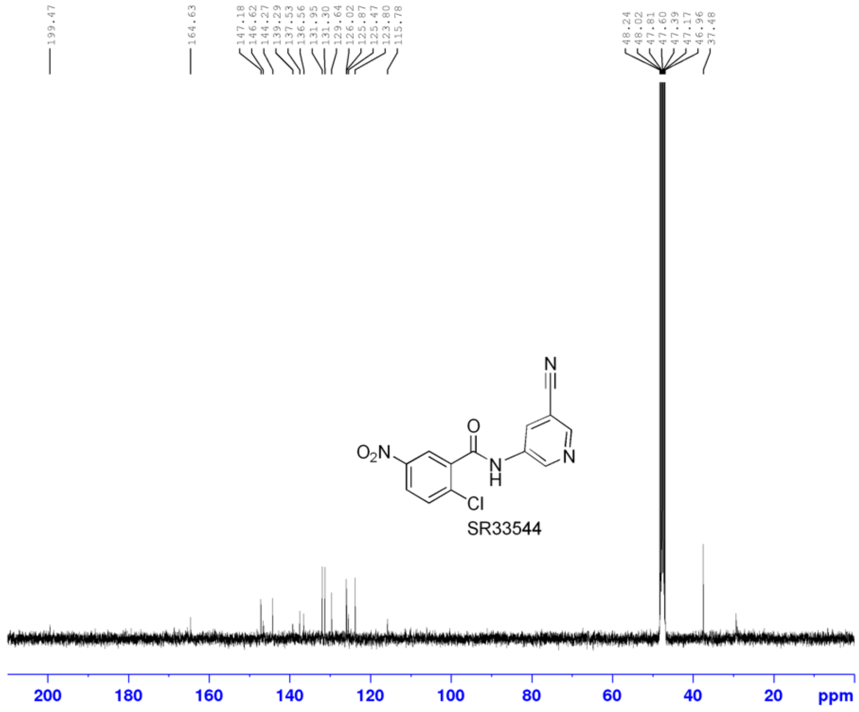


```
Current Data Parameters
NAME          zd-SR33544
EXPNO                 21
PROCNO                 1

F2 - Acquisition Parameters
Date_           20220831
Time               19.54 h
INSTRUM   CAB AV4 400 MHZ BASIC
PROBHD    Z863001_0028 (
PULPROG           zgpg30
TD                 65536
SOLVENT              MeOD
NS                  3042
DS                     4
SWH            23809.523 Hz
FIDRES          0.726609 Hz
AQ             1.3762560 sec
RG                   101
DW                21.000 usec
DE                  6.50 usec
TE                 295.8 K
D1            2.00000000 sec
D11           0.03000000 sec
TD0                    1
SFO1         100.6228298 MHz
NUC1                 13C
P0                  3.33 usec
P1                 10.00 usec
PLW1         53.28699875 W
SFO2         400.1316005 MHz
NUC2                  1H
CPDPRG[2         waltz65
PCPD2              90.00 usec
PLW2         10.22500038 W
PLW12         0.36482000 W
PLW13         0.18350001 W

F2 - Processing parameters
SI                 32768
SF           100.6127685 MHz
WDW                   EM
SSB      0
LB                  1.00 Hz
GB       0
PC                  1.40
```

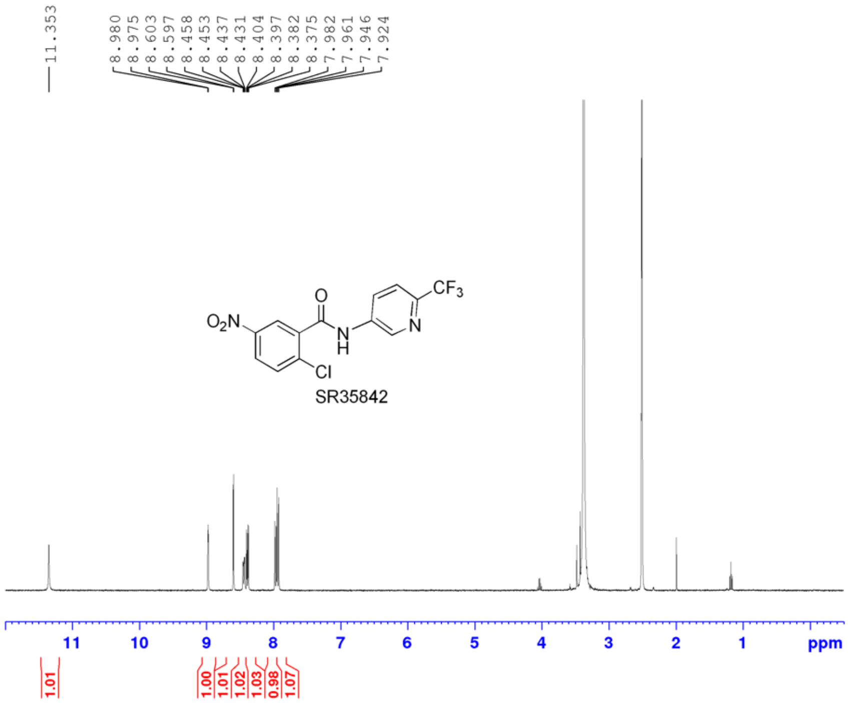


```
Current Data Parameters
NAME          zd-SR35842
EXPNO                 10
PROCNO                 1

F2 - Acquisition Parameters
Date_           20220815
Time             22.38 h
INSTRUM  CAB AV4 400 MHZ BASIC
PROBHD   Z863001_0028 (
PULPROG             zg30
TD                 65536
SOLVENT             DMSO
NS                    16
DS                     2
SWH             8196.722 Hz
FIDRES          0.250144 Hz
AQ             3.9976959 sec
RG                   101
DW                61.000 usec
DE                 12.35 usec
TE                 295.5 K
D1            2.00000000 sec
TD0                    1
SFO1         400.1324708 MHz
NUC1                  1H
P0                  5.67 usec
P1                 17.00 usec
PLW1         10.22500038 W

F2 - Processing parameters
SI                 65536
SF           400.1300000 MHz
WDW                   EM
SSB      0
LB                  0.30 Hz
GB       0
PC                  1.00
```

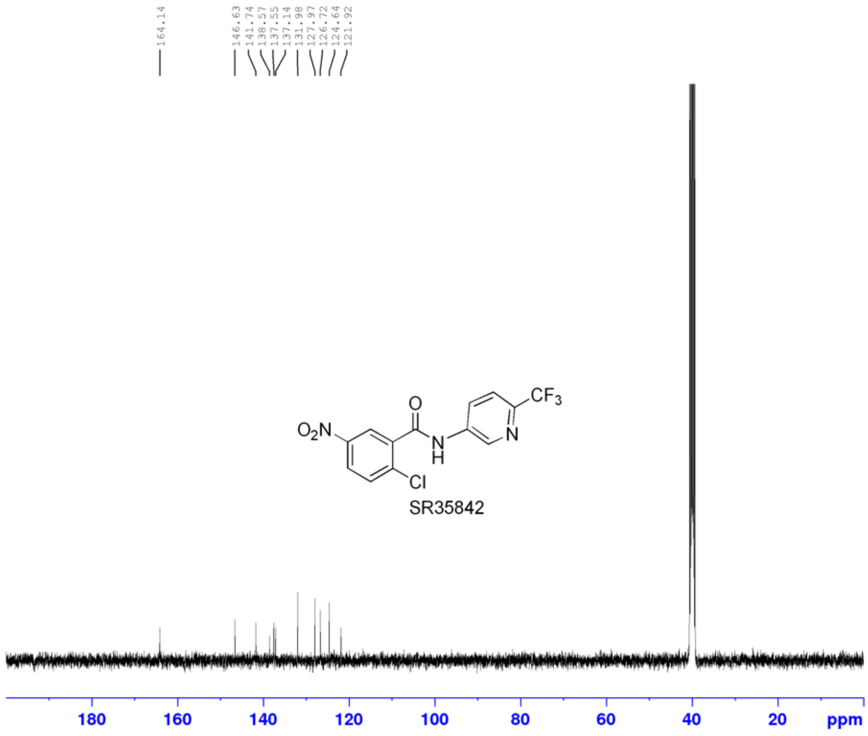


```
Current Data Parameters
NAME          zd-SR35842
EXPNO                 11
PROCNO                 1

F2 - Acquisition Parameters
Date_           20220816
Time              0.36 h
INSTRUM  CAB AV4 400 MHZ BASIC
PROBHD   Z863001_0028 (
PULPROG           zgpg30
TD                 65536
SOLVENT             DMSO
NS                  2048
DS                     4
SWH            23809.523 Hz
FIDRES          0.726609 Hz
AQ             1.3762560 sec
RG                   101
DW                21.000 usec
DE                  6.50 usec
TE                 296.6 K
D1            2.00000000 sec
D11           0.03000000 sec
TD0                    1
SFO1         100.6228298 MHz
NUC1                 13C
P0                  3.33 usec
P1                 10.00 usec
PLW1         53.28699875 W
SFO2         400.1316005 MHz
NUC2                  1H
CPDPRG[2         waltz65
PCPD2              90.00 usec
PLW2         10.22500038 W
PLW12         0.36482000 W
PLW13         0.18350001 W

F2 - Processing parameters
SI                 32768
SF           100.6127685 MHz
WDW                   EM
SSB      0
LB                  1.00 Hz
GB       0
PC                  1.40
```

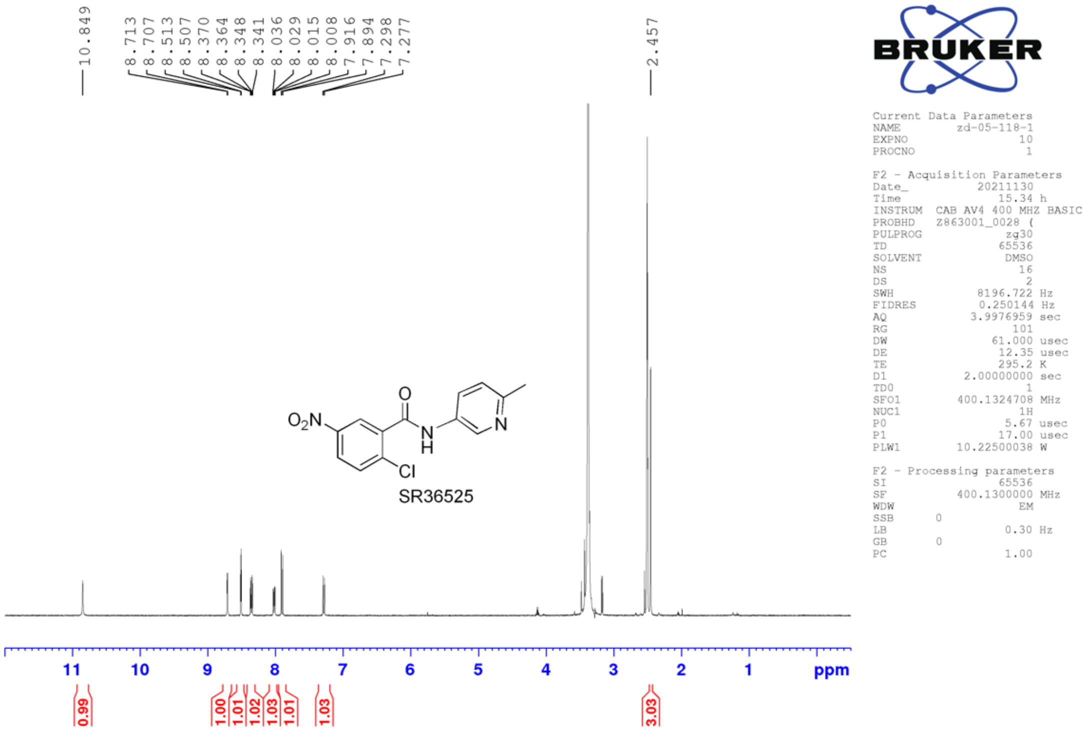
10.849
8.713
8.707
8.513
8.507
8.370
8.364
8.348
8.341
8.036
8.029
8.015
8.008
7.916
7.894
7.298
7.277
2.457
$O_2N$
O
N
H
N
Cl
SR36525
BRUKER
Current Data Parameters
NAME zd-05-118-1
EXPNO 10
PROCNO 1
F2 - Acquisition Parameters
Date_ 20211130
Time 15.34 h
INSTRUM CAB AV4 400 MHZ BASIC
PROBHD Z863001_0028 (
PULPROG zg30
TD 65536
SOLVENT DMSO
NS 16
DS 2
SWH 8196.722 Hz
FIDRES 0.250144 Hz
AQ 3.9976959 sec
RG 101
DW 61.000 usec
DE 12.35 usec
TE 295.2 K
D1 2.00000000 sec
TD0 1
SFO1 400.1324708 MHz
NUC1 1H
P0 5.67 usec
P1 17.00 usec
PLW1 10.22500038 W
F2 - Processing parameters
SI 65536
SF 400.1300000 MHz
WDW EM
SSB 0
LB 0.30 Hz
GB 0
PC 1.00
11 10 9 8 7 6 5 4 3 2 1 ppm
0.99
1.00
1.01
1.02
1.03
1.01
1.03
3.03

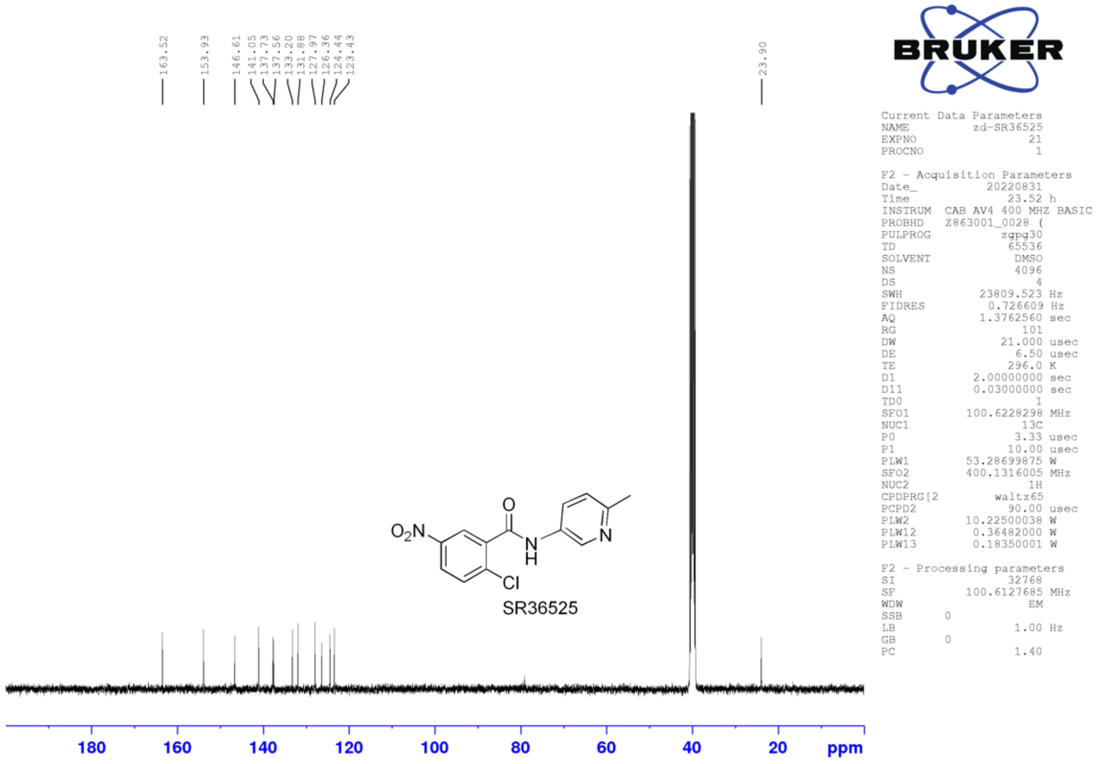
163.52
153.93
146.61
141.05
137.73
137.56
133.20
131.88
127.97
126.36
124.44
123.43
23.90
$O_2N$
O
N
H
N
Cl
SR36525
BRUKER
Current Data Parameters
NAME zd-SR36525
EXPNO 21
PROCNO 1
F2 - Acquisition Parameters
Date_ 20220831
Time 23.52 h
INSTRUM CAB AV4 400 MHZ BASIC
PROBHD Z863001_0028 (
PULPROG zgpg30
TD 65536
SOLVENT DMSO
NS 4096
DS 4
SWH 23809.523 Hz
FIDRES 0.726609 Hz
AQ 1.3762560 sec
RG 101
DW 21.000 usec
DE 6.50 usec
TE 296.0 K
D1 2.00000000 sec
D11 0.03000000 sec
TD0 1
SFO1 100.6228298 MHz
NUC1 13C
P0 3.33 usec
P1 10.00 usec
PLW1 53.28699875 W
SFO2 400.1316005 MHz
NUC2 1H
CPDPRG[2 waltz65
PCPD2 90.00 usec
PLW2 10.22500038 W
PLW12 0.36482000 W
PLW13 0.18350001 W
F2 - Processing parameters
SI 32768
SF 100.6127685 MHz
WDW EM
SSB 0
LB 1.00 Hz
GB 0
PC 1.40
180 160 140 120 100 80 60 40 20 ppm

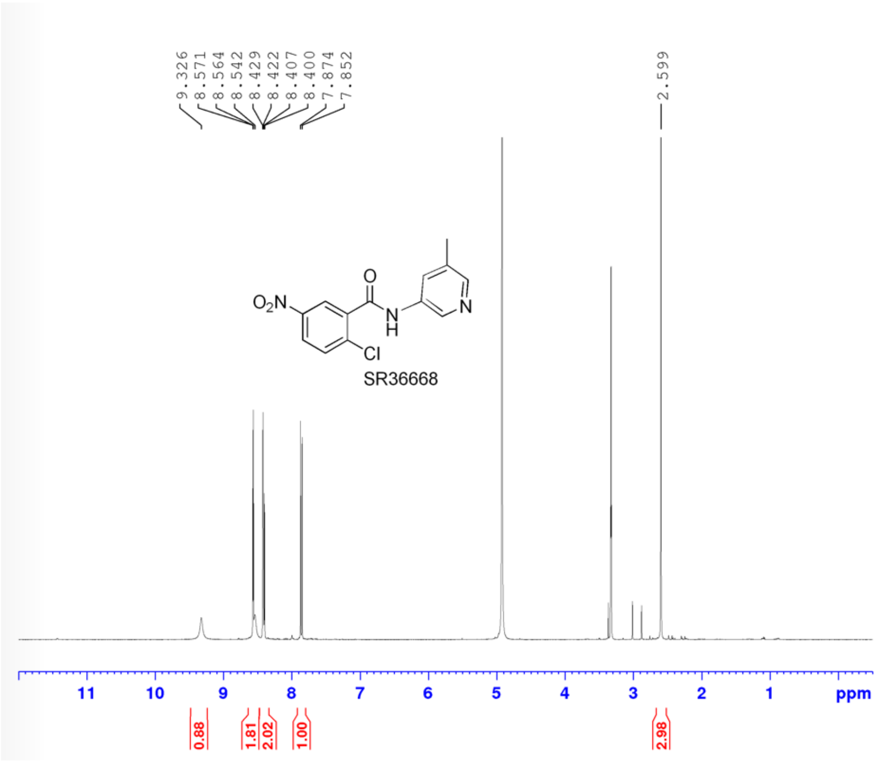


Current Data Parameters
NAME zd-05-118-3n-RT=1.414
EXPNO 10
PROCNO 1

F2 - Acquisition Parameters
Date_ 20220106
Time 18.56 h
INSTRUM CAB AV4 400 MHZ BASIC
PROBHD Z863001_0028 (
PULPROG zg30
TD 65536
SOLVENT MeOD
NS 16
DS 2
SWH 8196.722 Hz
FIDRES 0.250144 Hz
AQ 3.9976959 sec
RG 101
DW 61.000 usec
DE 12.35 usec
TE 295.3 K
D1 2.00000000 sec
TD0 1
SFO1 400.1324708 MHz
NUC1 1H
P0 5.67 usec
P1 17.00 usec
PLW1 10.22500038 W

F2 - Processing parameters
SI 65536
SF 400.1300000 MHz
WDW EM
SSB 0
LB 0.30 Hz
GB 0
PC 1.00

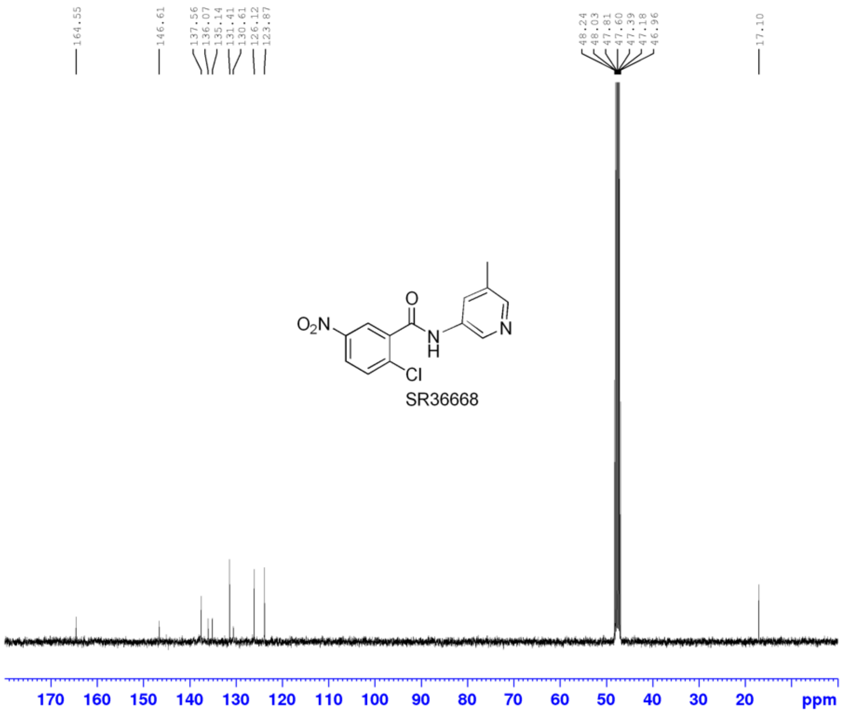


Current Data Parameters
NAME zd-05-118-3n-RT=1.414
EXPNO 11
PROCNO 1

F2 - Acquisition Parameters
Date_ 20220106
Time 19.28 h
INSTRUM CAB AV4 400 MHZ BASIC
PROBHD Z863001_0028 (
PULPROG zgpg30
TD 65536
SOLVENT MeOD
NS 450
DS 4
SWH 23809.523 Hz
FIDRES 0.726609 Hz
AQ 1.3762560 sec
RG 101
DW 21.000 usec
DE 6.50 usec
TE 296.5 K
D1 2.00000000 sec
D11 0.03000000 sec
TD0 1
SFO1 100.6228298 MHz
NUC1 13C
P0 3.33 usec
P1 10.00 usec
PLW1 53.28699875 W
SFO2 400.1316005 MHz
NUC2 1H
CPDPRG[2 waltz65
PCPD2 90.00 usec
PLW2 10.22500038 W
PLW12 0.36482000 W
PLW13 0.18350001 W

F2 - Processing parameters
SI 32768
SF 100.6127685 MHz
WDW EM
SSB 0
LB 1.00 Hz
GB 0
PC 1.40

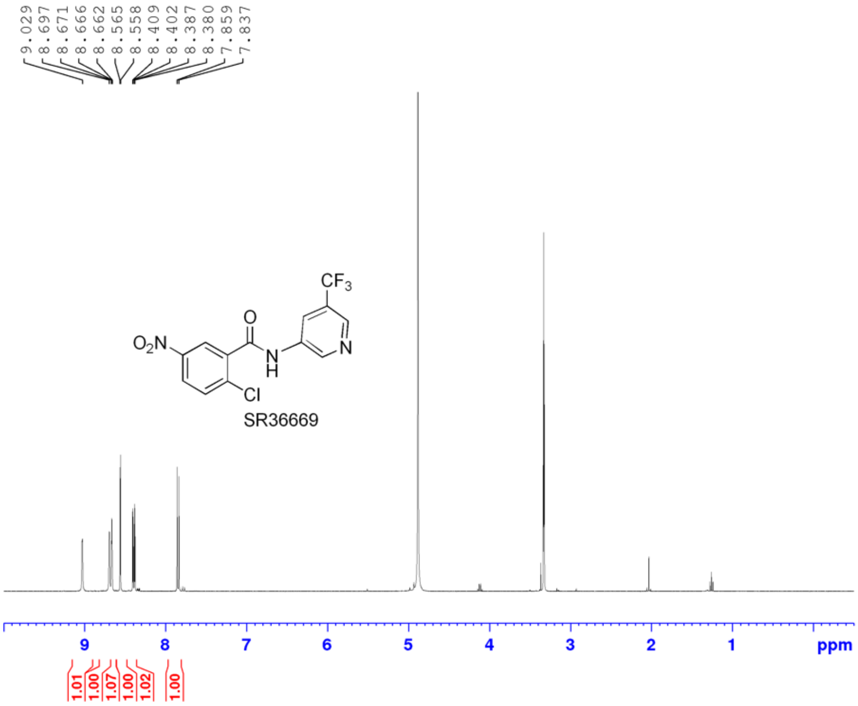


```
Current Data Parameters
NAME          zd-05-118-4
EXPNO                  10
PROCNO                  1

F2 - Acquisition Parameters
Date_            20220106
Time                15.07 h
INSTRUM   CAB AV4 400 MHZ BASIC
PROBHD    Z863001_0028 (
PULPROG              zg30
TD                  65536
SOLVENT              MeOD
NS                     16
DS                      2
SWH              8196.722 Hz
FIDRES           0.250144 Hz
AQ              3.9976959 sec
RG                    101
DW                 61.000 usec
DE                  12.35 usec
TE                  295.3 K
D1             2.00000000 sec
TD0                     1
SFO1          400.1324708 MHz
NUC1                   1H
P0                   5.67 usec
P1                  17.00 usec
PLW1          10.22500038 W

F2 - Processing parameters
SI                  65536
SF            400.1300000 MHz
WDW                    EM
SSB      0
LB                   0.30 Hz
GB       0
PC                   1.00
```

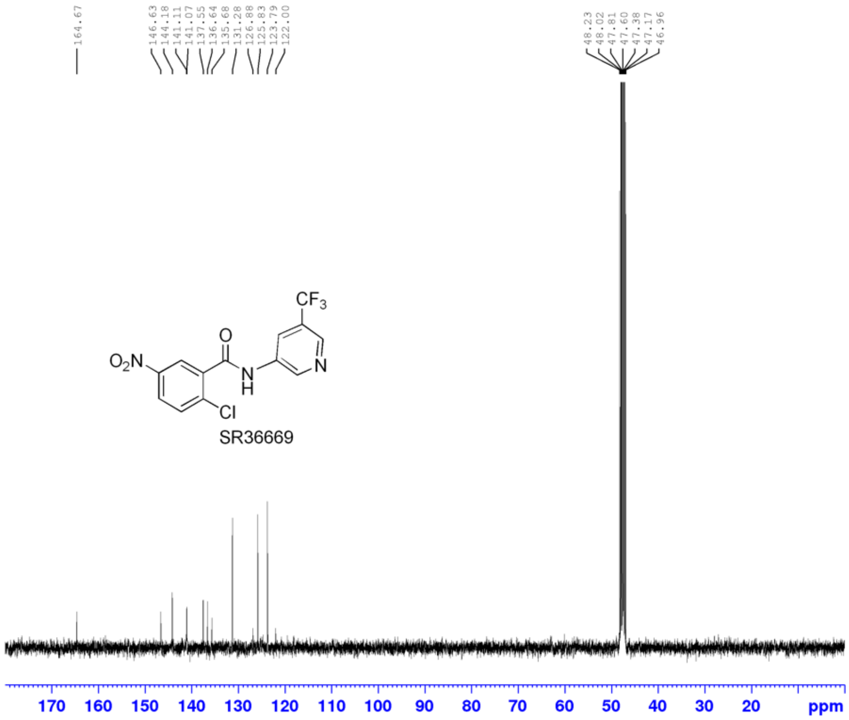


```
Current Data Parameters
NAME          zd-05-118-4
EXPNO                  20
PROCNO                  1

F2 - Acquisition Parameters
Date_            20220106
Time                14.06 h
INSTRUM   CAB AV4 400 MHZ BASIC
PROBHD    Z863001_0028 (
PULPROG            zgpg30
TD                  65536
SOLVENT              MeOD
NS                    400
DS                      4
SWH             23809.523 Hz
FIDRES           0.726609 Hz
AQ              1.3762560 sec
RG                    101
DW                 21.000 usec
DE                   6.50 usec
TE                  296.5 K
D1             2.00000000 sec
D11            0.03000000 sec
TD0                     1
SFO1          100.6228298 MHz
NUC1                  13C
P0                   3.33 usec
P1                  10.00 usec
PLW1          53.28699875 W
SFO2          400.1316005 MHz
NUC2                   1H
CPDPRG[2          waltz65
PCPD2               90.00 usec
PLW2          10.22500038 W
PLW12          0.36482000 W
PLW13          0.18350001 W

F2 - Processing parameters
SI                  32768
SF            100.6127685 MHz
WDW                    EM
SSB      0
LB                   1.00 Hz
GB       0
PC                   1.40
```

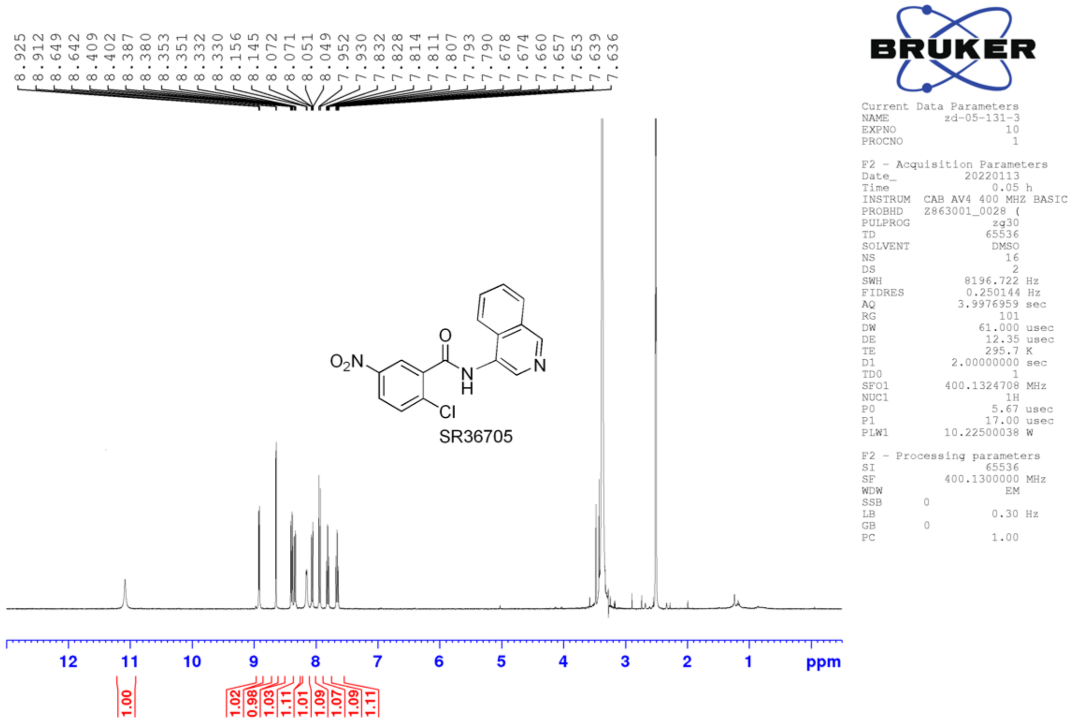
8.925
8.912
8.649
8.642
8.409
8.402
8.387
8.380
8.353
8.351
8.332
8.330
8.156
8.145
8.072
8.071
8.051
8.049
7.952
7.930
7.832
7.828
7.814
7.811
7.807
7.793
7.790
7.678
7.674
7.660
7.657
7.653
7.639
7.636
SR36705
1.00
1.02
0.98
1.03
1.11
1.01
1.09
1.07
1.09
1.11
12
11
10
9
8
7
6
5
4
3
2
1
ppm
BRUKER
Current Data Parameters
NAME zd-05-131-3
EXPNO 10
PROCNO 1
F2 - Acquisition Parameters
Date_ 20220113
Time 0.05 h
INSTRUM CAB AV4 400 MHZ BASIC
PROBHD Z863001_0028 (
PULPROG zg30
TD 65536
SOLVENT DMSO
NS 16
DS 2
SWH 8196.722 Hz
FIDRES 0.250144 Hz
AQ 3.9976959 sec
RG 101
DW 61.000 usec
DE 12.35 usec
TE 295.7 K
D1 2.00000000 sec
TD0 1
SFO1 400.1324708 MHz
NUC1 1H
P0 5.67 usec
P1 17.00 usec
PLW1 10.22500038 W
F2 - Processing parameters
SI 65536
SF 400.1300000 MHz
WDW EM
SSB 0
LB 0.30 Hz
GB 0
PC 1.00

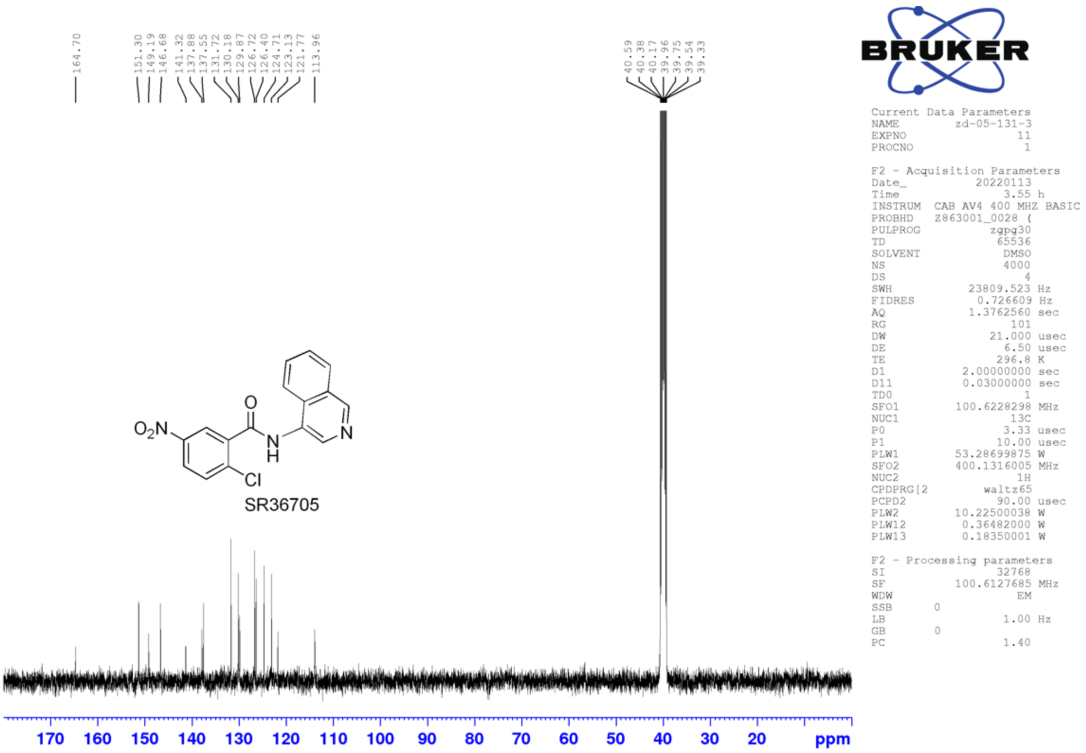
164.70
151.30
149.19
146.68
141.32
137.88
137.55
131.72
130.18
129.87
126.72
126.40
124.71
123.13
121.77
113.96
40.59
40.38
40.17
39.96
39.75
39.54
39.33
SR36705
170
160
150
140
130
120
110
100
90
80
70
60
50
40
30
20
ppm
BRUKER
Current Data Parameters
NAME zd-05-131-3
EXPNO 11
PROCNO 1
F2 - Acquisition Parameters
Date_ 20220113
Time 3.55 h
INSTRUM CAB AV4 400 MHZ BASIC
PROBHD Z863001_0028 (
PULPROG zgpg30
TD 65536
SOLVENT DMSO
NS 4000
DS 4
SWH 23809.523 Hz
FIDRES 0.726609 Hz
AQ 1.3762560 sec
RG 101
DW 21.000 usec
DE 6.50 usec
TE 296.8 K
D1 2.00000000 sec
D11 0.03000000 sec
TD0 1
SFO1 100.6228298 MHz
NUC1 13C
P0 3.33 usec
P1 10.00 usec
PLW1 53.28699875 W
SFO2 400.1316005 MHz
NUC2 1H
CPDPRG[2 waltz65
PCPD2 90.00 usec
PLW2 10.22500038 W
PLW12 0.36482000 W
PLW13 0.18350001 W
F2 - Processing parameters
SI 32768
SF 100.6127685 MHz
WDW EM
SSB 0
LB 1.00 Hz
GB 0
PC 1.40

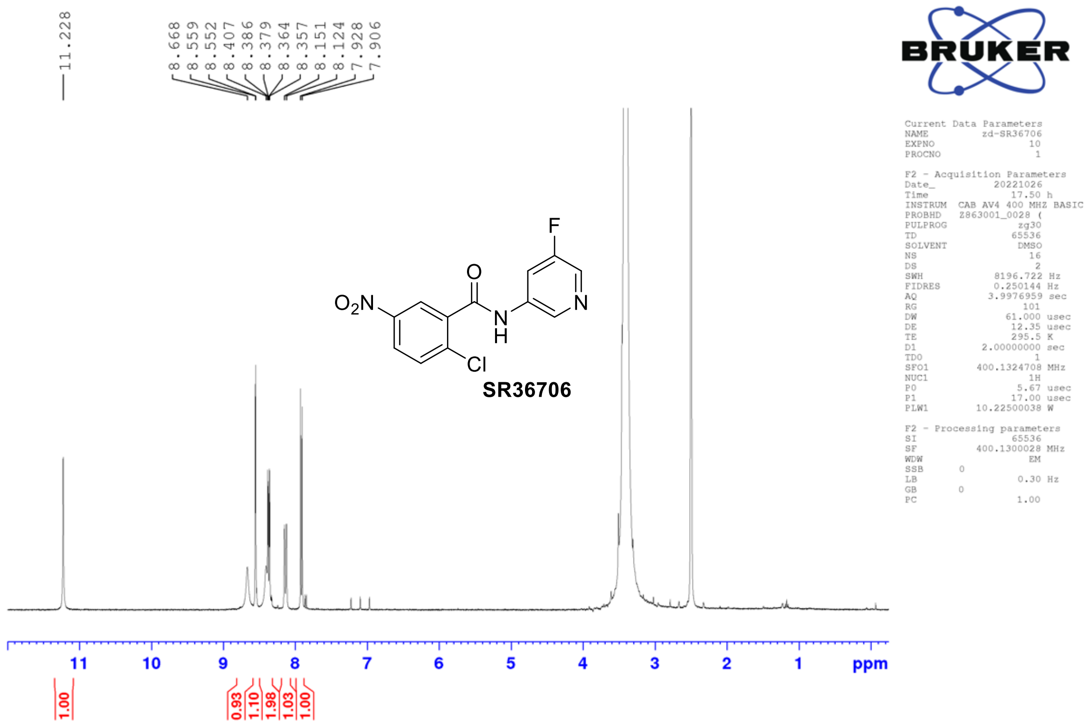
11.228
8.668
8.559
8.552
8.407
8.386
8.379
8.364
8.357
8.151
8.124
7.928
7.906
O$_2$N
O
N
H
F
N
Cl
SR36706
11
10
9
8
7
6
5
4
3
2
1
ppm
1.00
0.93
1.10
1.98
1.03
1.00
BRUKER
Current Data Parameters
NAME zd-SR36706
EXPNO 10
PROCNO 1
F2 - Acquisition Parameters
Date_ 20221026
Time 17.50 h
INSTRUM CAB AV4 400 MHZ BASIC
PROBHD Z863001_0028 (
PULPROG zg30
TD 65536
SOLVENT DMSO
NS 16
DS 2
SWH 8196.722 Hz
FIDRES 0.250144 Hz
AQ 3.9976959 sec
RG 101
DW 61.000 usec
DE 12.35 usec
TE 295.5 K
D1 2.00000000 sec
TD0 1
SFO1 400.1324708 MHz
NUC1 1H
P0 5.67 usec
P1 17.00 usec
PLW1 10.22500038 W
F2 - Processing parameters
SI 65536
SF 400.1300028 MHz
WDW EM
SSB 0
LB 0.30 Hz
GB 0
PC 1.00

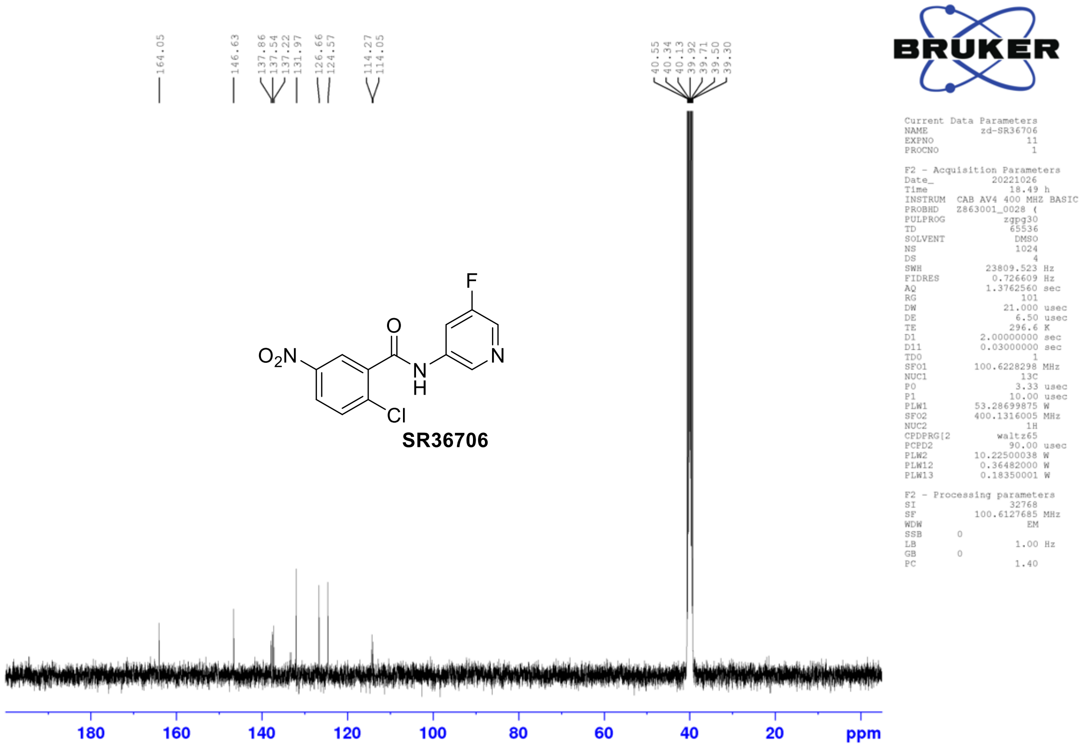
164.05
146.63
137.86
137.54
137.22
131.97
126.66
124.57
114.27
114.05
40.55
40.34
40.13
39.92
39.71
39.50
39.30
O$_2$N
O
N
H
F
N
Cl
SR36706
180
160
140
120
100
80
60
40
20
ppm
BRUKER
Current Data Parameters
NAME zd-SR36706
EXPNO 11
PROCNO 1
F2 - Acquisition Parameters
Date_ 20221026
Time 18.49 h
INSTRUM CAB AV4 400 MHZ BASIC
PROBHD Z863001_0028 (
PULPROG zgpg30
TD 65536
SOLVENT DMSO
NS 1024
DS 4
SWH 23809.523 Hz
FIDRES 0.726609 Hz
AQ 1.3762560 sec
RG 101
DW 21.000 usec
DE 6.50 usec
TE 296.6 K
D1 2.00000000 sec
D11 0.03000000 sec
TD0 1
SFO1 100.6228298 MHz
NUC1 13C
P0 3.33 usec
P1 10.00 usec
PLW1 53.28699875 W
SFO2 400.1316005 MHz
NUC2 1H
CPDPRG[2 waltz65
PCPD2 90.00 usec
PLW2 10.22500038 W
PLW12 0.36482000 W
PLW13 0.18350001 W
F2 - Processing parameters
SI 32768
SF 100.6127685 MHz
WDW EM
SSB 0
LB 1.00 Hz
GB 0
PC 1.40

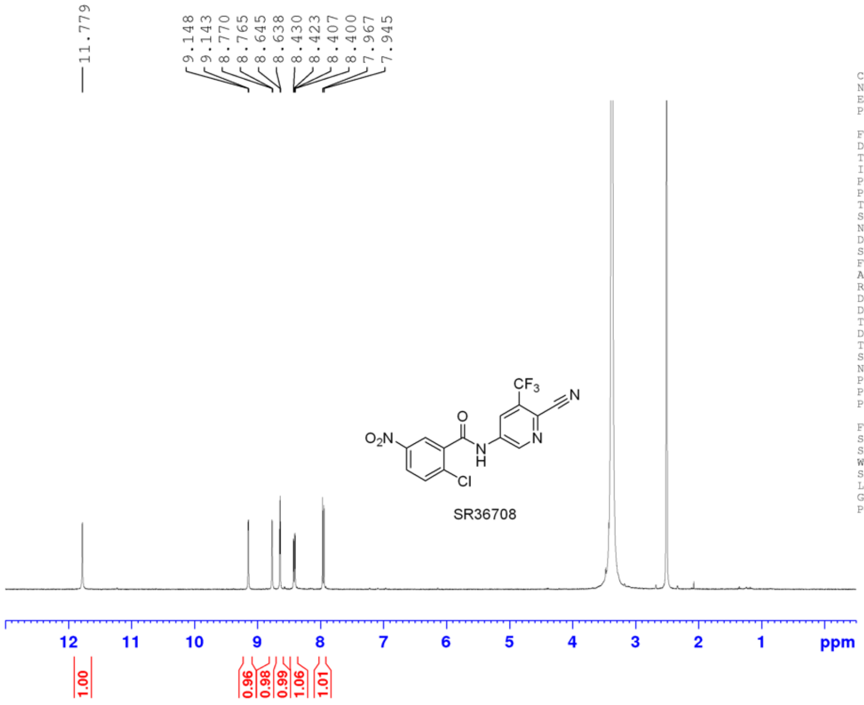


Current Data Parameters
NAME zd-05-132-3
EXPNO 10
PROCNO 1

F2 - Acquisition Parameters
Date_ 20220118
Time 10.56 h
INSTRUM CAB AV4 400 MHZ BASIC
PROBHD Z863001_0028 (
PULPROG zg30
TD 65536
SOLVENT DMSO
NS 16
DS 2
SWH 8196.722 Hz
FIDRES 0.250144 Hz
AQ 3.9976959 sec
RG 101
DW 61.000 usec
DE 12.35 usec
TE 295.2 K
D1 2.00000000 sec
TD0 1
SFO1 400.1324708 MHz
NUC1 1H
P0 5.67 usec
P1 17.00 usec
PLW1 10.22500038 W

F2 - Processing parameters
SI 65536
SF 400.1300000 MHz
WDW EM
SSB 0
LB 0.30 Hz
GB 0
PC 1.00

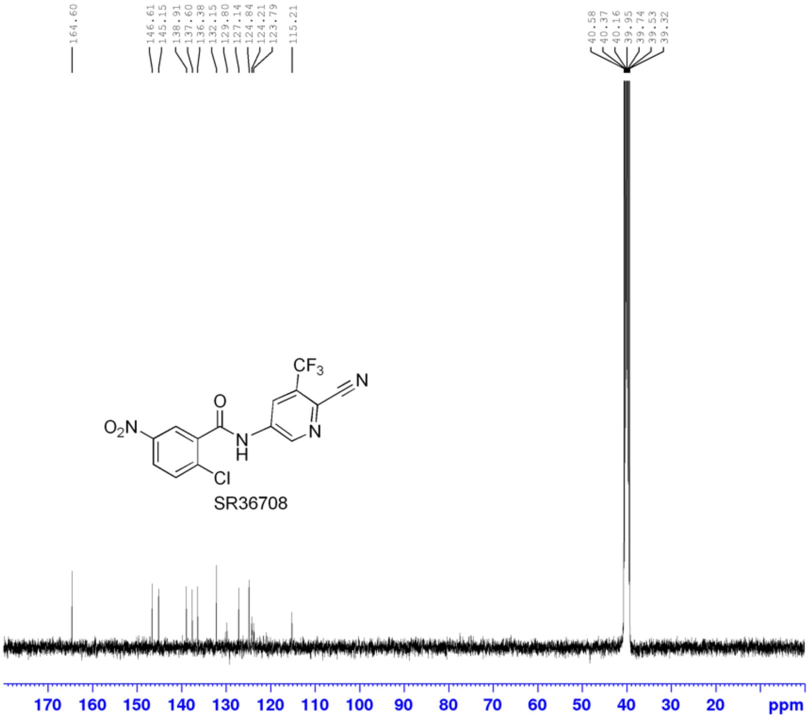


Current Data Parameters
NAME zd-05-132-3
EXPNO 20
PROCNO 1

F2 - Acquisition Parameters
Date_ 20220119
Time 0.43 h
INSTRUM CAB AV4 400 MHZ BASIC
PROBHD Z863001_0028 (
PULPROG zgpg30
TD 65536
SOLVENT DMSO
NS 4500
DS 4
SWH 23809.523 Hz
FIDRES 0.726609 Hz
AQ 1.3762560 sec
RG 101
DW 21.000 usec
DE 6.50 usec
TE 295.8 K
D1 2.00000000 sec
D11 0.03000000 sec
TD0 1
SFO1 100.6228298 MHz
NUC1 13C
P0 3.33 usec
P1 10.00 usec
PLW1 53.28699875 W
SFO2 400.1316005 MHz
NUC2 1H
CPDPRG[2 waltz65
PCPD2 90.00 usec
PLW2 10.22500038 W
PLW12 0.36482000 W
PLW13 0.18350001 W

F2 - Processing parameters
SI 32768
SF 100.6127685 MHz
WDW EM
SSB 0
LB 1.00 Hz
GB 0
PC 1.40

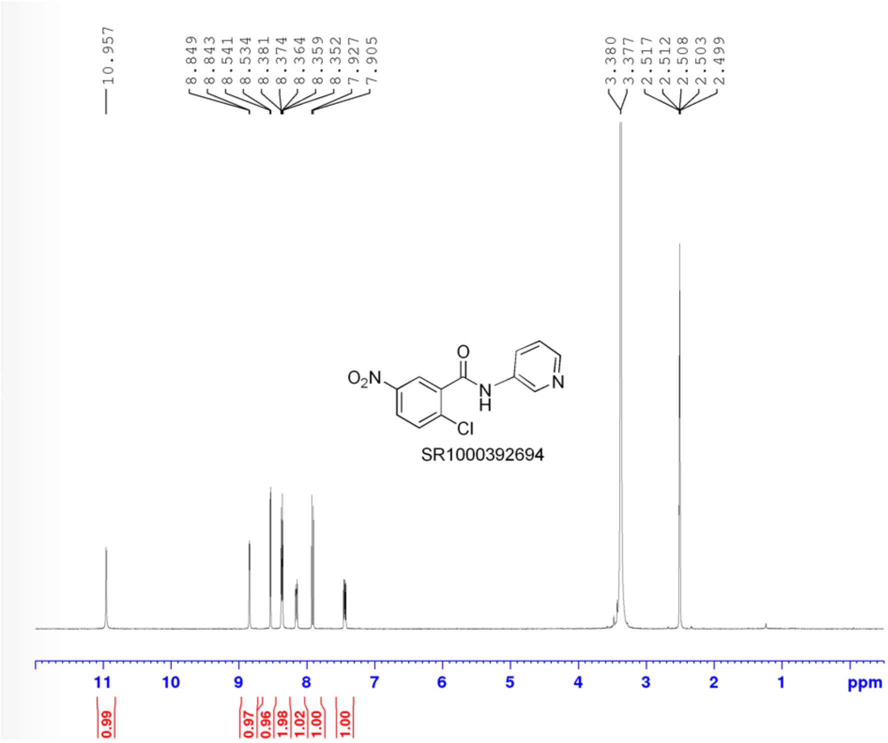


```
Current Data Parameters
NAME      zd-03-98-2-retest
EXPNO                    10
PROCNO                    1

F2 - Acquisition Parameters
Date_              20220814
Time                  16.03 h
INSTRUM   CAB AV4 400 MHZ BASIC
PROBHD    Z863001_0028 (
PULPROG                zg30
TD                    65536
SOLVENT                DMSO
NS                       16
DS                        2
SWH                8196.722 Hz
FIDRES             0.250144 Hz
AQ                3.9976959 sec
RG                      101
DW                   61.000 usec
DE                    12.35 usec
TE                    295.7 K
D1               2.00000000 sec
TD0                       1
SFO1            400.1324708 MHz
NUC1                     1H
P0                     5.67 usec
P1                    17.00 usec
PLW1            10.22500038 W

F2 - Processing parameters
SI                    65536
SF              400.1300000 MHz
WDW                      EM
SSB       0
LB                     0.30 Hz
GB        0
PC                     1.00
```

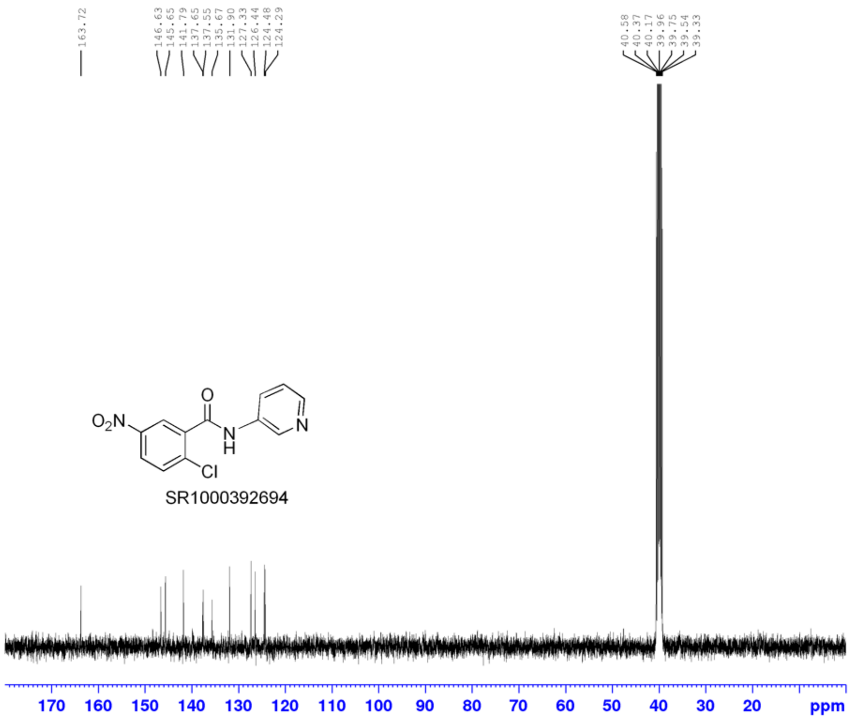


```
Current Data Parameters
NAME      zd-03-98-2-retest
EXPNO                    11
PROCNO                    1

F2 - Acquisition Parameters
Date_              20220814
Time                  16.33 h
INSTRUM   CAB AV4 400 MHZ BASIC
PROBHD    Z863001_0028 (
PULPROG              zgpg30
TD                    65536
SOLVENT                DMSO
NS                      500
DS                        4
SWH               23809.523 Hz
FIDRES             0.726609 Hz
AQ                1.3762560 sec
RG                      101
DW                   21.000 usec
DE                     6.50 usec
TE                    296.8 K
D1               2.00000000 sec
D11              0.03000000 sec
TD0                       1
SFO1            100.6228298 MHz
NUC1                    13C
P0                     3.33 usec
P1                    10.00 usec
PLW1            53.28699875 W
SFO2            400.1316005 MHz
NUC2                     1H
CPDPRG[2                waltz65
PCPD2                 90.00 usec
PLW2            10.22500038 W
PLW12            0.36482000 W
PLW13            0.18350001 W

F2 - Processing parameters
SI                    32768
SF              100.6127685 MHz
WDW                      EM
SSB       0
LB                     1.00 Hz
GB        0
PC                     1.40
```

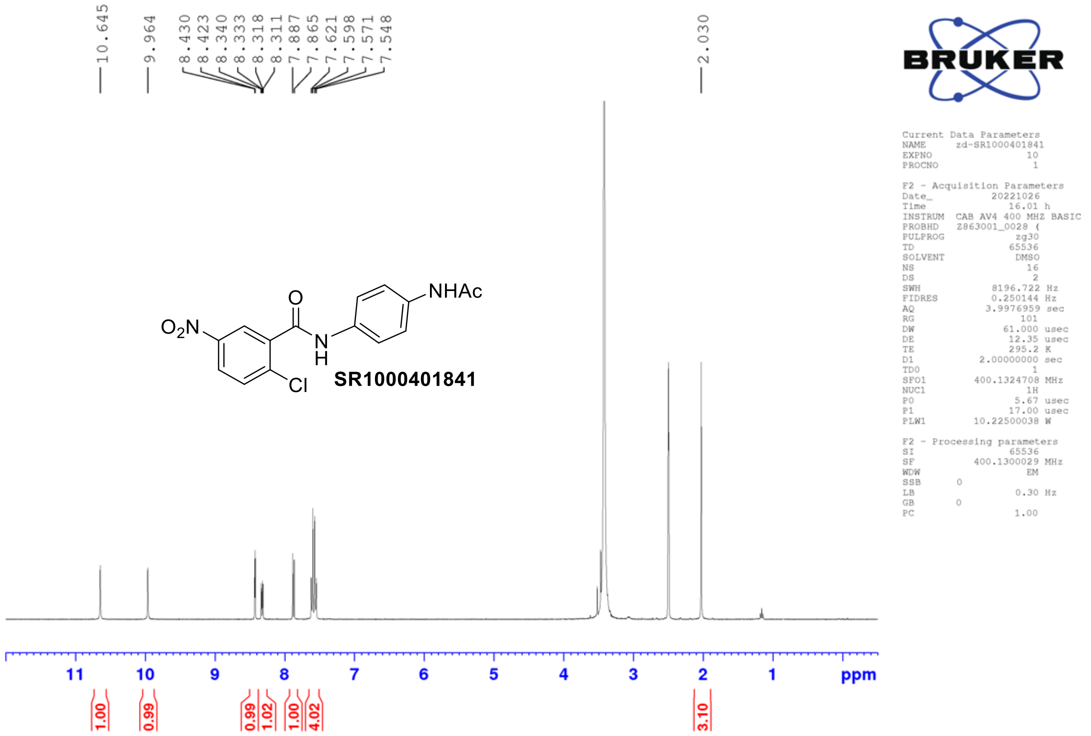
10.645
9.964
8.430
8.423
8.340
8.333
8.318
8.311
7.887
7.865
7.621
7.598
7.571
7.548
2.030
O
$O_2N$
N
H
Cl
NHAc
SR1000401841
BRUKER
Current Data Parameters
NAME zd-SR1000401841
EXPNO 10
PROCNO 1
F2 - Acquisition Parameters
Date_ 20221026
Time 16.01 h
INSTRUM CAB AV4 400 MHZ BASIC
PROBHD Z863001_0028 (
PULPROG zg30
TD 65536
SOLVENT DMSO
NS 16
DS 2
SWH 8196.722 Hz
FIDRES 0.250144 Hz
AQ 3.9976959 sec
RG 101
DW 61.000 usec
DE 12.35 usec
TE 295.2 K
D1 2.00000000 sec
TD0 1
SFO1 400.1324708 MHz
NUC1 1H
P0 5.67 usec
P1 17.00 usec
PLW1 10.22500038 W
F2 - Processing parameters
SI 65536
SF 400.1300029 MHz
WDW EM
SSB 0
LB 0.30 Hz
GB 0
PC 1.00
11
10
9
8
7
6
5
4
3
2
1
ppm
1.00
0.99
0.99
1.02
1.00
4.02
3.10

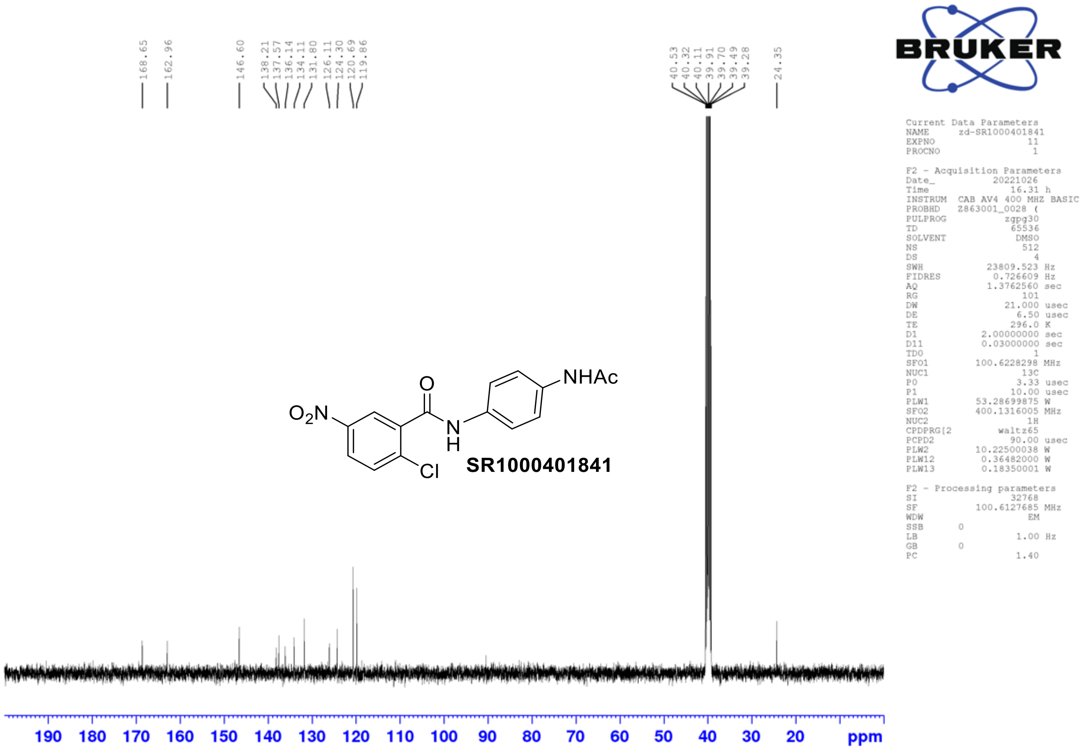
168.65
162.96
146.60
138.21
137.57
136.14
134.11
131.80
126.11
124.30
120.69
119.86
40.53
40.32
40.11
39.91
39.70
39.49
39.28
24.35
O
$O_2N$
N
H
Cl
NHAc
SR1000401841
BRUKER
Current Data Parameters
NAME zd-SR1000401841
EXPNO 11
PROCNO 1
F2 - Acquisition Parameters
Date_ 20221026
Time 16.31 h
INSTRUM CAB AV4 400 MHZ BASIC
PROBHD Z863001_0028 (
PULPROG zgpg30
TD 65536
SOLVENT DMSO
NS 512
DS 4
SWH 23809.523 Hz
FIDRES 0.726609 Hz
AQ 1.3762560 sec
RG 101
DW 21.000 usec
DE 6.50 usec
TE 296.0 K
D1 2.00000000 sec
D11 0.03000000 sec
TD0 1
SFO1 100.6228298 MHz
NUC1 13C
P0 3.33 usec
P1 10.00 usec
PLW1 53.28699875 W
SFO2 400.1316005 MHz
NUC2 1H
CPDPRG[2 waltz65
PCPD2 90.00 usec
PLW2 10.22500038 W
PLW12 0.36482000 W
PLW13 0.18350001 W
F2 - Processing parameters
SI 32768
SF 100.6127685 MHz
WDW EM
SSB 0
LB 1.00 Hz
GB 0
PC 1.40
190
180
170
160
150
140
130
120
110
100
90
80
70
60
50
40
30
20
ppm
