## Supplementary Figures S1-S5 and Tables S1-S2 for "Ligand efficacy shifts a nuclear receptor conformational ensemble between transcriptionally active and repressive states"

[1] Department of Integrative Structural and Computational Biology, Scripps Research and The Herbert Wertheim UF Scripps Institute for Biomedical Innovation & Technology, Jupiter, Florida 33458, United States
[2] Department of Molecular Medicine, Scripps Research and The Herbert Wertheim UF Scripps Institute for Biomedical Innovation & Technology, Jupiter, Florida 33458, United States
[3] School of Basic Medical Sciences, Guangzhou Laboratory, Guangzhou Medical University, Guangzhou, China.
[4] Department of Chemistry, Vanderbilt University, Nashville, Tennessee 37232, United States
[5] Center for Structural Biology, Vanderbilt University, Nashville, Tennessee 37232, United States
[6] Vanderbilt Institute of Chemical Biology, Vanderbilt University, Nashville, Tennessee 37232, United States
[7] Center for Applied AI in Protein Dynamics, Vanderbilt University, Nashville, Tennessee 37232, United States
[8] Data Science Institute, Vanderbilt University, Nashville, Tennessee 37232, United States
[9] Department of Chemical and Biomolecular Engineering, Vanderbilt University, Nashville, Tennessee 37232, United States
[10] Department of Biochemistry, Vanderbilt University, Nashville, Tennessee 37232, United States.
[#] Authors contributed equally

File contains:

- Supplementary Figures S1–S5
- Supplementary Tables S1–S2

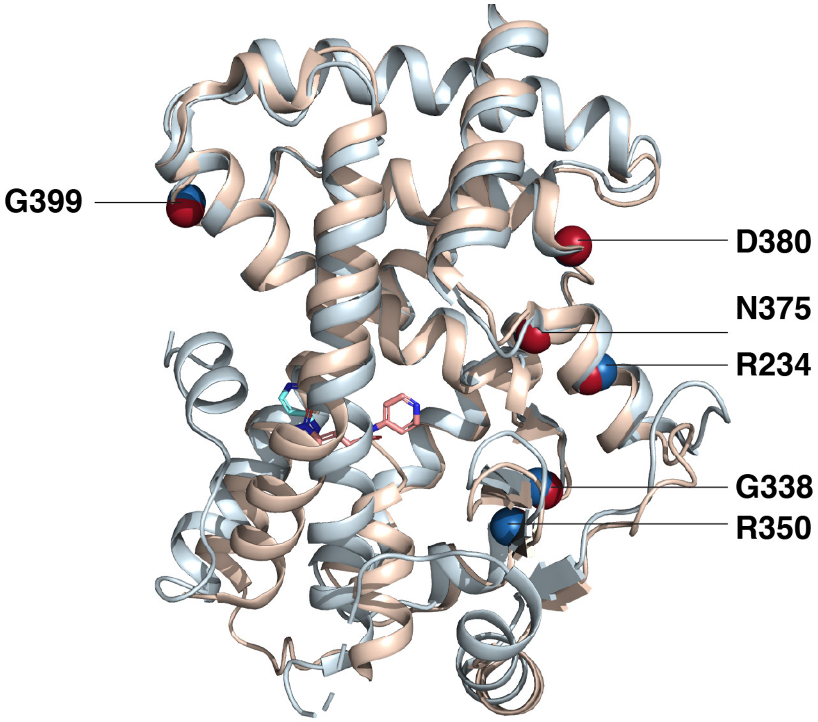


**Supplementary Figure S1.** Select PPARγ LBD residues with well-dispersed NMR peaks that display ligand-dependent shifts between active-like and repressive-like states.

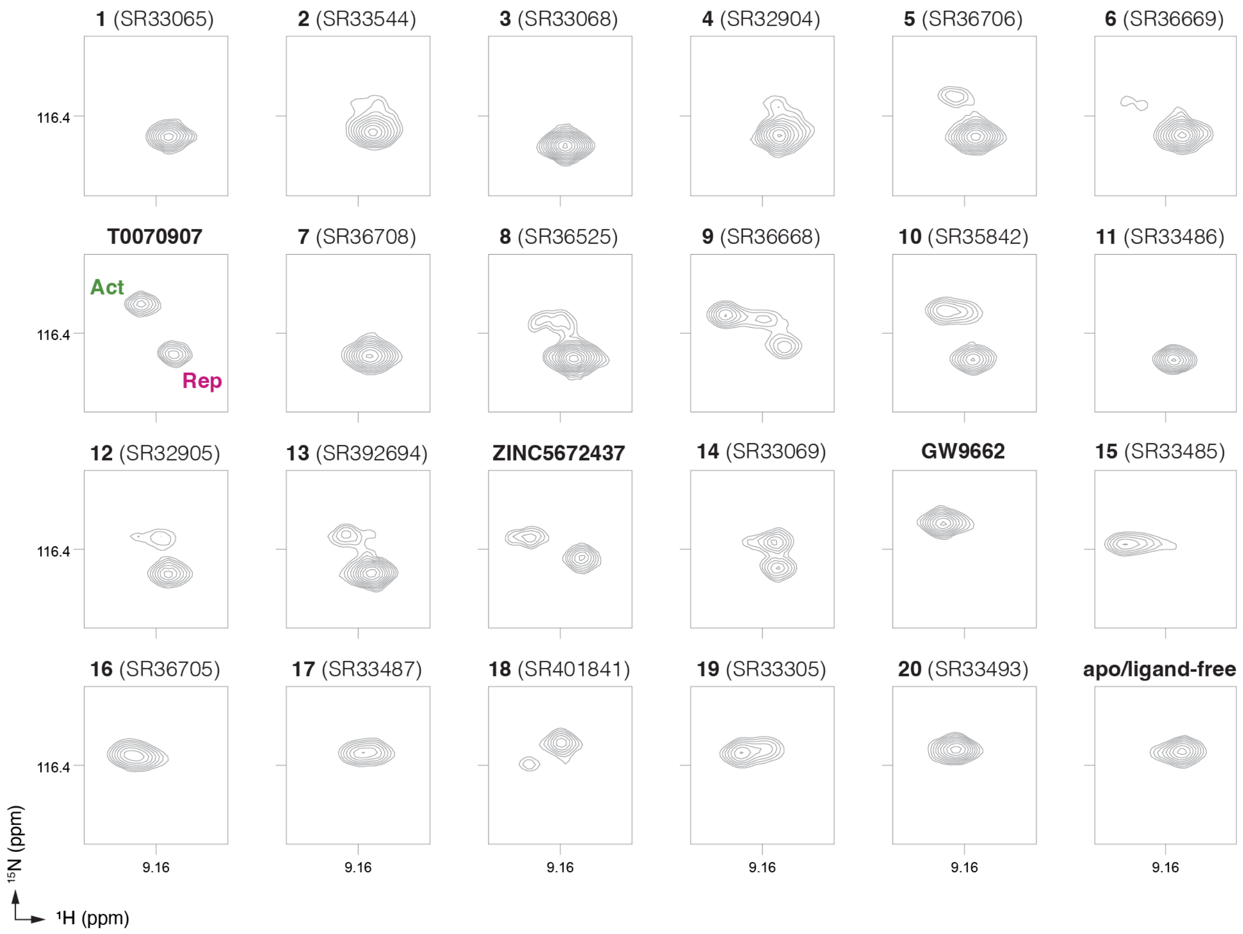


**Supplementary Figure S2.** 2D [$^{1}$H,$^{15}$N]-TROSY-HSQC NMR focused on Arg234 of $^{15}$N-labeled PPARγ LBD bound to compounds in the ligand series. The peak positions of the active-like (Act) and repressive-like (Rep) states populated by T0070907 are noted.

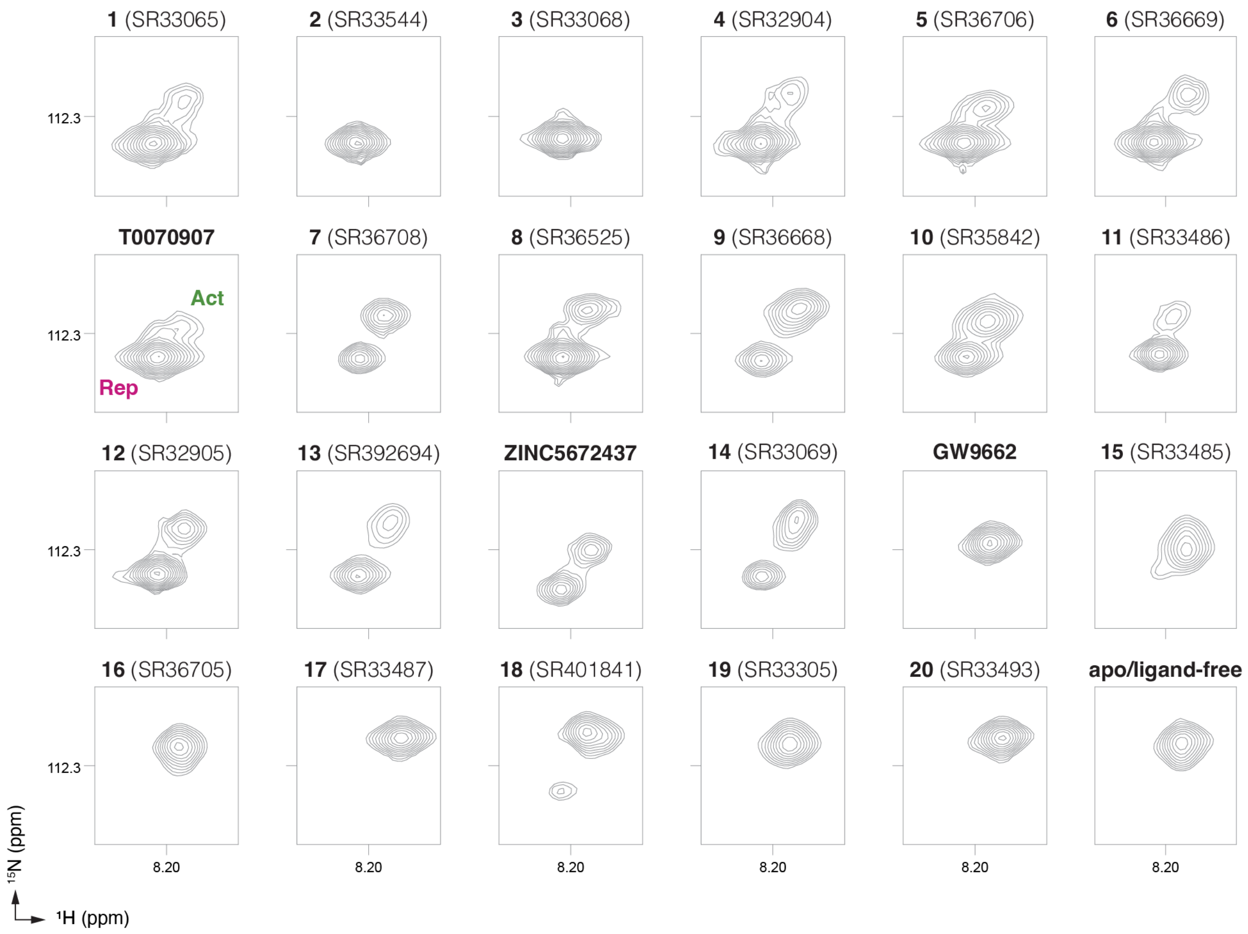


**Supplementary Figure S3.** 2D [$^1$H,$^{15}$N]-TROSY-HSQC NMR focused on Gly338 of $^{15}$N-labeled PPARγ LBD bound to compounds in the ligand series. The peak positions of the active-like (Act) and repressive-like (Rep) states populated by T0070907 are noted.

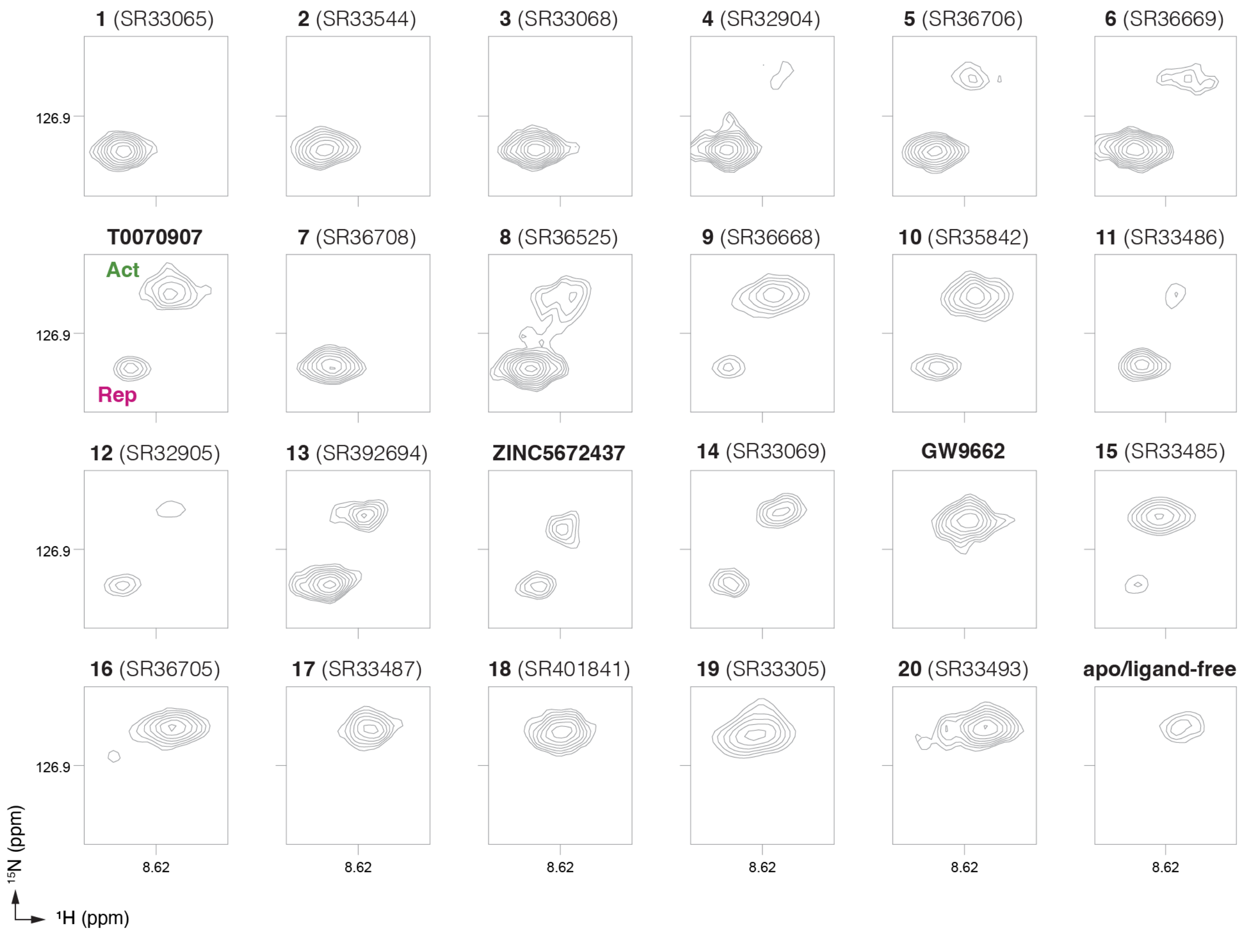


**Supplementary Figure S4.** 2D [$^{1}$H,$^{15}$N]-TROSY-HSQC NMR focused on Arg350 of $^{15}$N-labeled PPARγ LBD bound to compounds in the ligand series. The peak positions of the active-like (Act) and repressive-like (Rep) states populated by T0070907 are noted.

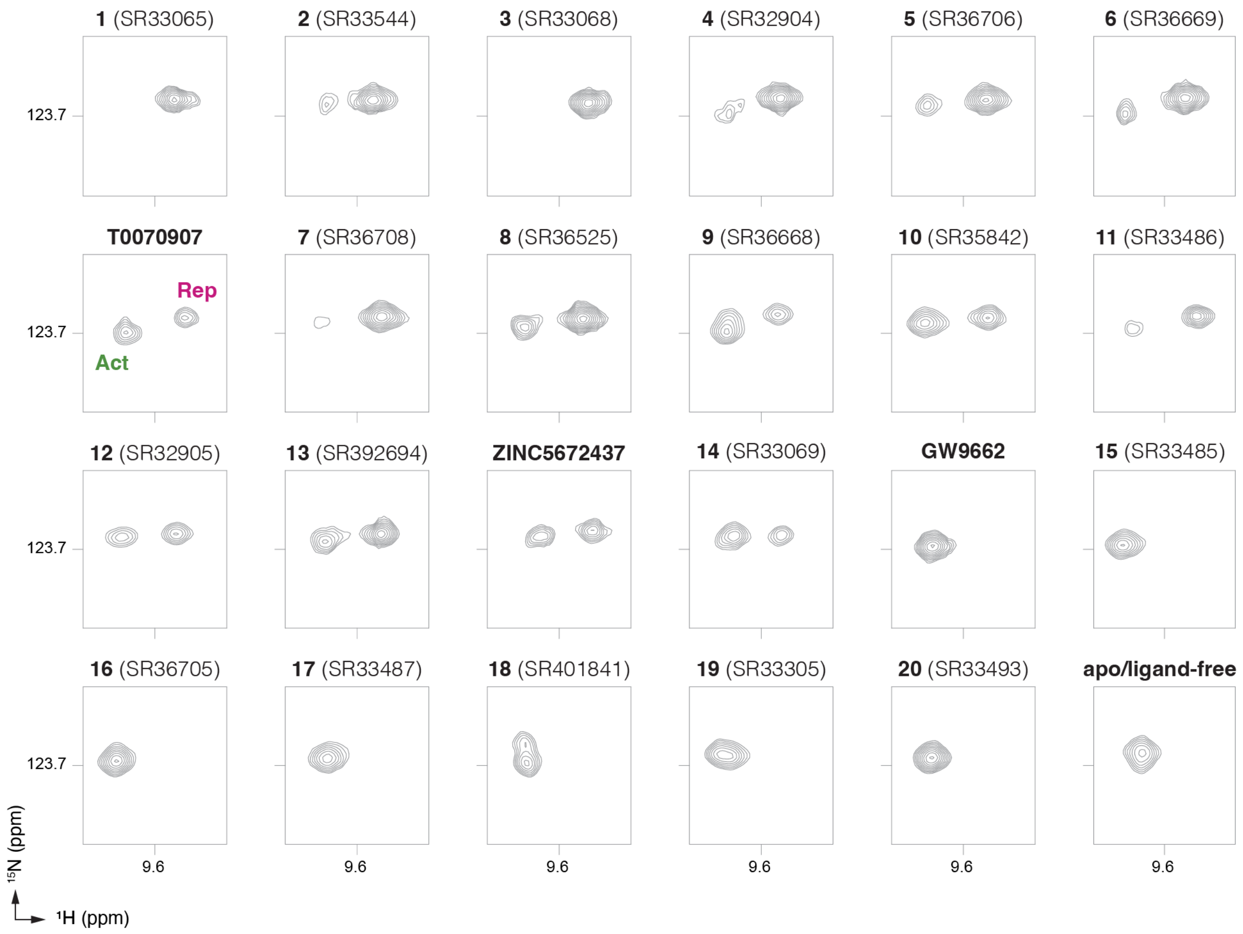


**Supplementary Figure S5.** 2D [$^1$H,$^{15}$N]-TROSY-HSQC NMR focused on Asn375 of $^{15}$N-labeled PPARγ LBD bound to compounds in the ligand series. The peak positions of the active-like (Act) and repressive-like (Rep) states populated by T0070907 are noted.

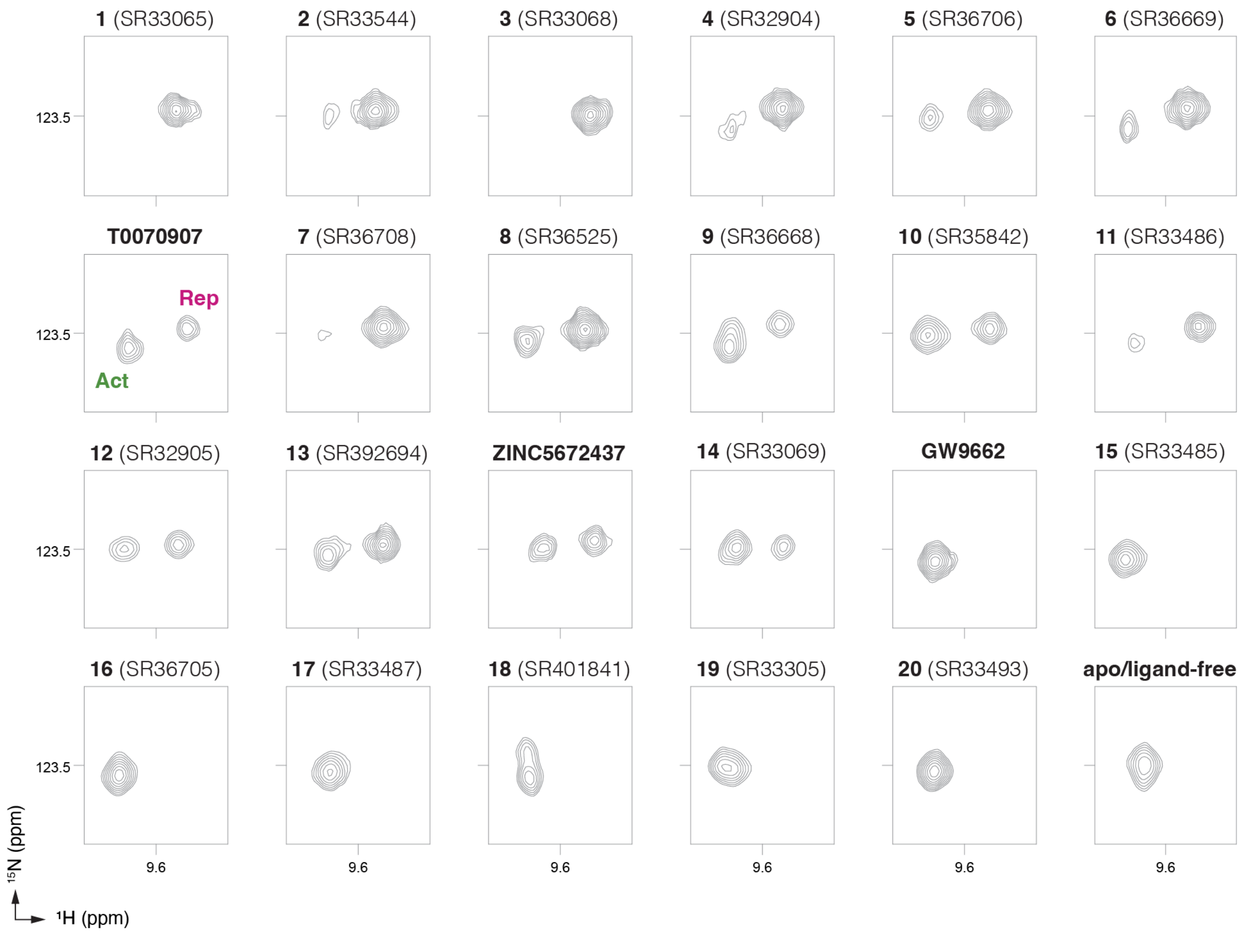


**Supplementary Figure S6.** 2D [$^{1}$H,$^{15}$N]-TROSY-HSQC NMR focused on Asp380 of $^{15}$N-labeled PPARγ LBD bound to compounds in the ligand series. The peak positions of the active-like (Act) and repressive-like (Rep) states populated by T0070907 are noted.

**Supplementary Table S1.** X-ray crystallography data collection and refinement statistics.

| | PPARγ LBD + GW9662 and NCoR-D2 | PPARγ LBD + ZINC5672437 and NCoR-D2 | PPARγ LBD + ZINC5672437 | PPARγ LBD + NCoR-D2 and SR33544 | PPARγ LBD + NCoR-D2 and SR33068 | PPARγ LBD + NCoR-D2 and SR32904 | PPARγ LBD + NCoR-D2 and SR36706 | PPARγ LBD + NCoR-D2 and SR33486 |
|---|---|---|---|---|---|---|---|---|
| **Data collection** | | | | | | | | |
| Space group | C $4_1$ $2_1$ $2_1$ | C $4_1$ $2_1$ $2_1$ | C 1 2 1 | P $4_1$ $2_1$ 2 | P $4_1$ $2_1$ 2 | P $4_1$ $2_1$ 2 | P $4_1$ $2_1$ 2 | P $4_1$ $2_1$ 2 |
| Cell dimensions | | | | | | | | |
| *a*, *b*, *c* (Å) | 61.72, 61.72, 163.92 | 61.05, 61.05, 159.29 | 91.63, 60.70, 117.08 | 62.054, 62.054, 160.719 | 62.125, 62.125, 162.209 | 61.888, 61.888, 164.288 | 61.765, 61.765, 161.08 | 61.968, 61.968, 163.608 |
| α, β, γ (°) | 90, 90, 90 | 90, 90, 90 | 90, 102.50, 90 | 90, 90, 90 | 90, 90, 90 | 90, 90, 90 | 90, 90, 90 | 90, 90, 90 |
| Resolution (Å) | 34.14-1.8 (1.864-1.8) | 37.95-1.8 (1.864-1.8) | 38.89-2.10 (2.175-2.10) | 43.88-1.42 (1.468-1.42) | 42.4-2.22 (2.302-2.22) | 41.07-2.02 (2.088-2.02) | 42.15-1.82 (1.89-1.82) | 30.98-2.12 (2.195-2.12) |
| $R_{merge}$ | 0.013 (0.156) | 0.020 (0.162) | 0.016 (0.246) | 0.03887 (1.065) | 0.1217 (1.479) | 0.07696 (1.073) | 0.05594 (0.4607) | 0.1486 (1.084) |
| $I / \sigma I$ | 28.00 (4.58) | 19.45 (3.89) | 16.87 (2.07) | 44.98 (2.64) | 13.50 (1.91) | 23.01 (2.28) | 39.82 (6.35) | 29.48 (4.01) |
| Completeness (%) | 99.93 (99.73) | 99.94 (99.96) | 97.43 (97.83) | 99.87 (98.90) | 98.61 (86.71) | 99.66 (97.86) | 99.83 (99.64) | 99.90 (100.00) |
| Redundancy | 2.0 (2.0) | 2.0 (2.0) | 2.0 (2.0) | 25.5 (20.8) | 12.4 (10.5) | 12.6 (12.0) | 25.1 (26.2) | 25.2 (26.4) |
| **Refinement** | | | | | | | | |
| Resolution (Å) | 1.80 | 1.80 | 2.10 | 1.42 | 2.222 | 2.02 | 1.825 | 2.12 |
| No. unique reflections | 30317 (2980) | 28829 (2799) | 35970 (3565) | 60517 (5857) | 16390 (1571) | 21902 (2109) | 28621 (2792) | 18917 (1851) |
| $R_{work}$ / $R_{free}$ | 19.8/22.6 | 20.2/23.4 | 24.4/28.8 | 18.53/19.75 | 21.66/25.14 | 19.43/24.91 | 18.08/21.64 | 18.52/21.83 |
| No. atoms | | | | | | | | |
| Protein | 2191 | 2230 | 4034 | 2190 | 2254 | 2258 | 2217 | 2214 |
| Ligand/ion | 18 | 21 | 64 | 24 | 32 | 27 | 41 | 70 |
| Water | 270 | 219 | 258 | 269 | 98 | 241 | 219 | 123 |
| *B*-factors | | | | | | | | |
| Protein | 28.13 | 26.11 | 29.94 | 30.09 | 50.82 | 41.18 | 33.62 | 44.94 |
| Ligand/ion | 25.67 | 22.64 | 36.65 | 29.45 | 46.72 | 40.05 | 37.15 | 51.17 |
| Water | 36.12 | 32.96 | 30.08 | 37.90 | 44.89 | 42.76 | 38.49 | 45.65 |
| R.m.s. deviations | | | | | | | | |
| Bond lengths (Å) | 0.009 | 0.008 | 0.011 | 0.011 | 0.021 | 0.002 | 0.010 | 0.020 |
| Bond angles (°) | 1.03 | 0.93 | 1.13 | 1.18 | 1.67 | 0.46 | 1.04 | 1.44 |
| Ramachandran favored (%) | 97.38 | 98.89 | 97.15 | 98.50 | 98.55 | 98.55 | 99.26 | 99.26 |
| Ramachandran outliers (%) | 0.37 | 0.00 | 0.00 | 0.00 | 0.00 | 0.00 | 0.00 | 0.00 |
| PDB accession code | 8FHE | 8FHG | 8FHF | 8FKC | 8FKD | 8FKE | 8FKF | 8FKG |

*Values in parentheses are for highest-resolution shell.

**Supplementary Table S2.** Density functional theory (DFT) quantum mechanical (QM) interaction free energies ($\Delta G_{bind}$) between compounds and PPARγ LBD residues comprising the pi-stacking aromatic triad residues (His323, His449, Tyr473) and others nearby residues (Cys285, Gln286, Tyr327, Met364, Lys367).

| **Compound number** | **$\Delta G_{BIND}$ (crystallized conformation)** | **$\Delta G_{BIND}$ (flipped $r_1$ ring conformation)** |
|---|---:|---:|
| **T0070907** | -15.5 | n.d. |
| 2 (SR33544) | -19.3 | n.d. |
| 3 (SR33068) | -17.4 | -16.9 |
| 4 (SR32904) | -17.0 | n.d. |
| 5 (SR36706) | -17.2 | n.d. |
| 11 (SR33486) | -21.3 | -27.5 |
